# Quantitative characterization of cell niches in spatial atlases

**DOI:** 10.1101/2024.02.21.581428

**Authors:** Sebastian Birk, Irene Bonafonte-Pardàs, Adib Miraki Feriz, Adam Boxall, Eneritz Agirre, Fani Memi, Anna Maguza, Rong Fan, Gonçalo Castelo-Branco, Fabian J. Theis, Omer Ali Bayraktar, Carlos Talavera-López, Mohammad Lotfollahi

**Affiliations:** Institute of AI for Health, Helmholtz Center Munich – German Research Center for Environmental Health, Neuherberg, Germany; School of Computation, Information and Technology, Technical University of Munich, Munich, Germany; University of Würzburg, Würzburg Institute of Systems Immunology (WüSI), 97074 Würzburg, Germany; Wellcome Trust Sanger Institute, Wellcome Genome Campus, Hinxton, Cambridge, UK; Institute of Computational Biology, Helmholtz Center Munich – German Research Center for Environmental Health, Neuherberg, Germany; Biomedical Center (BMC), Physiological Chemistry, Faculty of Medicine, Ludwig Maximilian University of Munich, Planegg-Martinsried, Germany; Laboratory of Molecular Neurobiology, Department of Medical Biochemistry and Biophysics, Karolinska Institutet, Stockholm, Sweden; University of Würzburg, Faculty of Medicine, 97074 Würzburg, Germany; Department of Biomedical Engineering, Yale University, New Haven, CT, USA; Yale Stem Cell Center and Yale Cancer Center, Yale School of Medicine, New Haven, CT, USA; Department of Pathology, Yale University School of Medicine, New Haven, CT, USA; Human and Translational Immunology Program, Yale School of Medicine, New Haven, CT, USA; Ming Wai Lau Centre for Reparative Medicine, Stockholm Node, Karolinska Institutet, Stockholm, Sweden; School of Life Sciences Weihenstephan, Technical University of Munich, Munich, Germany

## Abstract

Spatial omics allow us to identify and analyze communities of cells coordinating specific functions within a tissue. While these communities, defined as cell niches, are fundamentally shaped by interactions between spatially neighboring cells, we lack computational frameworks that can leverage spatial omics data to quantitatively characterize niches based on cell interaction events. To address this, we introduce NicheCompass, a graph deep learning method designed based on the principles of cellular communication. NicheCompass not only identifies cell niches, but also learns and informs about the signaling events shaping the identity of these niches. Unlike existing methods, it uniquely characterizes niches by quantifying their activity of spatial gene programs which represent diverse mechanisms of cell-cell communication and transcriptional regulation, thereby uncovering the underlying cellular processes constituting each niche. We showcase a comprehensive workflow encompassing data integration, niche identification, and functional interpretation, and demonstrate that, with its biologically informed design, NicheCompass outperforms existing methods. NicheCompass is broadly applicable to spatial transcriptomics data, which we illustrate by mapping the architecture of diverse tissues during mouse embryonic development, and delineating basal (*KRT14*) and luminal (*KRT8*) tumor niches in human breast cancer. We further introduce fine-tuning-based spatial reference mapping, revealing an SPP1+ macrophage-dominated tumor niche in non-small cell lung cancer patients. Additionally, we extend NicheCompass to multimodal spatial profiling of gene expression and chromatin accessibility, identifying and characterizing distinct white matter niches in the mouse brain. Finally, we apply NicheCompass to a whole mouse brain spatial atlas with 8.4 million cells demonstrating its scalability and ability to build foundational, interpretable spatial representations for entire organs. Overall, NicheCompass provides a novel approach to the challenge of identifying and analyzing niches, and suggests a more rigorous niche definition grounded in the quantitative characterization of underlying cellular processes.

## Introduction

Cell interactions play a key role in the formation of tissues, which comprise smaller, functionally diverse niches in different anatomical locations. Spatial signatures of the interaction activities of these niches can be found in their gene expression^1–3^, which can serve as a basis to identify these niches and analyze their function in healthy tissues and during development and disease. Identifying niches and quantifying their functions can help to unravel important insights into tissue architecture and spatial biomarkers, which can be harnessed to create novel diagnostic tools, identify new drug targets, and develop targeted therapies^4,5^.

Recent advances in spatial genomics promise to comprehensively resolve cell niches in tissues via imaging-based^6–9^ and sequencing-based^10–14^ spatial transcriptomics methods, as well as multi-omic profiling technologies^15^. Using these technologies, the community has constructed spatial atlases of whole organs across millions of cells and hundreds of tissue sections^16,17^. While spatial atlases can be used to reveal niches and cell interactions across multiple scales, there is a lack of computational approaches to quantitatively define niches from their biological functions while simultaneously informing about the mechanisms driving these such as cell interaction events. A quantitative definition can enhance our understanding of the physical and signaling cues that govern niches and allow them to adapt to homeostatic changes, and can facilitate the definition of higher-level tissue hierarchies.

Existing approaches define and identify cell niches by grouping cells from their tissue histology or from their gene expression similarity^18–31^. However, these approaches lack essential information about the underlying cellular processes, constraining the insights that can be obtained about niche function, and leading to the identification of niches that may be not biologically meaningful. Here, we present NicheCompass (Niche Identification based on Cellular Graph Embeddings of Communication Programs Aligned across Spatial Samples), the first approach that provides a quantitative characterization of cell niches. NicheCompass explicitly models cellular communication by predicting both the molecular profile of a cell and those of its neighbors as they relate to specific signaling events, all without any supervision. As a result, it can score each cell’s usage of different mechanisms of cellular communication in the context of their microenvironments, allowing for the identification and quantitative definition of niches based on underlying cell interactions. While existing methods can also address common tasks that have evolved in analyzing spatial omics data^32^, including the integration of multiple tissue samples to construct spatial atlases^20–26^, and downstream inference of cell-cell communication^33,34^, they differ from NicheCompass in at least two other analysis features and design choices: (1) they rely on single-cell data integration methods, which leads to suboptimal integration and inaccurate niche recovery^20,31^, (2) they do not scale to large datasets^18,25^, (3) they are limited to transcriptomic data and are unable to model spatial multi-omics data^18,21–23,25,31^, or (4) they are unable to map new spatial query data onto existing spatial reference atlases while retaining novel biological variation and removing technical variation to rapidly identify and characterize novel niches in the query^18,20–23,25,31^.

We demonstrate the utility and applicability of NicheCompass across diverse scenarios, including varying species, biological conditions, technologies, and modalities. In the process of mouse organogenesis, we reveal a highly resolved hierarchy of functional niches consistent across embryos and characterized by niche-specific GPs. In benchmarks across diverse datasets, we show that NicheCompass accurately recovers anatomical niches and removes batch effects in challenging integration scenarios. In human breast and lung cancer, we illustrate how NicheCompass aids in decoding the tumor microenvironment (TME) and capturing donor-specific spatial organization and cellular processes. In this context, we also highlight how NicheCompass enables spatial reference mapping to contextualize a query dataset with a large-scale reference, facilitating the identification of novel niches and contrasting underlying cellular processes. Additionally, we apply NicheCompass to a multimodal mouse brain dataset, demonstrating how multi-omics integration enhances niche resolution and provides more comprehensive niche characterization based on multimodal GPs. Finally, we demonstrate the scalability of NicheCompass by constructing a spatial atlas for the entire mouse brain across millions of cells.

## Results

### NicheCompass enables comprehensive quantitative analysis of spatial omics data

NicheCompass takes as input a spatial omics dataset with cell- or spot-level resolution and uses the spatial coordinates of cells/spots to construct a spatial neighborhood graph, where cells/spots constitute the nodes of the graph and edges indicate that cells/spots are spatial neighbors (**Fig. 1a** & **Methods**). Each node contains a feature vector composed of the cell’s/spot’s omics features (gene expression in unimodal scenarios and paired gene expression and chromatin accessibility peaks in multimodal scenarios). Additionally, each node is associated with potentially confounding biological or technical covariates whose effects we want to model (e.g. sample). To generate spatially and molecularly consistent latent representations of cells/spots, NicheCompass employs a graph neural network (GNN) encoder module with dynamic attention^35^ (**Fig. 1b** & **Methods**). This encoder jointly encodes the omics features of the subgraphs composed of nodes and their immediate neighbors, thus capturing cellular microenvironments. A separate encoding module generates embedding vectors for the node-specific associated covariates used for batch-effect removal^36^ (**Methods**). To make the latent cell/spot representations interpretable, NicheCompass incorporates domain knowledge of inter- and intracellular interaction pathways^37–41^ to form spatial prior GPs (**Methods**), each containing the set of genes involved in a specific pathway. The model learns spatially localized cellular activities of these GPs, with each latent feature being incentivized to learn the activity of a specific GP (**Fig. 1c**)^42^. To avoid limitations arising from the use of domain knowledge (e.g. quality issues, incompleteness, lack of niche-relevant factors such as global spatial gradients from morphogens^43^), NicheCompass can also learn the activity of spatial *de novo* GPs, represented as additional latent features (**Fig. 1c** & **Methods**). In contrast to prior GPs, *de novo* GPs do not consist of a predetermined set of genes; instead, they learn a set of genes that are not included in prior knowledge but have spatially co-localized expression.

**Fig. 1.**
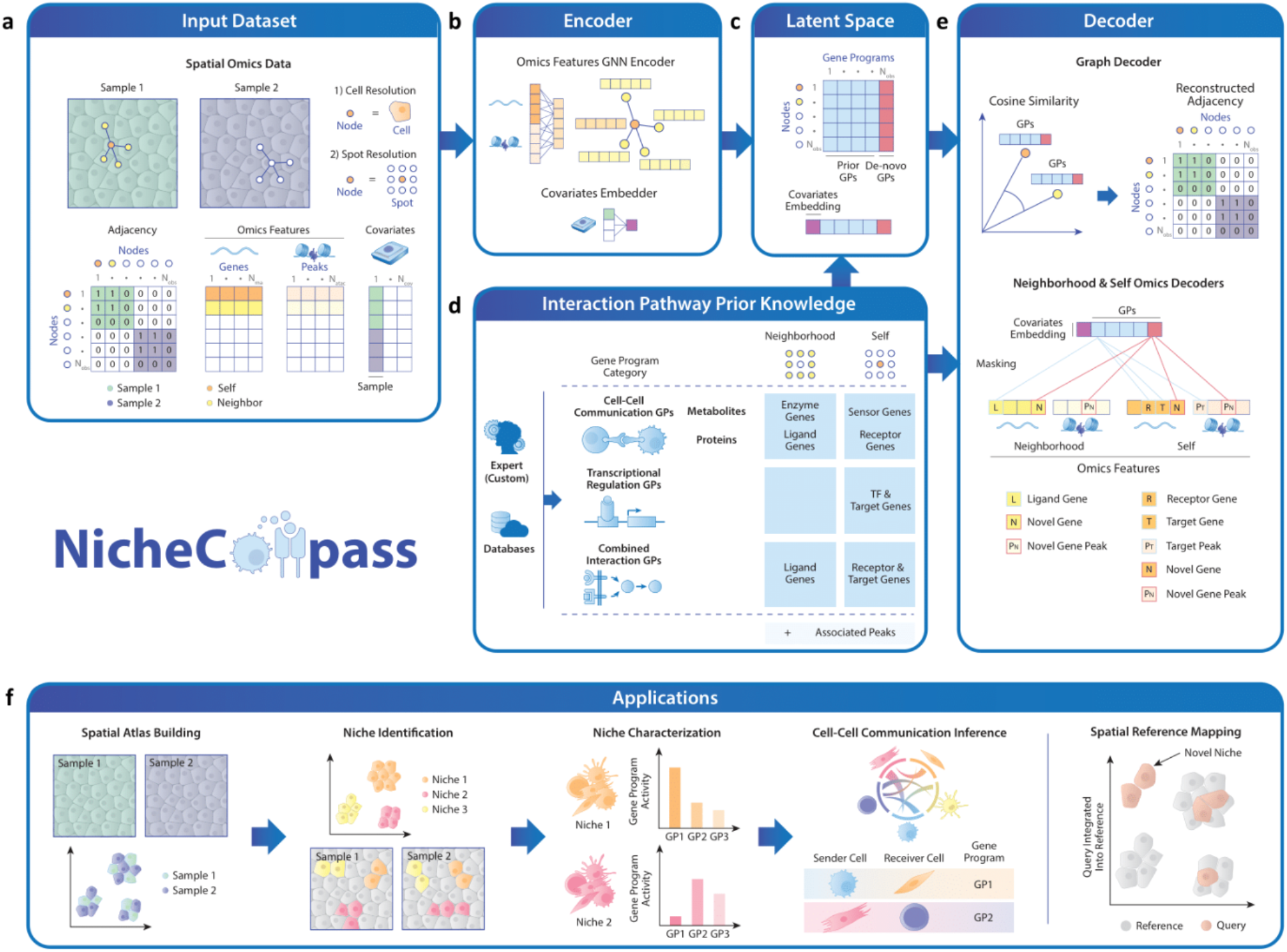
Overview of NicheCompass. **a,** NicheCompass takes as input a single- or multi-sample spatial omics dataset with cell- or spot-level observations, positioned in 2D space with x- and y-coordinates. A spatial neighborhood graph is constructed and represented with a binary adjacency matrix where each cell or spot represents a node, connected if spatially adjacent. Each observation consists of omics features and covariates that might introduce batch effects. Omics features contain either only gene expression counts or, in addition, matched chromatin accessibility peak counts. **b,** NicheCompass encodes the input omics features into a latent feature space using a graph neural network encoder. Covariates are embedded for batch effect removal. **c,** The model is incentivized to learn a latent feature space where each latent feature represents the spatially localized activity of an interaction pathway retrieved from domain knowledge, represented as a prior gene program (GP). In addition to prior GPs, the model can discover *de novo* GPs, which learn a set of spatially co-occurring genes and peaks during model training. **d,** Interaction pathways are extracted from databases or provided by an expert and fed to the model as GPs. GPs are classified into three categories and comprise a neighborhood- and self-component to reflect inter- and intracellular interactions. The neighborhood-component contains genes linked to the interaction source of intercellular interactions, and the self-component contains genes linked to the interaction target of intercellular interactions and genes linked to intracellular interactions. Peaks are associated with genes in the GPs if locationally proximal. **e,** Multiple decoder modules encourage the latent feature space to preserve spatial and molecular similarities while constraining each latent feature to represent the activity of a GP. A graph decoder reconstructs the input adjacency matrix. Omics decoders reconstruct a node’s gene expression and chromatin accessibility counts and aggregated counts of its neighborhood. The omics decoders are constrained to be linear and are masked based on GPs, thus enabling interpretability (exemplified by a combined interaction GP). **f,** NicheCompass facilitates critical downstream applications of spatial omics data analysis.

To model intercellular interactions between a cell/spot and its microenvironment, GPs consist of a self- and neighborhood-component (**Fig. 1d**). For prior GPs, the neighborhood-component includes pathway genes related to the source of intercellular interactions, reflecting that we model the microenvironment as signaling source; the self-component includes pathway genes related to the target of intercellular interactions or related to intracellular interactions, reflecting that we model a cell/spot as signaling receiver and responder. Depending on the nature of the underlying interactions, we categorize prior GPs into cell-cell communication GPs, transcriptional regulation GPs, and combined interaction GPs (**Fig. 1d**). Cell-cell communication GPs represent intercellular interactions and are divided into protein-mediated ligand-receptor GPs (**Supplementary Fig. 1a**) and metabolite-mediated metabolite-sensor GPs, where the expression of enzyme genes serves as a proxy for the presence of metabolites (**Supplementary Fig. 1b**). Transcriptional regulation GPs represent intracellular interactions, containing regulons comprising transcription factors and target genes (**Supplementary Fig. 1c**). Combined interaction GPs represent pathways of linked inter- and intracellular interactions, and include ligand and receptor genes as well as downstream targets regulated by ligand-receptor-induced signaling (**Supplementary Fig. 1d**). In multimodal scenarios, we link peaks to genes if they are located in the gene body or promoter regions (**Methods**)^44^. Peaks linked to genes in the neighborhood- or self-component of a GP are then added to the GP’s neighborhood- or self-component, respectively (**Fig. 1d**). We offer a default set of GPs for each category via built-in APIs to pathway databases^37–40,45^ (**Methods**); however, users can customize GPs to address specific biological questions.

The latent representations including prior and *de novo* GPs are decoded via multiple modules to reconstruct spatial and molecular information (**Fig. 1e** & **Methods**). First, a graph decoder module computes similarities between sample-specific pairwise latent node feature vectors to reconstruct the input spatial graph using an edge reconstruction loss. This encourages neighboring nodes to have similar representations, thus preserving spatial structure and prioritizing GPs with spatially localized activity. Second, masked omics decoder modules with linear layers reconstruct omics features. This enforces each latent feature to only reconstruct member genes of a specific GP, disentangling it from other sources of variation. Such a decoder design has been shown to have competitive performance while enabling interpretability through disentanglement in (dissociated) single-cell methods^42,46^. For our spatial GPs, we propose two different omics decoder modules that reconstruct the neighborhood- and self-component of GPs, respectively. The first module reconstructs the omics features of a node’s neighborhood, which are obtained by aggregating the input omics feature vectors across a node’s neighbors. The second module reconstructs the omics features of the node itself. Both these modules are masked so that only component-specific member genes and peaks of a GP are reconstructed; for example, a GP’s ligand gene is reconstructed in the node’s neighborhood and the corresponding receptor and target genes and associated peaks are reconstructed in the node itself. All omics decoders can access the covariate embeddings to remove undesired variation from the latent GP space^36^. In addition, to account for redundancy in prior GPs, we devise a mechanism to prioritize informative GPs and prune others during training (**Methods**). To encourage gene sparsity of target genes within GPs, we implement selective regularization (**Methods**).

Overall, the complete architecture of NicheCompass manifests as a multimodal conditional graph variational autoencoder^47,48^ (**Methods**). Our architectural design allows NicheCompass to quantitatively characterize tissue niches, and provides an end-to-end framework for spatial omics data analysis (**Fig. 1f** & **Supplementary Note 1**).

### NicheCompass elucidates tissue architecture across distinct mouse embryos

To demonstrate NicheCompass’ core capabilities, we applied it on a seqFISH^6^ mouse organogenesis dataset (**Methods**)^49^. This dataset consists of three spatially disparate embryo tissues extracted with varying positioning on a left-right axis (**Supplementary Fig. 2**), thus providing a challenging integration scenario. Integrating the embryo tissues with NicheCompass, we observed high cell type purity among neighbors in the inferred joint latent GP space (**Fig. 2a**). This is consistent with tissue architecture during mouse organogenesis being mainly characterized by colocation of homogenous cells forming spatially consistent niches^49^. We then clustered the latent GP space and, based on two characteristic cluster-enriched GPs (**Methods**), anatomical locations and cell type compositions (**Supplementary Fig. 3**), annotated the identified clusters with niche labels (**Fig. 2b**). The identified niches were spatially consistent but showed varying cell type patterns (**Fig. 2a**, **Fig. 2b** & **Supplementary Fig. 3**); this included niches consisting of homogeneous cell populations that closely matched cell type annotations (**Supplementary Fig. 4a**), niches comprising homogenous cell populations but at more fine-grained resolution compared to the annotated cell types (**Supplementary Fig. 4b**), and niches comprising heterogeneous cell populations (**Supplementary Fig. 4c**), emphasizing the importance of incorporating spatial information. While in the original study only one cluster was defined for the anterior central nervous system (CNS) (**Fig. 2a**), with NicheCompass the Forebrain, Midbrain and Hindbrain were clearly segregated into distinct niches (**Fig. 2b**). Moreover, with NicheCompass we were able to identify the Floor Plate as another CNS-associated niche. This niche was enriched in the *Shh* combined interaction GP, consistent with the ligand *Shh* being secreted from the floor plate^50^. Accordingly, we found specific expression of bona-fide floor plate marker genes in this niche (**Supplementary Fig. 5**). This illustrates the fine-grained niche identification performance and utility of our biologically informed, interpretable approach. In addition, niches successfully integrated across all embryo tissue sections (**Fig. 2c**), with all niches present in at least two embryos and most niches present in all three embryos (**Fig. 2d**). In cases where niches were not present in all three embryos, this was explained by very low embryo-specific abundance of cells of the major cell type constituting those niches (**Supplementary Fig. 6**).

**Fig. 2.**
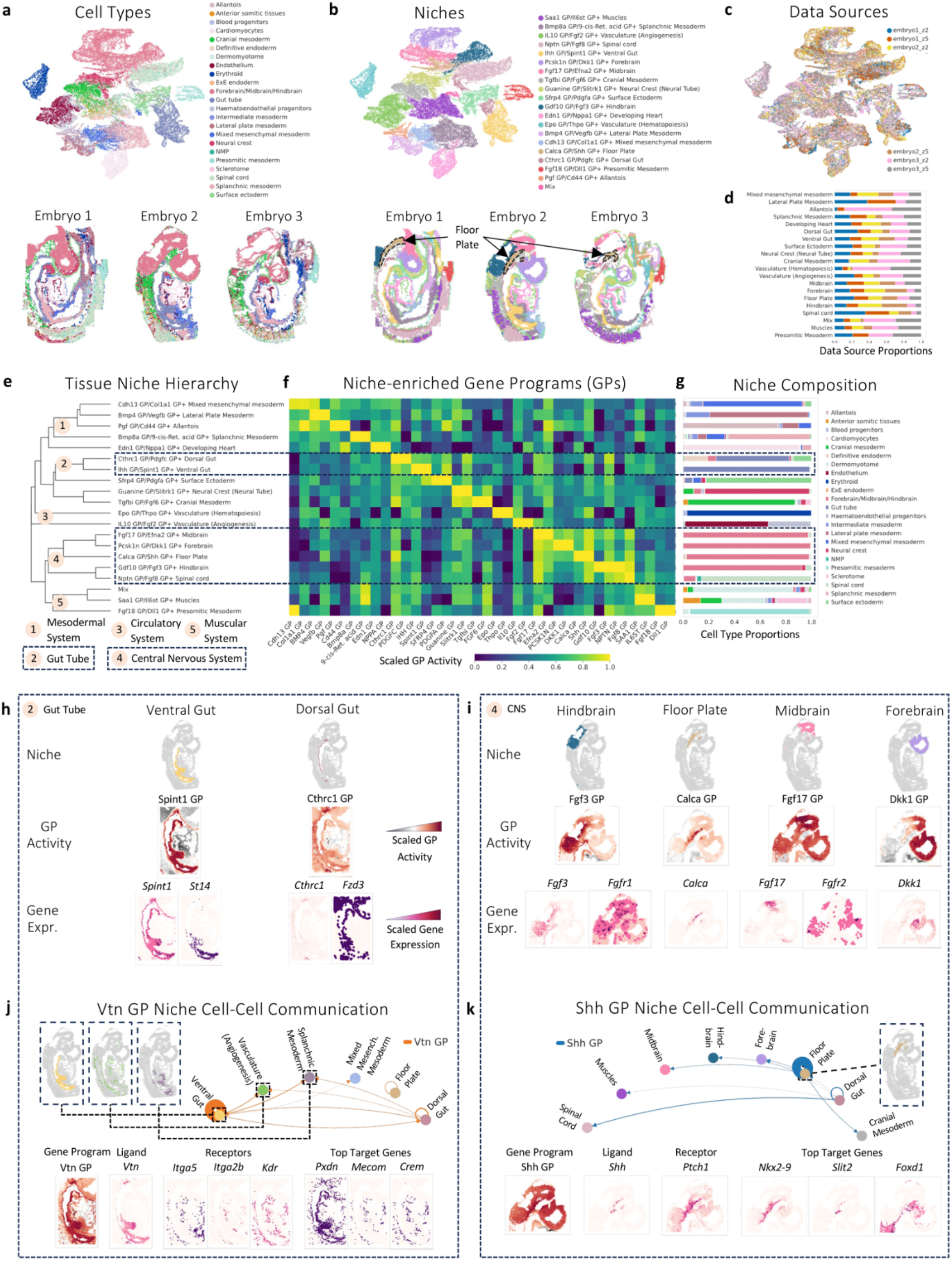
NicheCompass reveals cellular interactions driving tissue organization during mouse development. **a,** UMAP representation of the NicheCompass latent GP space after integrating three 8-12 somite stage embryo tissues and, below it, the embryo tissues, colored by originally annotated cell types/regions. **b,** Same as **a** but colored by the niches identified and annotated using NicheCompass. The nomenclature of niches is derived from two characterizing GPs and the region identity. **c,** The integrated latent GP space colored by data source (tissue sections), showing successful integration. **d,** The proportion of cells from each data source in each niche. Niche annotations are abbreviated. **e,** A dendrogram computed on average GP activities of the annotated niche labels shows a hierarchy of higher-order functional components. **f,** A heatmap containing for each niche from **e** the normalized activity of two characteristic GPs. A gradient along the clustering obtained from **e** is visible. **g,** Cell type proportions for each niche from **e**. **h,** GP activities of characteristic GPs differentiating the Ventral and Dorsal Gut niches from other niches. The gene expression of ligand and receptor genes of each GP correlates with the GP activity. **i,** Same as **h** but highlighting the four identified niches in the brain. CNS: Central Nervous System. **j,** Cell-cell communication analysis for one characteristic GP in the Ventral Gut niche. The circular plot shows the inferred communication strengths. Additionally, the gene expression levels of ligand, receptor and important target genes are consistent with the inferred main interacting niches. **k,** Same as **j** but highlighting one characteristic GP in the Floor Plate niche.

To investigate the preservation of global spatial organization by NicheCompass, we applied hierarchical clustering to group individual niches into higher-order functional components (**Fig. 2e** & **Methods**). This revealed a higher-level tissue organization consistent with anatomy, with the Midbrain, Forebrain, Floor Plate, Hindbrain and Spinal Cord niches constituting a Central Nervous System cluster, and the Dorsal and Ventral Gut niches forming a Gut Tube cluster (**Fig. 2e**). The activities of the characterizing GPs of each niche further supported this hierarchy and distinctly differentiated individual niches (**Fig. 2f**). This suggests that a few prior GPs prioritized by NicheCompass already explain a big part of tissue organization during mouse organogenesis. Additionally, niches that clustered into higher-order components were characterized by more similar cell type composition compared to niches distant in the cluster tree, highlighting meaningful incorporation of molecular information (**Fig. 2g**). All niches that comprised heterogeneous cell type populations had an idiosyncratic cell-type signature with distinct majority cell types compared to other niches, thus justifying their identification as molecularly distinct, separate niches. Overall, these results emphasized NicheCompass’ capability to unveil a hierarchy of biologically meaningful niches based on a latent feature space of spatial GPs.

We then analyzed GP activities in the identified gut and brain niches to characterize them based on underlying interaction mechanisms driving their niche identity. In each niche, we identified distinctly enriched activity of meaningful GPs (**Fig. 2h-i** & **Supplementary Note 2**). In the Ventral Gut niche, we found the highest activity of the Spint1 combined interaction GP (**Fig. 2h**). Based on gene importances (**Methods**), this GP was primarily driven by the ligand gene *Spint1*, encoding the inhibitor protein Hepatocyte growth factor activator inhibitor type 1 (HAI-1), and the receptor gene *St14*, encoding the tumor-associated type II membrane-anchored serine protease protein matriptase. It has previously been shown that an interaction between these proteins regulates the integrity of the intestinal epithelial barrier^51,52^. In the Dorsal Gut niche, we observed significant upregulation of the Cthrc1 combined interaction GP (**Fig. 2h**), mainly driven by the ligand gene *Cthrc1*. This GP was distinctly highly active in only a subregion of the Dorsal Gut niche, and, at this stage of embryo development, its ligand gene is known to be expressed explicitly in the notochord^53^, an embryonic midline structure that arises next to the dorsal gut and is critical for proper vertebrate development and foregut compartmentalization^54,55^. We therefore hypothesized that this GP marked the notochord as a further sub-niche, which we subsequently validated with highly spatially correlated expression of the notochord marker gene *Nog*^54^ (**Supplementary Fig. 7**). The GP also picked up the receptor gene *Fzd3*, albeit with much lower importance likely due to its wide-ranging expression in different tissue regions. Binding of Cthrc1 to Fzd proteins has been reported as a stimulator of the planar cell polarity pathway of Wnt signaling during the development of mice embryos^53^. In the Hindbrain niche, we identified upregulation of the Fgf3 combined interaction GP (**Fig. 2i**), primarily driven by the ligand gene *Fgf3* and its *Fgfr1* receptor^56^. It has been shown that Fgf3 signaling is active in hindbrain compartment boundaries which act as organizing centers to segregate adjacent areas into molecularly consistent domains, and is therefore required for establishment of correct segmental identity throughout the hindbrain and for neuronal development^57,58^. Among the enriched GPs in the Floor Plate niche, we identified the Calca combined interaction GP (**Fig. 2i**), which unambiguously demarcated this niche from other niches. Its GP activity was driven by the ligand gene *Calca,* a gene encoding peptide hormones such as calcitonin, important in a subpopulation of glutamatergic neurons at the midbrain–hindbrain junction implicated in various homeostatic functions^59^. In the Midbrain niche, we identified enriched activity of the Fgf17 combined interaction GP (**Fig. 2i**), driven by the ligand gene *Fgf17*, which plays a key role in the regulation of early vertebrate midbrain patterning^60^, and the *Fgfr2* receptor gene with expression in the surrounding brain niches. The resulting Fgf/Fgfr signaling pathway is known to be critically involved in a multitude of processes during embryonic development^61^. In the Forebrain niche, we identified the Dkk1 ligand-receptor GP (**Fig. 2i**), which was solely driven by the ligand *Dkk1* and whose activity distinctly separated the niche from other brain niches. Dkk1 promotes the formation of precursors of forebrain neurons in mice and a loss of Dkk1 results in anterior truncation^62,63^. Lastly, to validate the integrity of the learned GP activities, we compared the expression of ligand and receptor genes with their reconstructed expression by the model, and observed ample congruence, indicating an excellent preservation of molecular features in the NicheCompass latent GP space (**Supplementary Fig. 8** & **9**).

Next, we used the inferred GP activities to analyze underlying cell-cell interactions. For this, we devised a strategy to compute source- and target-specific cell-cell communication potential scores for each cell (**Methods**) to compute communication strengths between cell pairs (**Methods**), aggregated at niche- and cell-type-level. Using this strategy, we analyzed the Vtn combined interaction GP, which was enriched in the Ventral Gut niche (**Fig. 2j** & **Supplementary Fig. 10**) and extracted known interactions of the extracellular matrix (ECM) protein Vitronectin (Vtn) with the Kdr receptor and the integrin receptors encoded by *Itga5* and *Itga2b*. These signaling interactions represent main regulatory patterns of cellular responses during development; for instance, screening by an experimental assay has previously identified Vitronectin as a main ECM protein component that promoted differentiation of embryonic stem cells into definitive endoderm, an early embryonic cell population fated to give rise to internal organs such as the gut^64^. Beyond the ligand-receptor interactions, important genes of this GP also included target genes (*Pxdn, Mecom, Crem*) that showed spatially correlated expression patterns (**Fig. 2j**). Based on the computed communication strengths, the GP was characterized not only by intra-niche interactions in the Ventral Gut niche but also by interaction across niches, in particular from the Ventral Gut niche to the neighboring Vasculature (Angiogenesis) and Splanchnic Mesoderm niches. This finding was consistent with Vitronectin-integrin signaling being a crucial causal contributor to mouse angiogenesis^65^. Moreover, the depletion of *Mecom*, an endothelial cell lineage regulator and one of the target genes, has further been shown to impair Zebrafish angiogenesis^66^. Following the same strategy, we also interrogated the *Shh* combined interaction GP, enriched in the Floor Plate niche (**Fig. 2k** & **Supplementary Fig. 11**). Alongside the ligand gene *Shh* and the receptor gene *Ptch1*, NicheCompass identified various genes regulated downstream of Shh signaling as important genes; these include *Nkx2-9*, which has a possible role in the specification of dopaminergic neurons^67–69^, *Slit2*, thought to act as a matrix guiding or supporting the migration of the ventral nerve cord axons^70^, and *Foxd1*, a known *Shh* target in retina patterning^71^ and necessary component for the proliferative response of immortalized mouse embryonic fibroblasts to *Shh* ligand stimulation^72^. While we observed Shh GP activity mainly in the Floor Plate niche, it also extended to other brain niches, driven by receptor and target gene expression. Despite *Shh* not being experimentally detected in these niches, this is in accordance with Shh signaling occurring more broadly in the brain^73^.

Overall, these results demonstrate how NicheCompass can jointly infer niches and their underlying cell interaction mechanisms in the important biological process of development, highlighting NicheCompass’ unique capabilities ranging from biologically meaningful niche identification across multiple unaligned tissues to GP-based niche characterization and cell-level cell-cell communication inference.

### NicheCompass accurately identifies niches across diverse technologies, species and tissues

To assess NicheCompass’ niche identification performance, we benchmarked it against other spatial embedding methods^18,20,25,34^ across diverse datasets, and compared the obtained cellular representations and niche labels. In the first phase of this process, we applied all methods on a SlideSeqV2 dataset of the mouse hippocampus (**Methods**), whose spatial tissue organization closely mirrors anatomical structure^10^. Initially, we qualitatively compared the identified niches of NicheCompass with an anatomical map from the Allen Mouse Brain Reference Atlas^74^ (Mouse, P56, Coronal, Image 71; **Fig. 3a**). We observed close conformity between niches and anatomical subcomponents, leading to accurate recovery of local tissue structure. To assess whether our model also captured global tissue structure, we derived a tissue niche hierarchy through hierarchical clustering of the latent GP space (**Methods**). We compared this hierarchy with the anatomical taxonomy of the Allen Mouse Brain Reference Atlas^74^, and found that subcomponents clustered into higher level functional components (**Fig. 3b** & **Supplementary Fig. 12a**). Of the obtained functional components, the Isocortex and Hippocampus formed a cluster tree entirely consistent with anatomical taxonomy; contrarily, the Thalamus cluster tree showed slight deviations, illustrated by the separation of the MH and LH niches into different branches and close integration of the MH and V3 niches instead. However, this deviation was reasonably explained by differential cell type composition: the MH and V3 niches were both characterized by abundant Ependymal cells not present in the LH niche, whose cell type composition was very similar to its neighboring LP niche, marked by high cell type entropy with an increased number of astrocytes and oligodendrocytes. Close resemblance of cell type composition for niches close in latent GP space was, in fact, generally observable and aligns with our objective to learn latent representations that reflect spatial and molecular similarities. Subsequently, we compared our identified niches with those from similar methods (**Fig. 3c, Methods** & **Supplementary Note 3**). One of the methods, STACI^25^, failed to train successfully due to memory overflow on our 40GB GPU. Among methods that trained successfully, only NicheCompass could identify four separable cortical layer niches and preserve their spatial adjacency in the latent space to form a high-level Isocortex cluster (**Fig. 3c**). In contrast, CellCharter could not clearly separate the cortical layer 5 and while DeepLinc could retrieve three separable cortical layer niches, these did not cluster into a joint functional component composing the Isocortex (**Supplementary Fig. 12b**). Moreover, NicheCompass showed superior spatial preservation and purity of niches whereas other methods produced some spatially scattered niches that did not represent a defined anatomical subcomponent (**Fig. 3c**).

**Fig. 3.**
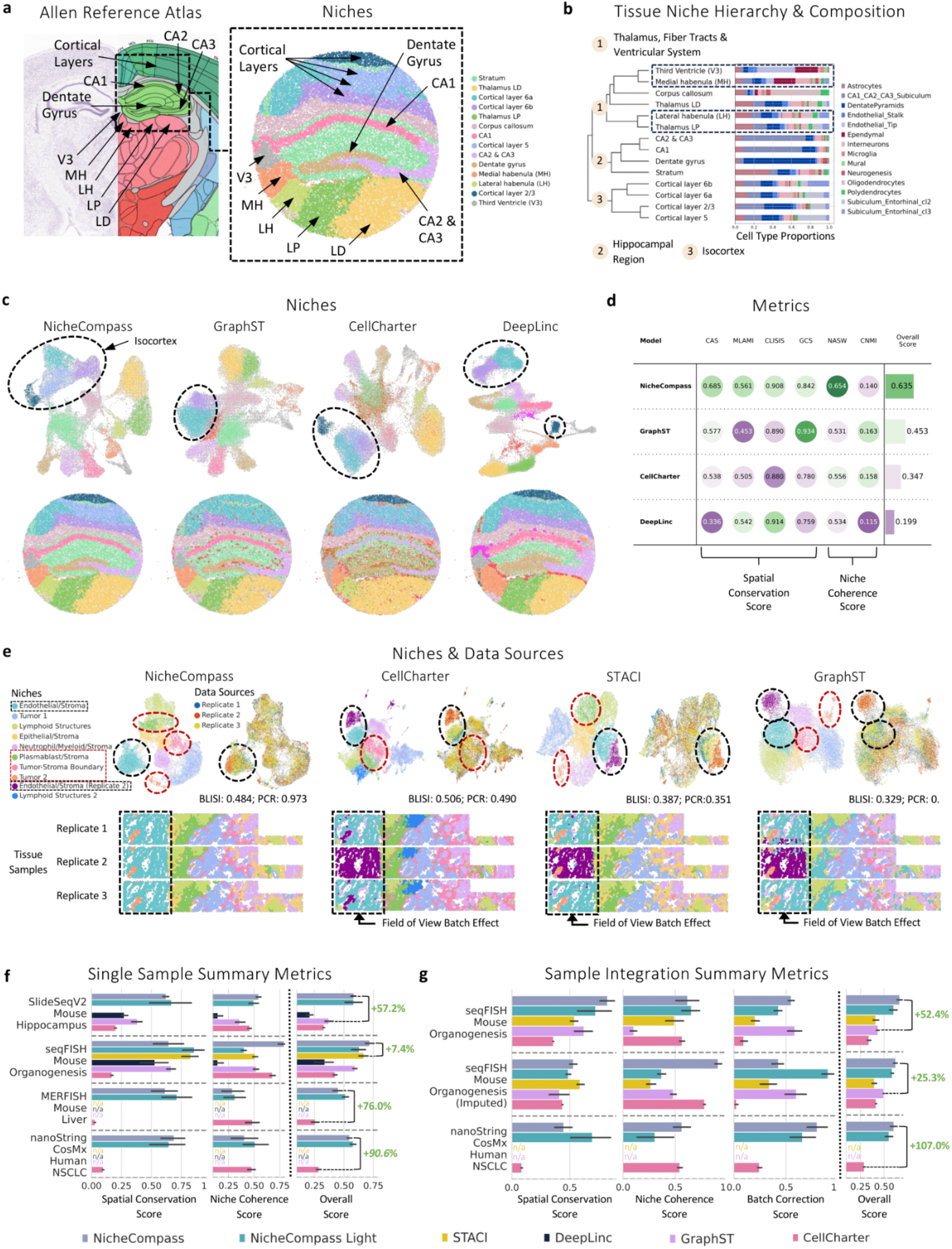
Benchmarking NicheCompass across various scenarios. **a,** A coronal mouse brain image from the Allen Reference Atlas showing anatomical subcomponents in the isocortex, thalamus, and fiber tracts. Next to it, a mouse hippocampus tissue, sequenced with the SlideSeqV2 technology, highlighting corresponding niches obtained with NicheCompass. **b,** A dendrogram computed on average GP activities, uncovering a hierarchy of anatomically and molecularly similar niches. **c,** UMAP representations of the latent feature spaces from NicheCompass and three competing methods, and the mouse hippocampus tissue below them, colored by niches identified through clustering of the latent feature spaces. The number and color of clusters match with **a**. **d**, Six metrics across the two categories spatial conservation and niche coherence display the performance of each method on this dataset. The overall score is computed by normalizing and aggregating the individual metrics into categories, followed by equal weighting of categories. **e,** A comparison of the integration performance of NicheCompass, CellCharter, STACI, and GraphST on a subsample of three replicates from the NanoString CosMx human NSCLC dataset. Illustrated are the UMAP representations of the latent feature spaces of the models and below them the tissue, colored by annotated niches. Niches in the first field of view and additional niches only jointly identified by NicheCompass are highlighted, showing differences in batch effect removal capabilities and obtained resolutions. UMAP representations colored by data source further emphasize differences in batch-effect removal. **f, g,** Performance summary metrics of NicheCompass and similar methods on four single-sample (**f**) and three multi-sample (**g**) datasets. The bars display the mean across eight training runs for each method and dataset with varying numbers of neighbors. The error bars display the 95% confidence interval. A percentage change accentuates the difference between the overall score of NicheCompass and the second-best method. Missing bars reflect unsuccessful model training due to memory constraints.

Next, we quantified the quality of each method’s cellular representations and niche labels using six metrics, encompassing two categories: spatial conservation and niche coherence (**Methods**). We combined these metrics into a normalized overall score (**Methods**) to provide a measure that accurately balances spatial coherence and biological meaningfulness (**Supplementary Note 4**). Consistent with our qualitative assessment, NicheCompass outperformed other methods (**Fig. 3d**). GraphST^18^, previously shown to outperform similar methods^19^ on the same dataset, ranked second. While other methods showed selectively superior performance on individual metrics, NicheCompass ranked first in both categories with reliably high performance across metrics, displaying high robustness to metrics selection. To enable a fair comparison with STACI, which did not successfully run on the full dataset, we further performed benchmarking on a 25% subsample (**Methods** & **Supplementary Fig. 13a**). The results were more similar between methods than the ones from the full dataset, with the major difference limited to the separation of the CA1 niche from the Stratum niche. NicheCompass and GraphST were the only methods to achieve an accurate identification of parts of the Stratum niche between the CA1 and Corpus Callosum niches, as reflected in a higher niche coherence score (**Supplementary Fig. 13b**). In a direct comparison, niches identified by GraphST showed lower spatial consistency with more mixed niche assignments of adjacent cells, resulting in lower spatial conservation and overall scores.

In the second phase of the benchmarking process, we aimed to compare NicheCompass’ integration capability with that of CellCharter, GraphST, and STACI, whose authors highlighted their ability to jointly analyze multiple tissue slices while removing batch effects. For this purpose, we selected a NanoString CosMx human non-small cell lung cancer dataset (NSCLC, **Methods**). As it was not feasible to run GraphST and STACI on the full dataset due to memory overflow, we used a 10% subsample that exhibited strong batch effects in the first field of view of the second replicate (**Methods** & **Supplementary Fig. 14a**). NicheCompass was the only method that successfully integrated the first field of view across all three replicates (**Fig. 3e**). In addition, it was able to identify a Tumor niche surrounded by endothelial stroma, a Tumor-Stroma-Boundary niche, which is known to be a critical target for cancer therapy^75^, and could differentiate between Stroma enriched by endothelial cells and Stroma enriched by plasmablast cells (**Supplementary Fig. 14b**). While CellCharter and STACI also achieved the latter, CellCharter could not identify the Tumor niche, and STACI failed to identify the Tumor-Stroma-Boundary niche. GraphST, on the other hand, was able to recognize the Tumor and Tumor-Stroma-Boundary niches but grouped the Plasmablast-enriched Stroma into the Endothelial Stroma niche. Analyzing cell type composition of these niches across the four methods further highlighted improved functional meaningfulness of NicheCompass niches, which were again characterized by an idiosyncratic cell-type signature, dominated by different cell types, namely plasmablast cells in the Plasmablast-enriched Stroma, fibroblasts in the Tumor-Stroma-Boundary and endothelial cells in the Endothelial-enriched Stroma (**Supplementary Fig. 14b**). Besides, we evaluated multiple method variants, including a version of our model that did not include the field of view as covariate (**Supplementary Fig. 14c**). While NicheCompass remained the only method to successfully remove field of view effects, excluding the field of view as covariate led to a breakdown of integration, attributing our superior performance to the architectural design of covariate embeddings. We also quantified the batch integration performance with two batch correction metrics previously proposed in a single-cell benchmarking paper (**Methods**)^76^. These metrics reinforced advantages in NicheCompass’ integration performance, while other metrics highlighted competitiveness regarding spatial conservation and niche coherence (**Fig. 3e** & **Supplementary Fig. 14d**).

To assess the methods’ general applicability and scalability, we compared them across datasets of different sizes and gene coverage, generated with different technologies, including next-generation sequencing-based and imaging-based protocols (**Fig. 3f, Supplementary Fig. 15a** & **Methods**). NicheCompass and CellCharter were the only methods that ran successfully on the two bigger datasets (77,391 and 367,335 cells, respectively) due to memory constraints, and NicheCompass outperformed other methods on all datasets in terms of the overall score (+57.2%, +7.4%, +76.0%, and +90.6% compared to the second-best method on the four datasets, respectively) and in most categories. In addition, NicheCompass’ superior performance was robust against data subsampling (**Methods**) and maintained in small-scale data regimes, with STACI being the only competitive method (**Supplementary Fig. 15b**). We further benchmarked NicheCompass, CellCharter, GraphST and STACI, the methods amenable to sample integration, across three datasets with multiple samples (**Fig. 3g**, **Supplementary Fig. 16** & **Methods**). Again, we found that NicheCompass considerably outperformed other methods in terms of the overall score (+52.4%, +25.3%, and +107.0% compared to the second-best method on the three datasets, respectively). While GraphST achieved better batch correction scores on one dataset, this came at a significant cost concerning other metric categories due to excessive integration. Overall, across both single- and multi-sample benchmarks, NicheCompass demonstrated its ability to handle large volumes of data efficiently. While other methods failed due to memory constraints or were not performant when dealing with larger subsets of data, NicheCompass seamlessly scaled due to its efficient use of memory achieved through mini batch loaders with inductive neighbor sampling and sparse internal data representations (**Methods**, **Supplementary Fig. 17** & **Supplementary Note 5**).

### NicheCompass learns *de novo* gene programs to delineate basal and luminal niches in breast cancer

We investigated the utility of NicheCompass in discovering *de novo* GPs using imaging-based Xenium human breast cancer data^77^ (**Methods**). This dataset was created with a gene panel consisting of cell type markers, with a limited number of genes in our prior knowledge databases (23% of genes in the dataset). Consequently, it presented an interesting scenario to investigate whether NicheCompass could effectively identify niches when it primarily has to rely on *de novo* GPs.

NicheCompass could integrate multiple replicates from the same tissue (**Fig. 4a-d**). Using the 313 probes from the data, we identified and annotated 11 cell types and 27 cell states (**Fig. 4a**, **Supplementary Fig. 18** & **Methods**). Further, clustering the NicheCompass latent GP space, we identified 14 niches (**Fig. 4b**) that displayed specific anatomical localizations, providing a detailed overview of the tissue architecture of the resected tumor samples (**Fig. 4e**). Because the probes in this experiment were limited, it was not possible to assign niches a functional annotation as previously. Instead, we annotated the niches based on the two most abundant cell types in each cluster (**Supplementary Fig. 19**). Niches were rich in Immune, Epithelial and Epithelial-Mesenchymal-Transition (EMT) cellular states; Epithelial-Fibroblast (Epi-FB), CD4+T cells (CD4+T), and EMT-Immune niches were the biggest (26.9%, 24.9%, and 18.6% of all cells, respectively).

**Fig. 4.**
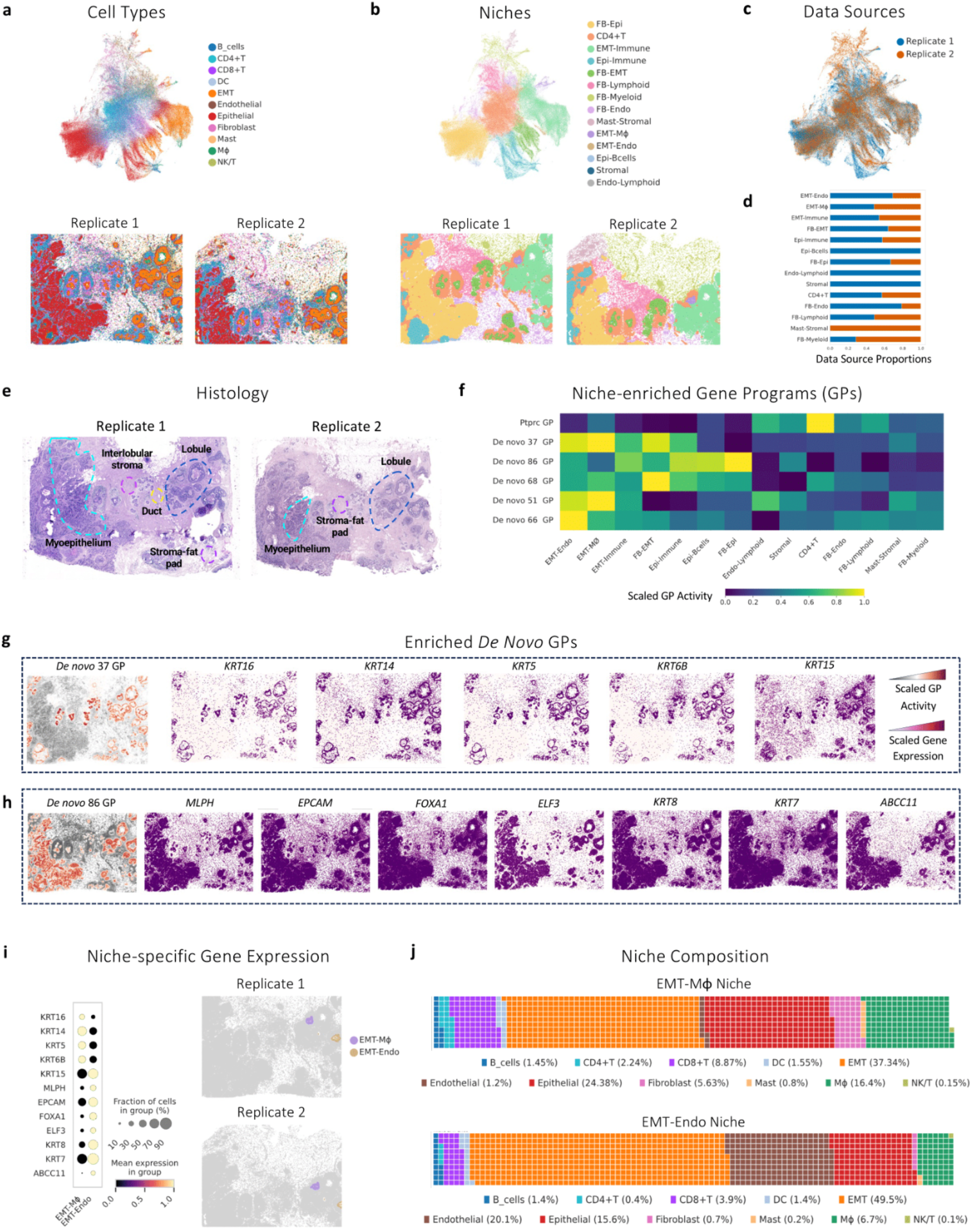
NicheCompass identifies meaningful niches and *de novo* gene programs in human breast cancer. **a,** UMAP representation of the NicheCompass latent GP space after integrating two replicate tissues of a 313 probe 10X Genomics Xenium dataset, and below it the tissue replicates, colored by cell types. **b,** Same as **a** but colored by the niches identified and annotated using NicheCompass. Niches are named as follows: FB-Epi (Fibroblast-Epithelial), CD4+T (CD4+ T cells), EMT-Immune, Epi-Immune (Epithelial-Immune), FB-EMT (Fibroblast-EMT), FB-Lymphoid (Fibroblast-Lymphoid), FB-Myeloid (Fibroblast-Myeloid), FB-Endo (Fibroblast-Endothelial), Mast-Stromal (Mast Cells-Stromal), EMT-Mϕ (EMT-Macrophage), EMT-Endo (EMT-Endothelial), Epi-Bcells (Epithelial-B cells), Stromal, Endo-Lymphoid (Endothelial - Lymphoid). **c,** The integrated latent GP space colored by data source, displaying successful integration. **d,** The proportion of cells from each data source in each niche. **e,** Annotated Hematoxylin and Eosin (H&E) slides of the breast cancer tumor resection. **f**, A heatmap of the normalized activity of characterizing *de novo* GPs associated with cancer progression and pathological histology. **g**, GP activity and expression of important genes of *de novo* 37 GP, showing high correlation between GP activity and member gene expression. **h**, Same as **g** but for *de novo* 86 GP. **i,** Niche-specific normalized expression of important member genes of de novo 37 and de novo 86 GPs, showing a clear separation between the two niches based on KRT14 (basal breast tumor cells) and KRT8 (luminal breast tumor cells) member gene transcriptional activities. The dot size represents the fractions of cells in a niche with expression higher than 0, while the dot color represents the mean expression level within the expressing cells, normalized across the two niches. **j**, Cellular composition of the EMT-Macrophage and EMT-Endo niches.

The number and nature of probes in the dataset made inference of cell-cell communication and transcriptional regulation difficult and unreliable; however, despite the inherent limitations in the dataset, NicheCompass demonstrated remarkable efficacy in identifying niche-specific GPs, crucial for understanding the intricate cellular tumor microenvironments. We identified the Ptprc combined interaction GP as enriched in the CD4+T niche, which has been described as a biomarker for cancer prognosis^78^ and as an interaction marker between breast cancer stem cells (BCSCs) and CD4+ T cells^79^ (**Fig. 4f**). In addition, various enriched *de novo* GPs picked up highly correlated genes, among them two *de novo* GPs with increased activity across immune and Epithelial-Mesenchymal transition (EMT)-associated niches with potential as pathology biomarkers and drug targets (**Fig. 4f** & **Supplementary Fig. 20**).

One novel GP identified by NicheCompass, *de novo* 37 GP (**Fig. 4g** & **Supplementary Fig. 21a**), comprised genes *KRT16*, *KRT14*, *KRT5*, *KRT6B*, and *KRT15*. *KRT14* and *KRT16* frequently exhibit co-expression and serve as basal markers. *KRT16* has been reported to be a metastasis-associated protein, promoting EMT and cellular motility^80^. The association of *KRT6B* with basal-like breast cancer and the correlation of *KRT15* overexpression with tumor metastasis in breast cancer patients highlight the critical role these markers play in oncological studies^81^. Moreover, *KRT16* has been identified as a target of the *ATF4* transcription factor, integral to the adaptive integrated stress response^82^, which has been described in HCT116 and HeLa cells.

Another novel GP, *de novo* 86 GP (**Fig. 4h** & **Supplementary Fig. 21b**), included genes such as *MLPH*, *EPCAM*, *FOXA1*, *ELF3*, and *KRT8*, each contributing uniquely to breast cancer pathology. *ELF3*, a transcription factor, activates *KRT8* and is implicated in epithelial cell differentiation and tumorigenesis^83,84^. The interaction between *ELF3* and *FOXA1* in endocrine-resistant ER+ breast cancer cells, evidenced by the enrichment of *ELF3* motifs at FOXA1/H3K27ac-gained enhancers, exemplifies the intricate transcriptional regulation mechanisms in cancer.

Additionally, NicheCompass was able to adequately delineate niches anatomically, identifying *de novo* GPs with genes previously associated with the pathology of histological structures (**Fig 4i**). For instance, *de novo* 37 GP and *de novo* 86 GP highlighted a transcriptional signature for *KRT14*+ proliferative epithelial tumor cells that cohabit with myeloid cells^85^, and an epithelial-vascular niche driven by *EPCAM* and *KRT8* genes, associated with preneoplastic and luminal tumor progression. Previous studies have described *KRT14* as a biomarker for basal breast cancer cells, and *KRT8* as a biomarker for luminal breast cancer cells^86^, in line with our results. The corresponding GPs were active in the EMT-Mϕ (Epithelial-Mesenchymal-Transition and Macrophages) and EMT-Endo (Epithelial-Mesenchymal-Transition and Endothelial cells) niches, respectively (**Fig. 4f-j**).

Despite the constraints of the Xenium dataset, NicheCompass correctly identified GPs relevant to specific niches. This use case highlights the capacity of NicheCompass to identify niches through meaningful GPs, even with limited data, allowing for the exploratory analysis of publicly available datasets from different experimental designs or chemistry versions.

### NicheCompass constructs a spatial atlas of heterogeneous non-small cell lung cancer patients

To identify and characterize the complex physical and molecular interactions between cancer cells and infiltrating cells, and to highlight patient-specific tumor microenvironment (TME) features, we applied NicheCompass to the previously described NSCLC dataset (**Methods**)^8^, now using all samples. The complete dataset consists of eight tissue sections, with five different donors, two of which have two and three replicates, respectively. The selected panel of 960 gene probes captures key aspects of tumor biology, including cell identity and state, intracellular signaling, and cell-cell interactions. Consequently, this dataset was well suited for a spatial reference mapping scenario, prompting us to test NicheCompass’ utility in such a scenario, and validating whether it could identify donor-specific niches and their underlying cellular mechanisms.

First, we built a NSCLC reference atlas using four donors and two replicates. Clustering of the NicheCompass latent GP space identified 12 niches (**Fig. 5a**), driven by differential cell composition (**Fig. 5b** & **Supplementary Fig. 22c**), spatial organization (**Supplementary Fig. 22e**), and gene expression (**Supplementary Fig. 22f**). In line with the idiosyncratic expression patterns characteristic of tumor cells, the majority (92%) of cancer cells formed tumor-exclusive niches (defined as more than 75% of tumor cells), with only highly infiltrative stromal niches containing a significant number of tumor cells (e.g. niche 6 – Infiltrating neutrophil expansion, **Supplementary Fig. 22c**). Consistent with inter-tumoral heterogeneity, tumor niches were donor-specific but shared across technical replicates, demonstrating that this was not driven by technical batch effects (**Fig. 5c** & **Supplementary Fig. 22d**). While a strong donor-dependency was also observed in stroma niches, shared niches were identified when resembling structures existed (**Fig. 5c** & **Supplementary Fig. 22d**). This is in line with the finding that patients with NSCLC can be stratified based on the infiltration patterns of their TME^87^. Hierarchical clustering of the identified niches based on average GP activities correctly separated tumor from stromal niches, and divided stroma niches into neutrophil-dominated stroma, immune-deserted stroma, immune-rich stroma, and immune expansions (**Supplementary Fig. 22a**), showing that cells from different donors are correctly mapped at the global level, even in a context of high inter-sample heterogeneity.

**Fig. 5.**
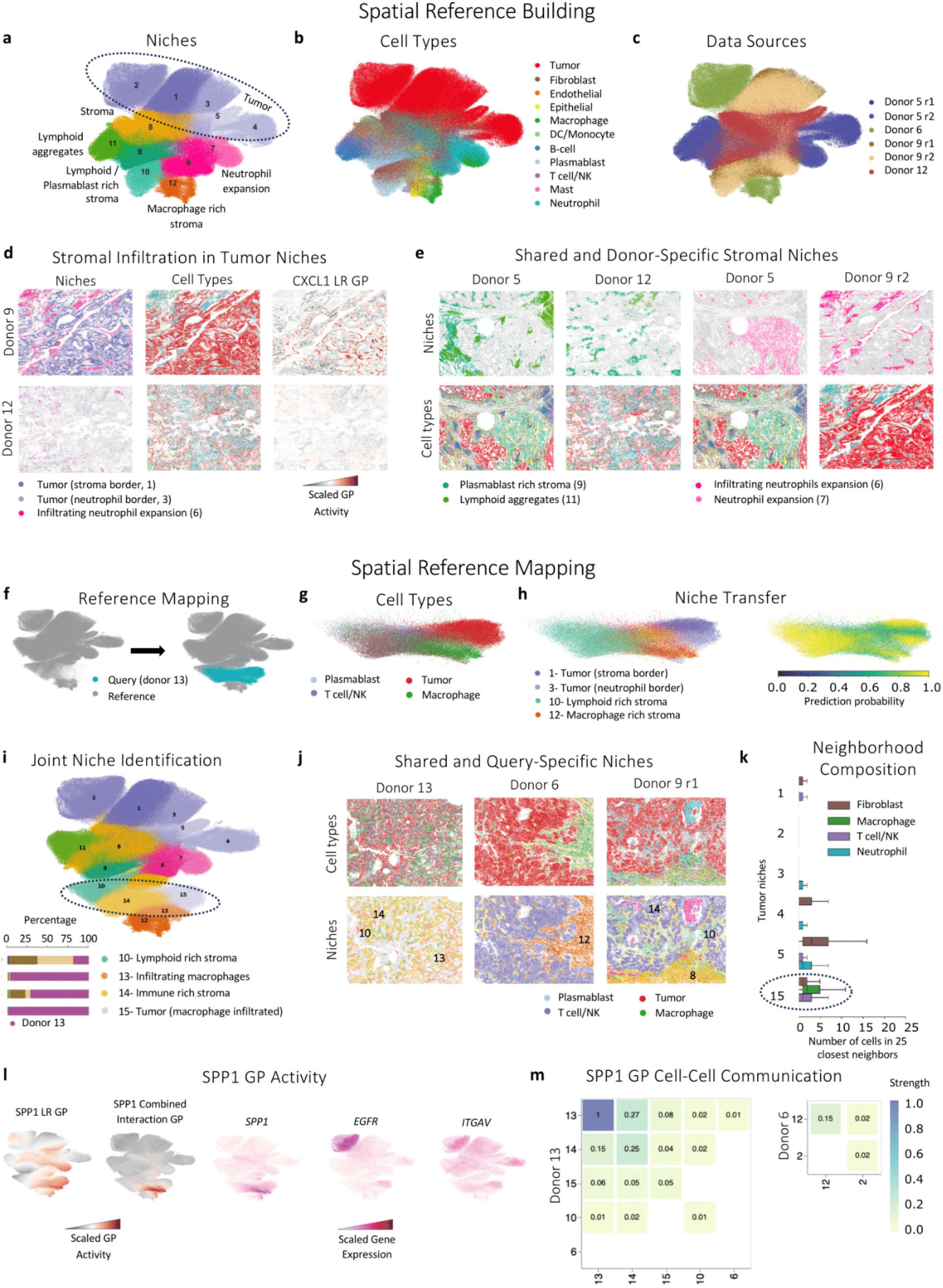
NicheCompass enables spatial reference mapping to contextualize new donors with a spatial atlas, revealing emergent niches. **a-c,** UMAP representations of the NicheCompass latent GP space after integrating six lung samples from NSCLC patients, colored by identified and annotated niches using NicheCompass (**a**), pre-annotated cell types (**b**), and sample donor and replicate (**c**). **d**, Spatial visualization of donor 9 and 12 tissue sections, colored by identified niche, cell type and the CXCL1 ligand-receptor GP activity, for comparing tumor niches interacting with stromal tissue (1) or neutrophils (3). **e**, Spatial visualization of tissue sections colored by niche and cell type, highlighting shared and donor-specific stromal structures identified across donors. **f**, UMAP representation of the NicheCompass spatial reference and the query cells mapped via fine-tuning. **g**, UMAP representation of the mapped query cells, colored by pre-annotated cell types. **h**, same as **g** but colored by niche label as predicted by a k-nearest neighbors classifier trained on the reference. Next to it, the prediction probability of the classifier. **i**, UMAP representation of the joint reference and query NicheCompass latent GP space, colored by niche as identified by re-clustering. In addition, barplots representing the donor distribution of the niches the query sample maps to. **j**, Spatial visualization of the query tissue (donor 13), and the most similar samples from the reference, colored by cell type and niche, comparing the newly identified niches to the most similar ones in the reference. **k**, Neighborhood composition in tumor niches. A boxplot per tumor niche and neighboring cell type represents the niche-specific distribution of the number of cells of a given cell type among the 25 physically closest cells. Boxplot elements are defined as: center line, median; box limits, upper and lower quartiles; whiskers, 1.5x interquartile range. For clarity, only cell types composing on average more than 5% and less than 60% of the neighborhood of any niche are shown. The query tumor niche is highlighted with a circle. **l**, UMAP representation of the joint reference and query NicheCompass latent GP space, colored by SPP1 ligand-receptor GP and combined interaction GP activity, as well as the gene expression of the ligand and its receptors. **m**, Heatmap of the communication strengths between niches via the SPP1 ligand-receptor GP in the query sample (donor 12) and reference sample (donor 6) – the two donors with highest macrophage infiltrates.

In donor 9, tumor cells were divided into two niches. Based on the histological images (**Fig. 5d**) and neighborhood cell composition (**Fig. 5k**), we identified that niche 1 corresponded to tumor cells in the stroma border, while niche 3 consisted of tumor cells infiltrated by neutrophils. Differential GP analysis (**Methods**) between the two niches revealed the CXCL1 ligand-receptor GP as one of the most enriched GPs in niche 3 (**Fig. 5d** & **Supplementary Fig. 23a**), with CXCL1 being a well-established chemoattractant for neutrophils^88^. This indicates that NicheCompass can distinguish niches interacting with different cell types, even with similar spatial distribution and expression profiles. Additionally, it highlights its capacity to extract GPs mediating niche-specific interactions. By further analyzing the niche 3 composition, we observed that 11% of tumor cells from donor 12 clustered into niche 3 instead of its own donor-specific niche 5 (**Supplementary Fig. 23b**). Neighborhood composition analysis within that sample revealed that neutrophils closely surround these cells (**Supplementary Fig. 23c**), showing that niches that are conserved across patients can be identified.

The distinct stroma clusters captured stromal structures that were mainly distinguished by the dominating expanded immune cell type (**Fig 5b** & **Supplementary Fig. 22c**) and its spatial arrangement (i.e. tumor infiltrating, immune aggregates or surrounding stromal tissue or immune expansions, **Supplementary Fig. 22e**). This is exemplified by two neutrophil-dominated niches that mapped closely in the latent GP space. Although their cellular composition was very similar, with some structural stroma cells, some macrophages and a high percentage of neutrophils (**Supplementary Fig. 22c**), niche 7 formed a big expansion that did not infiltrate tumor tissue, while niche 6 consisted of small expansions that infiltrated the cancerous tissue (**Fig. 5e**). This shows that by incorporating spatial and cellular interaction information, NicheCompass can identify infiltrating immune cells. Conversely, when structures were similar across donors in composition and spatial arrangement, they were correctly identified as shared structures; such is the case for lymphoid aggregates niche 11 surrounded by the plasmablast-rich stroma niche 9, present both in donor 5 and 12 and correctly identified as shared structures (**Fig. 5e**).

In summary, we built a spatial reference atlas for NSCLC, and showed that NicheCompass can integrate samples even in a context of high heterogeneity, distinguishing shared from donor-specific niches, and incorporating both spatial organization and molecular composition information, while detecting the mediating interaction GPs.

### NicheCompass discovers an SPP1+ macrophage-dominated tumor niche via spatial reference mapping

To evaluate NicheCompass’ spatial reference mapping capacity, we used a biological replicate (**Supplementary Fig. 24a**) and a sample from a new donor (**Fig. 5f**), and mapped them onto our integrated reference, using minimal, weight-restricted fine-tuning (**Methods**). To simulate a scenario where one does not have access to the whole dataset but only to a classification model trained on the reference, we additionally trained a k-nearest neighbors classifier on the reference latent GP space, and transferred the niche labels to the query cells (**Fig. 5h** & **Supplementary Fig. 24d**). Query cells were correctly integrated when mapping the biological replicate (new section from donor 5) onto our reference (batch Silhouette Width = 0.97, **Supplementary Fig. 24b**). Cells were assigned with high probability to the donor 5 specific reference niches (**Supplementary Fig. 24b,e** & **25a**), and pre-annotated cell types were correctly classified (**Supplementary Fig. 24c-d**). This shows our method’s capability to remove batch effects while preserving biological features.

A successful mapping should integrate matching niches between the reference and the query, while retaining distinct disease and donor variation. This includes accounting for the emergence of novel spatial niches that may not be present in the reference. To test our method in this context, we conducted further evaluations by mapping a completely new donor (**Fig. 5f**). Label transfer correctly distinguished tumor niches from macrophage-rich niches and niches rich in lymphoid cells (**Fig. 5g-h**). Compared to the biological replicate, we observed a low probability in some of the assignments, with 40% of the cells having a classification probability lower than 0.7 and 15% lower than 0.5 (**Supplementary Fig. 25a**). This suggested the presence of unseen niches in our query. Therefore, we decided to re-cluster the reference and query cell latent GP space together (**Fig. 5i**), which revealed in the query two lymphoid-rich stroma niches shared with donor 9 (niche 10 and 14), and two previously unseen niches with tumor cells (niche 15) and macrophages (niche 13), respectively (**Fig. 5g,i**). Closer inspection of the niches’ cellular composition and spatial distribution (**Fig. 5j**) showed between-donor similarity of the stromal niches, which overlapped with donor 9. In both donors, the stromal niches had a tumor-infiltrating localization, and were dominated either by stromal cells (niche 14) or lymphoid cells (niche 10, **Supplementary Fig. 25b**). Conversely, since all query cells were tumor infiltrating, none mapped to the second niche of stromal cells observed in donor 9 (niche 8), which did not infiltrate the tumor tissue.

Macrophage niche 13 mapped closely but differentiated from the macrophage-rich niche 12 found in the reference (**Fig. 5i**). These two clusters were distinguished in that the query niche 13 consisted of tumor-infiltrating macrophages, while the reference niche 12 macrophages were in the adjacent tissue (**Fig. 5j**). In line with this, the query-exclusive tumor niche 15 mapped closely in the latent GP space to its interacting macrophages (**Fig. 5i**). Based on neighborhood composition analysis, this was the only tumor niche that closely interacted with macrophages (**Fig. 5k**), which suggests a driving role of macrophage interaction in the formation of this segregated tumor cluster. Differential GP analysis of tumor niche 15 compared to other tumor niches revealed an over-activation of the SPP1 (secreted phosphoprotein 1, commonly referred to as osteopontin) ligand-receptor GP (**Fig. 5l**). Similarly, the SPP1 combined interaction GP, which also included the downstream targets of the ligand and receptors, was overexpressed in the infiltrating macrophage niche 13 compared to the other macrophage-rich stromal niches (**Fig. 5l**). SPP1 is a well-established driver of macrophage polarity in the TME^89^ and, as a marker of pro-tumor infiltrating macrophages, is associated with poor prognosis in lung cancer^90,91^. Closer analysis of the *SPP1* gene expression as well as its receptors confirmed that *SPP1* was over-expressed in the infiltrating macrophages niche 13 compared to the other macrophage-rich stromal niches (T-test adjusted P-value <0.001), while both the gene and its receptors (*ITGAV, ITGB1* and the transactivated *EGFR*^92^) were over-expressed in the associated tumor niche, compared to the other tumor niches (T-test adjusted P-value <0.001, **Fig. 5l**). This suggests an important role of the SPP1 pathway in this donor. To further explore this, we inferred cell-cell communication based on this GP both in the query sample and the other macrophage-rich sample, which was from donor 6 (**Fig. 5m**). Compared to the reference, the query sample had higher communication strengths both within the macrophage niche and outgoing to the rest of the niches of the sample (**Fig. 5m**).

Our analysis highlights NicheCompass’ capability to detect novel niches in terms of spatial distribution, as well as their relevant interaction GPs. To the best of our knowledge, this demonstrates, for the first time, the feasibility of mapping query spatial datasets onto single-cell resolution spatial atlases to identify novel spatial variation in the query, using minimal fine-tuning.

### NicheCompass identifies fine-grained niches in the mouse brain using multimodal gene programs

To understand the interaction mechanisms and spatial arrangement of diverse cell types within tissues, it is crucial to explore not just gene expression but also spatially resolved epigenetic factors like chromatin accessibility^15^. Leveraging multimodal GPs, including transcriptional regulation GPs that can capture gene regulatory networks (GRNs)^39,93^ across modalities (**Methods**), we trained a NicheCompass model on a spatial multi-omics mouse postnatal day 22 brain dataset generated with the Spatial ATAC-RNA-seq technology^15^ (**Methods**). After clustering the latent GP space learned by NicheCompass, we annotated the identified clusters with functional and anatomical niche labels. In comparison with an anatomical map from the Allen Mouse Brain Reference Atlas^74^ (Mouse, P56, Coronal, Image 44), we found that the identified niches closely corresponded to anatomical subcomponents, uncovering multiple cortical layers, compartments in the striatum, and fiber tracts (**Fig. 6a**). Strikingly, NicheCompass could successfully delineate between two commissural fiber niches, the Corpus Callosum and Anterior Commissure. Both were dominated by oligodendrocyte lineage cell types (myelinating and newly formed oligodendrocytes) with only slight compositional differences (presence of astrocytes in the Corpus Callosum versus vascular and leptomeningeal cells in the Anterior Commissure). This shows that our model can resolve anatomically distinct but molecularly similar niches (**Fig. 6b**).

**Fig. 6.**
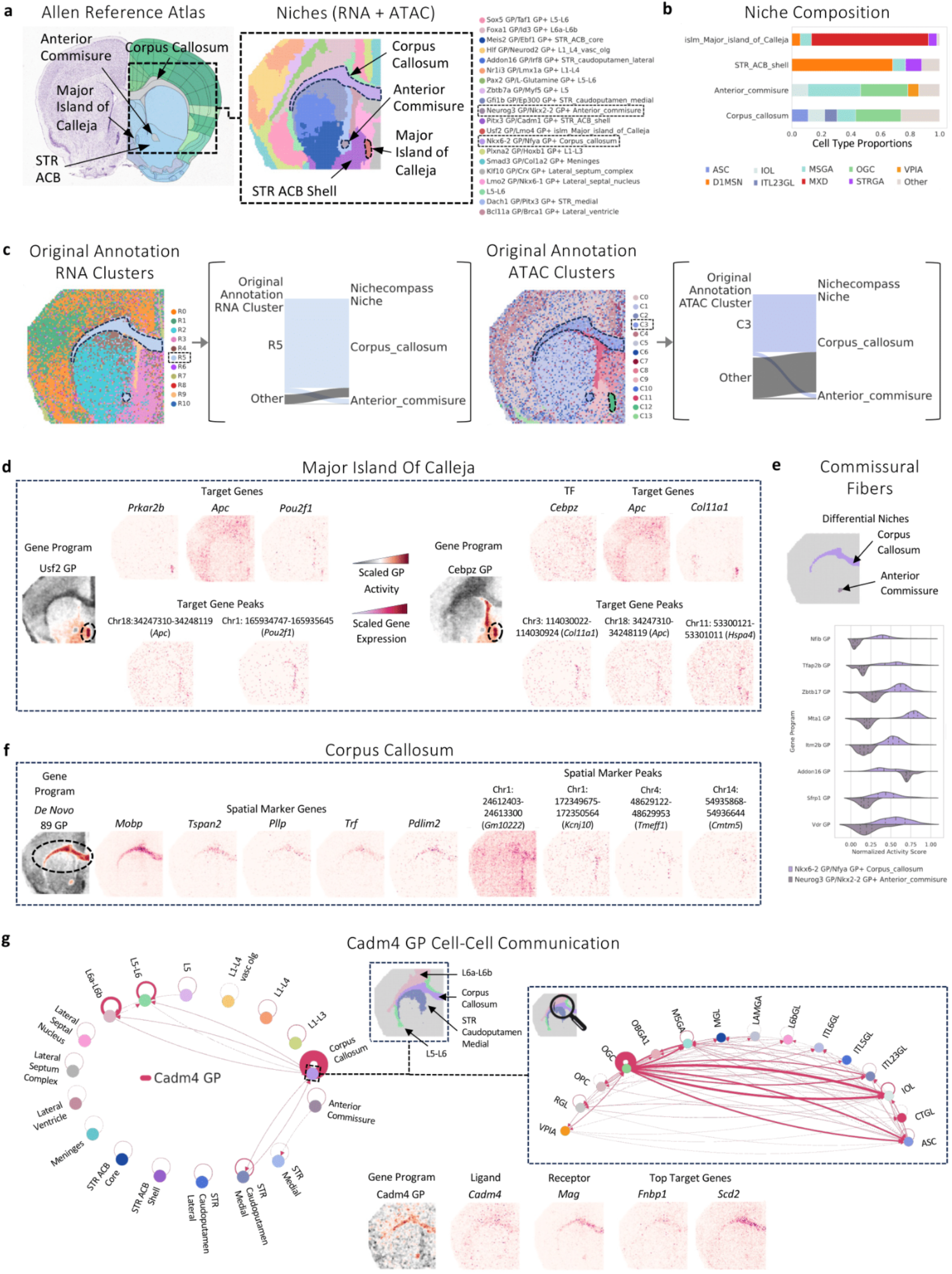
NicheCompass integrates spatial gene expression and chromatin accessibility to learn multimodal gene programs in the mouse brain. **a,** A coronal mouse brain image from the Allen Reference Atlas showing selected subcomponents in the striatum and fiber tracts; next to it the mouse brain tissue sequenced with the Spatial ATAC-RNA-seq technology, highlighting corresponding niches obtained with NicheCompass. The nomenclature of niches is derived from two characterizing GPs and region identity. Region abbreviations are STR: striatum; ACB: nucleus accumbens. **b,** Niche-specific cell type composition (based on Zhang et al.^15^ cellular annotations for each pixel) for highlighted niches from **a**. The annotated cell types are ASC: astrocytes, grey matter; IOL: newly formed oligodendrocytes; MSGA: inhibitory neurons, medial septal; OGC: myelin forming oligodendrocytes; VPIA: vascular and leptomeningeal cells; D1MSN: D1 like medium spiny neurons, striatum; ITL23GL: excitatory neurons, cortex L2/L3; STRGA: *Foxp2* positive inhibitory neurons; MXD: matrix D1 neurons, striatum. **c,** The mouse brain tissue colored by cluster annotations from the original authors for the RNA and ATAC modalities, respectively. Sankey plots illustrate how the original cluster labels map to the identified niches. **d,** Two multimodal transcriptional regulation GPs enriched in the Major Island of Calleja niche. GP activities align with the expression of the target genes and the transcription factor, as well as with target gene-associated peaks. **e,** Cellular activities of differential GPs in the Corpus Callosum and Anterior Commissure niches. **f,** A *de novo* GP enriched in the Corpus Callosum niche, the expression of the most important genes, and openness of the most important peaks. **g,** A detailed analysis of the Cadm4 GP enriched in the Corpus Callosum niche. The first circle plot displays inferred niche-to-niche cell-pair communication strengths. Interactions within the Corpus Callosum niche dominate the GP with fewer interactions between the Corpus Callosum niche and neighboring niches L5-L6, L6a-L6b, and STR Caudoputamen Medial. The GP activity and expression of ligand, receptor and top target genes substantiate this analysis. Zooming into interactions involving the Corpus Callosum niche highlights communicating cell types. RGL: radial glia-like cells; OPC: oligodendrocytes precursor cells; OBGA1: inhibitory neurons, olfactory bulb; MGL: microglia; LAMGA: CGE-derived neurogliaform cells, Lamp5; L6bGL: excitatory neurons, cortex L6b; ITL6GL: excitatory neurons, cortex L6; ITL5GL: excitatory neurons, cortex L5; CTGL: excitatory neurons, cortex CT.

To evaluate the advantage of incorporating epigenetic information, we further trained a unimodal NicheCompass model on only the RNA modality. We observed that while most niches could be accurately recovered, the Major Island of Calleja (Insula Magna) niche was only detectable when additionally including the ATAC modality and would otherwise not be separable from the Striatum Nucleus Acumbens Shell niche (**Fig. 6a** & **Supplementary Fig. 26**). This finding underscores the value of incorporating epigenetic information through ATAC measurements in the niche identification process. Moreover, we compared our identified niches with the modality-specific annotations from the original study^15^ (**Fig. 6c**). Similar to our RNA-only model, the Major Island of Calleja was not identifiable as a separate niche in the original RNA-based clusters but manifested in the ATAC-based clusters, additionally verifying the importance of epigenetic characterization. In contrast to both original annotations, NicheCompass disentangled the Corpus Callosum and Anterior Commissure niches, subdivided the striatum into its different functional compartments, and recovered niches with a unique resolution and superior spatial consistency without mixing of clusters (**Fig. 6a,c**).

Following the niche identification process, we pursued our previously introduced niche characterization workflow. First, we built a niche hierarchy to decipher the tissue’s global spatial organization (**Supplementary Fig. 27a**). We observed that niches mainly clustered into higher-order anatomical structures such as the cortical layers constituting a Cortex cluster (1) and spatially adjacent niches of the striatum grouping into two Striatum clusters (2,4). Due to their molecular similarities, the Corpus Callosum and Anterior Commissure niches formed a commissural fiber cluster (3), despite being located in distant tissue regions. In addition, the characterizing GPs (**Methods**) of each niche again showed distinctive signatures, consistent with the identified hierarchy (**Supplementary Fig. 27b**), and niches adjacent in the hierarchy showed similar cell type composition (**Supplementary Fig. 27c-d**).

We then investigated the Corpus Callosum, Anterior Commissure and Major Island of Calleja niches more in detail. The Major Island of Calleja has been described to contain dopamine D3 receptor-expressing granule cells^94^ involved in depression^95^, and to express *Gad65*^96^ and *Nos1* (Nitric Oxide Synthase 1)^97^; accordingly, we observed spatially localized *Drd3* expression^94^ (**Supplementary Fig. 28**). However, it has not yet been characterized in detail at the molecular level, presenting an opportunity for exploration with NicheCompass. GP enrichment analysis (**Methods**) of the Major Island of Calleja niche revealed interesting transcriptional regulation GPs with a multimodal footprint (**Fig. 6d**). These included the Usf2 transcriptional regulation GP, driven by several target genes (*Prkar2b*, *Apc*, *Pou2f1*) and associated peaks with signal in the Major Island of Calleja niche. The regulatory interaction between *Usf2* and *Prkar2b* plays a central role in the regulation of glycolitic glucose metabolism^98^, while *Apc* and *Pou2f1* have also been linked to glycolysis^99–101^, a key pathway for brain physiology^102^. Further, we found upregulation of the Cebpz transcriptional regulation GP, driven by the transcription factor (TF) gene *Cebpz*, the target genes *Col11a1* and *Apc*, and various target gene-associated peaks with correlated signals. *Cebpz* has been functionally related with these target genes in colorectal and gastrointestinal cancer, respectively^103,104^; however, there is yet no evidence for an interaction in the brain, rendering it a candidate for future investigation. Moreover, we found enrichment of various additional GPs (**Supplementary Fig. 29a**), including the Rreb1 transcriptional regulation GP, driven by the TF gene *Rreb1*, one of few marker genes for a GABAergic spiny projection neuron subpopulation specific to the Islands of Calleja^105^, and an interesting *de novo* GP that comprised various correlated peaks with distinctive signal in the Major Island of Calleja niche (**Supplementary Fig. 29b**). Overall, these examples illustrate how peaks can enhance gene-based GPs and elucidate transcriptional regulatory mechanisms.

The Corpus Callosum and Anterior Commissure connect different areas within the two brain hemispheres and are mainly composed of oligodendroglia and fibers of myelinated axons. While these fibers can be readily distinguished anatomically, such distinction has not yet been achieved at a molecular level. To investigate spatially-induced molecular differences between them, we conducted differential testing between cells belonging to these niches. Our model identified several GPs that significantly delineated the two niches (**Fig. 6e**) and contained multiple important genes with unequivocal gene expression differences between the two niches (**Supplementary Fig. 30**). Moreover, closer inspection of the Corpus Callosum niche revealed extensive enrichment of diverse prior GPs from different categories (**Supplementary Fig. 31**) and of a *de novo* GP (**Fig. 6f**). Among the enriched prior GPs, we detected transcriptional regulation GPs, including the Klf6 GP, whose TF *Klf6* is required for CNS myelination^106^, ligand-receptor GPs, including the Cd82 GP, driven by the *Cd82* and *Gsn* genes both known to be involved in the regulation of oligodendrocyte differentiation^107,108^, and metabolite-sensor GPs, including the Glycine GP, driven by the *Gatm* gene, which plays a role in the synthesis of creatine in oligodendrocytes^109^. These prior GPs are meaningful in the context of the Corpus Callosum niche mainly consisting of oligodendrocytes, one function of which is the myelination of axons in the central nervous system. The detected *de novo* 89 GP was highly enriched specifically in the Corpus Callosum niche and comprised several highly correlated, spatially variable genes (*Mobp*, *Tspan2*, *Pllp*, *Trf*, *Pdlim2*), of which *Mobp* is an oligodendrocyte marker gene and the remaining genes are related with myelinating oligodendrocytes^110–113^ and have been classified under a Gene Ontology category specific to myelination processes.

Lastly, we investigated cell-cell communication within and between niches and cell types. For this, we focused on two combined interaction GPs enriched in the Corpus Callosum niche, utilizing our previously introduced cell-cell communication potential scores and cell-pair communication strengths (**Methods**). The first GP, Cadm4 combined interaction GP, plays a key role in regulating axoglial adhesion during CNS myelination^114^. We found that activity in this GP was driven by highly spatially correlated genes, including the ligand gene *Cadm4*, the receptor gene *Mag,* and target genes *Fnbp1* and *Scd2* (**Fig. 6g**). Computing communication strengths, we found that interactions occurred mostly within the Corpus Callosum niche with fewer interactions involving the cortical layer niches L5-L6 and L6a-L6b (**Fig. 6g**). In terms of interacting cell types in this GP, oligodendrocytes were the main interaction source and target entity with different neuron subpopulations acting as additional targets. This is aligned with the fact that myelination in the central nervous system (CNS) involves a complex set of intercellular contacts between oligodendrocytes and their underlying axons^114^. The second GP, Cldn11 combined interaction GP, represents a critical interaction pathway for assembling tight junctions in CNS myelin^115^. This GP was driven by the *Cldn11* gene, which was highly differentially expressed in the Corpus Callosum niche, in both source and target cells, consistent with *Cldn11* acting as the ligand and receptor in this GP (**Supplementary Fig. 32**). Next to *Cldn11*, the GP also picked up two downstream target genes, *Pde8a*, known to be expressed in murine oligodendrocytes^116^, and *Ugt8a*, involved in the synthesis of myelin-related lipids^117^. In accordance with its functional role in the formation of tight junctions in CNS myelin, interactions utilizing the Cldn11 GP were almost exclusively situated within the Corpus Callosum niche and mainly limited to pair-wise oligodendrocyte interaction partners.

### NicheCompass can construct spatial atlases of entire organs across millions of cells

We concluded our analysis by demonstrating the capability of NicheCompass to scale to millions of cells and construct a spatial atlas of a whole organ. To this end, we first applied it on a mouse brain spatial dataset (**Methods**) with over one million cells, profiled using a panel of 1,022 genes across 17 coronal and 3 sagittal slices^17^. Clustering of the NicheCompass latent GP space identified 15 niches that were successfully aligned across sequential tissue sections (**Supplementary Fig. 33a-b**). We observed that all recovered niches were consistently composed of multiple cell types from various tissue sections (**Supplementary Fig. 33c**), highlighting the efficacy of NicheCompass in a whole organ setting. Further analysis of niches revealed that they corresponded to anatomical regions from the Allen Mouse Brain Reference Atlas^74^ (**Supplementary Fig. 34**), demonstrating accurate identification of tissue organization.

Finally, we validated the scalability of NicheCompass toward creating foundational spatial cell representations by integrating 8.4 million cells from a different, recently published, whole mouse brain dataset (**Methods**). This dataset was profiled using 1,100 genes from 239 tissue sections, including 213 coronal and 26 sagittal slices from four different mice, making it by far the largest spatial atlas. To the best of our knowledge, this is the first time that a spatial omics dataset of such scale was integrated with spatially aware representation learning. The learned representations showed that NicheCompass could successfully integrate cells from matching brain regions (**Supplementary Fig. 35a-b**), resulting in spatially consistent niches (**Supplementary Fig. 35a**) across mouse donors (**Supplementary Fig. 35c**).

In conclusion, this demonstrates that NicheCompass can assemble maps of molecular and cellular organization at organ level across different individuals, paving the way toward construction of whole organism spatial atlases^118^.

## Discussion

We introduced NicheCompass, the first approach to quantitatively characterize tissue niches based on the principles of cellular communication. We benchmarked our approach qualitatively and quantitatively against existing methods for spatial representation learning, highlighting NicheCompass’ accurate identification of anatomical niches and superior performance across multiple datasets and scenarios (**Fig. 2**). Our scalable design and implementation allow seamless scaling to datasets with millions of cells (**Supplementary Fig. 33-35**), a crucial requirement for building and contributing to large-scale spatial atlas projects like the Human Cell Atlas (HCA)^119^, as well as digital pathology analyses. Finally, we demonstrated the extensibility of our approach to enable spatial reference mapping for iterative integration (**Fig. 5f-i**), and to learn from spatial multi-omics data, providing enhanced niche characterization through multimodal GPs (**Fig. 6**).

We used NicheCompass’ interpretability and analysis capabilities to elucidate tissue architecture and important cellular interaction processes during mouse organogenesis, in the adult mouse brain, and for different human cancer phenotypes, highlighting niche-specific GP activity and cell-cell communication patterns. As evidenced by these applications, NicheCompass offers an innovative and comprehensive workflow to quantitatively characterize tissue niches and analyze spatial omics data; nonetheless, future extensions will be beneficial to further improve the quality and applicability of this workflow. For instance, similar to other prior knowledge-based methods, NicheCompass is limited by incomplete, noisy and inadequate information in the underlying databases. While we mitigate this issue through GP pruning, gene sparsity and *de novo* GPs (**Methods**), which are useful mechanisms to remove low quality priors and learn spatial correlations beyond prior knowledge, the identified genes by *de novo* GPs might not be linked to molecules that can structurally interact (e.g. ligands and receptors). Incorporating structural protein information, for example with insights from AlphaFold2^120,121^, could therefore be an interesting avenue to increase the biological relevance of inferred GPs. In terms of prior GPs, a comparable limitation regarding causality can arise. This is the case when prior GPs have high activity despite absence of a causal mechanism, for instance if a GP is dominated by target gene expression without expression of the causal ligand or TF genes. We partly address this through selective regularization based on gene functions (**Methods**); nonetheless, we recommend to interpret GP activities in combination with expression of important GP genes, as we have done in the selection of characterizing GPs (**Methods**). Another limitation of the current GP design is that GPs are always composed of the same genes reflected in the gene importances obtained from the omics decoder weights (**Methods**). These cannot vary per cell and therefore suboptimally reflect biological reality in which different target genes of a signaling pathway might be activated based on the cell’s intrinsic characteristics. A dynamic GP design with attentive mechanisms coupled to a cell’s representation could be an idea to account for this. Finally, to fully utilize the potential of prior GPs, it is necessary that datasets are not limited by small gene panels. Whereas many currently available experimental technologies are either limited in resolution or throughput and show uneven coverage of the ligand and receptor genes due to high sparsity and noise, NicheCompass will benefit from further innovations leading to high-resolution, high-throughput spatial readouts^122^. Future database enhancements and the detection of new interaction pathways will further strengthen NicheCompass’ capabilities.

Although having demonstrated NicheCompass’ applicability to spot and single-cell resolution datasets, we believe that an architectural incorporation of spot deconvolution could increase its relevance for the widely available spot-level spatial transcriptomics data, such as 10x Genomics Visium. This will allow more accurate inference of cell-level processes in these datasets. Another important aspect to improve NicheCompass’ applicability is the expansion of its predictive capabilities to more accurately transfer niche annotations from spatial reference atlases and identify the effects of disease-specific niches on clinical patient outcomes. We believe this can be achieved by adding supervised fine-tuning after self-supervised pre-training^31,123^. Furthermore, while spatial reference mapping holds great potential in iteratively integrating datasets, it requires comprehensive reference atlases with many cells from multiple batches and ideally different datasets^124^. In this context, another important caveat is that integration requires availability of the same genes across different samples, which can be challenging given the limited gene panels of current technologies. Finally, architectural extensions may further enhance niche identification performance and the inference of cellular interactions at larger length scales. These could include employing alternative graph-based encoders, such as graph transformers^125^ and graph adaptations of state space models^126,127^, and incorporating additional modalities into NicheCompass’ multimodal architecture (such as histone modifications, chromatin organization, or protein expression), which will provide an even more holistic niche characterization.

With the increasing availability of spatial omics datasets, we expect NicheCompass to become an essential tool to quantitatively characterize tissue niches in health and disease, empowering clinicians and cell biologists to understand the cellular dynamics of tissue response to injury and infection.

## Data availability

All datasets analyzed in this manuscript are publicly available, referenced in the manuscript, and accessible for download as described at https://github.com/Lotfollahi-lab/nichecompass-reproducibility.

## Code availability

NicheCompass is available as a Python package, maintained at https://github.com/Lotfollahi-lab/nichecompass. The code to reproduce our analyses and benchmarking experiments can be retrieved from https://github.com/Lotfollahi-lab/nichecompass-reproducibility. Documentation is provided at https://nichecompass.readthedocs.io/.

## Acknowledgments

S.B. and I.B.-P. acknowledge support from the Munich School for Data Science. M.L. appreciates feedback and fruitful discussions with Kerstin Meyer and Giovanni Ciriello regarding cancer applications. M.L. acknowledges financial support from the Joachim Herz Stiftung. G.C-.B. acknowledges financial support from the Knut and Alice Wallenberg Foundation (grants 2019-0107, 2019-0089), the Swedish Cancer Society (Cancerfonden, 190394 Pj), and the Swedish Brain Foundation (FO2023-0032). S.B. is thankful to Sergei Rybakov and all members of the Lotfollahi Group for valuable feedback, in particular Stathis Megas, Amirhossein Vahidi, Kevin Ly, Marie Moullet and Erick Antonio Armingol Gonzalez. S.B. is grateful to Paula Villa Fulton for feedback on figure design and to Paula, Mickey, and Octavious O. Villa Fulton for their inspirational support. We thank Di Zhang for providing cell type annotations for the spatial ATAC-RNA seq mouse brain dataset.

## Author contributions

M.L. conceived the project. S.B. designed the algorithm with feedback from M.L. S.B. implemented the algorithm. M.L. designed the experiments with contributions from S.B. and C.T.-L. S.B. performed the benchmarking experiments. A. Boxall curated the STARmap PLUS mouse data and analyzed it with contributions from S.B. and M.L. C.T.-L. curated the Xenium human breast cancer data and C.T.-L., S.B. and A. Maguza analyzed it. I.B.-P. performed the analysis of the NanoString CosMx human NSCLC data with contributions from S.B., M.L. and C.T.-L. S.B. curated the remaining datasets and performed all other analyses with contributions from M.L., A.M.F., F.M., O.A.B., E.A., G.C.-B, and R.F. F.J.T. supported M.L. during his work on the project and provided the environment to perform the work. S.B., I.B.-P., M.L., and C.T.-L. wrote the manuscript with contributions from A. Boxall. M.L. and C.T.-L. supervised the research. All authors reviewed the manuscript.

## Supplementary information

### Supplementary Note 1: Applications

NicheCompass’ model design enables a comprehensive workflow to analyze spatial omics data. First, disparate tissue samples can be integrated to build large-scale, comprehensible spatial atlases. Effective integration is achieved through sample-specific neighborhood subgraphs (**Methods**), which dispense of prior coordinate alignment, and covariate embeddings that regress out unwanted batch effects. High scalability is obtained with a memory-efficient framework based on PyTorch Geometric^128^, including sparse and memory-efficient internal data representations, graph mini-batch data loaders, and model training features such as neighbor sampling. Consequently, our model can handle millions of cells/spots seamlessly. Comprehensibility is obtained via the interpretable latent space of GPs^42^, whose spatially localized activities can be statistically tested to interrogate differences between samples. For instance, tissue samples from different donors can be integrated to interrogate differences in molecular mechanisms, and to decipher spatial biomarkers of disease phenotypes. If a particular sample does not integrate with others, its difference can be explained based on differential GP activity. Second, our model allows for highly performant, biologically informed niche identification via clustering of the obtained GP representations. The resulting clusters represent niches that are distinctly defined based on the common cellular processes of its cells, as reflected in shared activity patterns of GPs. Differential testing of GP activity can then be conducted to elucidate niche-specific cellular processes, and to contrast niches of interest. Varying the cluster resolution can further uncover interesting niche hierarchies, reflecting different levels of tissue organization. Third, cellular GP activity can be used to annotate and characterize the identified niches in detail. Moreover, our interoperable Python package (https://github.com/Lotfollahi-lab/nichecompass), built with a backbone supporting the established scverse ecosystem^129^, simplifies further functional niche characterization and frictionless analyses at spatial^130^ and single-cell^131^ level. For instance, niches can be additionally characterized through cell type composition and neighborhood enrichment of specific cell types. Fourth, the learned activity of cell-cell communication and combined interaction GPs can be used to infer cellular signaling processes between cells. In the presence of cell type labels, this can be utilized to infer cell-type-specific interaction patterns in a niche, thereby adding a further dimension of delineation between niches. Fifth, our method empowers iterative integration of samples through spatial reference mapping via transfer learning^124^. This is particularly useful when training data is not available simultaneously, as is the case for most large-scale atlassing efforts spanning across datasets. In these cases, initially a reference can be built and characterized, and, subsequently a query can be mapped through transfer learning followed by niche annotation transfer. The reference could reflect a collection of healthy samples and the query a diseased sample. If the niche annotation transfer unveils niches that are not present in the reference, these can be characterized based on differential GP activity.

### Supplementary Note 2: Additional characterization of gut and brain niches

We identified additional meaningful enriched GPs in gut and brain niches. In the Ventral Gut niche, we observed enriched activity of the Indian Hedgehog (Ihh) ligand-receptor GP (**Supplementary Fig. 8**), suggesting signaling between the Ihh protein and the multipass membrane protein Smoothened (*Smo*), which is crucial for healthy gut development during mouse organogenesis. Mutations affecting the pathway lead to a variety of malformations of the murine gastrointestinal tract^132–134^. In the Dorsal Gut niche, we identified considerable upregulation of the Pdgfc combined interaction GP (**Supplementary Fig. 8**), which, similar to the Cthrc1 combined interaction GP, was specific to the notochord niche. Notably, its ligand gene *Pdgfc* is known to be expressed in the notochord^135^.

Concerning the brain niches, we found enriched activity of the Gdf10 combined interaction GP in the Hindbrain niche (**Supplementary Fig. 9**), primarily driven by the *Gdf10* ligand gene. *Gdf10* is known to be involved in Bergmann glial cell development under *Shh* regulation in the cerebellum, a part of the hindbrain^136^. Aligned with this finding, the Sonic Hedgehog (Shh) combined interaction GP, a well known signaling pathway during brain development and morphogenesis^137,138^ was active across all brain niches with enriched activity in the neighboring Floor Plate niche where the *Shh* gene was also highly expressed (**Supplementary Fig. 9**). In the Midbrain niche, we observed enriched activity of the Efna2 combined interaction GP (**Supplementary Fig. 9**), driven by the *Efna2* gene. This gene encodes the ephrin A2 protein, crucial in region-specific apoptosis during early brain development^139^. Finally, in the Forebrain niche, there was upregulation of the Psck1n ligand-receptor GP (**Supplementary Fig. 9**), which highly correlated with the *Psck1n* gene expression.

### Supplementary Note 3: SlideSeqV2 mouse hippocampus benchmarking

In addition to spatially aware methods, we also applied two non-spatially aware deep generative methods, namely scVI^140^ and expiMap^42^, to assess whether they capture niches based on tissue anatomy in their latent representations. On the one hand, as expected, the resulting latent representations were more influenced by intrinsic cell type variation, leading to spatially scattered clusters. This reflects that cells of the same cell type are located in different parts of the tissue, and highlights the need for spatially aware methods to adequately capture spatially consistent niches of heterogeneous cells. On the other hand, however, these methods also partially recovered tissue anatomy independent of cell type, with some spatially adjacent cells of different cell types located close to each other in the latent feature space (**Supplementary Fig. 12c-d**). This emphasizes that spatial effects are already evident in single-cell transcriptomes (without considering the microenvironment), necessitating future method development to investigate the disentanglement of intrinsic and spatially-induced variation.

Moreover, we also performed clustering on the spatial coordinate feature space to investigate whether cell positioning alone would lead to reasonable niches (**Supplementary Fig. 12e**). This resulted in spatially consistent niches, yet these did not correspond to anatomical structures, highlighting the importance of incorporating molecular information.

### Supplementary Note 4: Metrics

We argue that the utility of latent representations in the context of niche identification depends on fulfilling various, partially adversarial criteria, reflected in the categories spatial conservation and niche coherence. First, latent cellular representations ought to preserve global spatial structures. It should be encouraged that tissue patterns of spatially co-occurring cell types are preserved in the latent feature space of the model, and that the spatial organization of a tissue at different resolutions is captured, so that niches representing fine-grained substructures merge to build niches of coarser structures, thus allowing the identification of niches at different levels of detail. To compute our overall score, we therefore include two metrics to measure global spatial conservation of the tissue architecture: Cell Type Affinity Similarity (CAS) and Maximum Leiden Adjusted Mutual Information (MLAMI) (**Methods**). Second, cellular representations of niches must closely abide by local spatial structure as spatial colocalization of cells is one of the defining factors of a niche. This means that proximal cells in the tissue should also be nearby in the latent feature space of the model so that niches are composed of neighboring cells in physical space. We measure this local dimension of spatial conservation through two additional metrics: Cell Type Local Inverse Simpson’s Index (CLISIS) and Graph Connectivity Similarity (GCS) (**Methods**). Third, it is important that niches are biologically meaningful, and do not merely consist of a random colocated group of cells, which is the case when molecular features are not considered (**Supplementary Fig. 12e**). For this, we measure on the one hand how compact niches are within themselves while being distinct from neighboring niches, using the Niche Average Silhouette Width (NASW) (**Methods**), and on the other hand how pure niches are in terms of cell type composition, using the Cell Type Normalized Mutual Information (CNMI) (**Methods**). We combine these two metrics into the niche coherence score. Taken in isolation, the niche coherence score encourages clear separation of the latent space by cell types. However, to compare methods holistically, our metrics enforce striking a balance with spatial preservation metrics by aggregating metric scores into an overall score through normalization followed by equal weighting of the two categories spatial conservation and niche coherence (**Methods**). This ensures that niches are spatially and biologically coherent.

### Supplementary Note 5: Scalability and speed

While NicheCompass seamlessly scaled to large datasets, other methods (except for CellCharter) were limited by relying on loading the full graph into memory (instead of mini-batch training). This led to memory overflow when loading big datasets on our 40GB GPU (**Supplementary Fig. 17**). In smaller data regimes, other methods exhibited faster runtimes, mainly due to the graph attention mechanism employed by our model. In fact, a variation of our model, NicheCompass Light, which used convolutional layers instead, showed competitive runtimes in the small data regime (**Supplementary Fig. 17**), and performed comparably to NicheCompass regarding the overall score (**Supplementary Fig. 15-16**).

---

**Supplementary Fig. 1.**
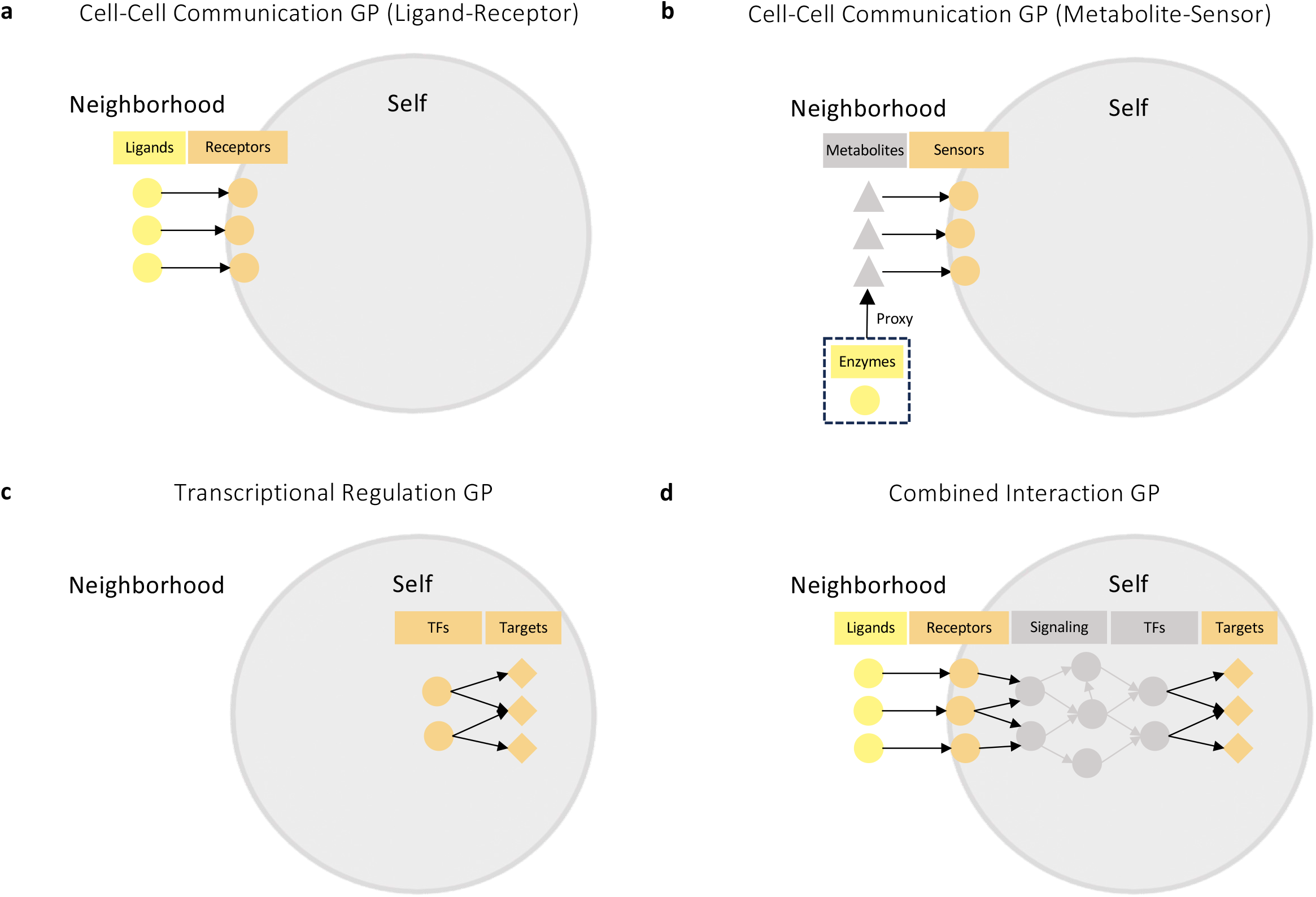
Categorization of prior gene programs. **a,** Ligand-receptor gene program (GP) comprising ligands in the neighborhood-component and receptors in the self-component. **b,** Metabolite-sensor GP comprising metabolites in the neighborhood-component and sensors in the self-component. Enzyme expression serves as a proxy for metabolite presence. **c,** Transcriptional regulation GP comprising transcription factors and target genes in the self-component. **d,** Combined interaction GP comprising ligands in the neighborhood-component and receptors and target genes in the self-component. Circles represent proteins, triangles represent metabolites and rhombi represent genes involved in the interactions.

**Supplementary Fig. 2.**
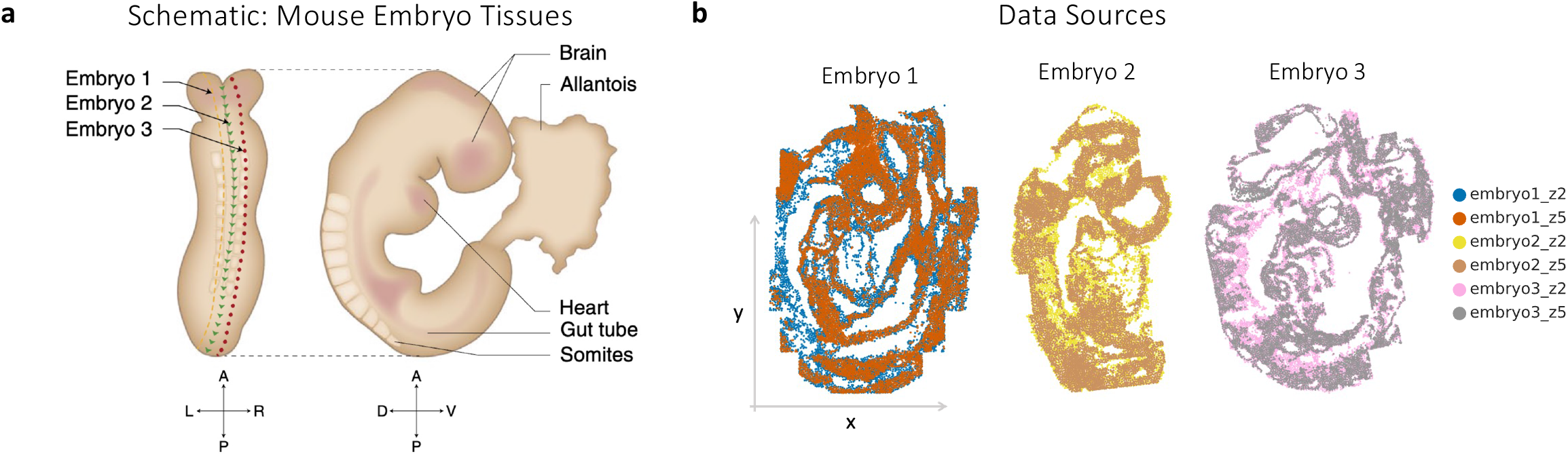
Mouse organogenesis tissue samples. **a,** Schematic of the 8-12 somite stage mouse embryos from the original authors. Dotted lines indicate the estimated position of the sagittal embryo tissues shown in **b**, illustrating slight deviations on a left-right axis. **b,** Embryo tissues colored by section (data source), which is used as covariate during model training to remove batch effects.

**Supplementary Fig. 3.**
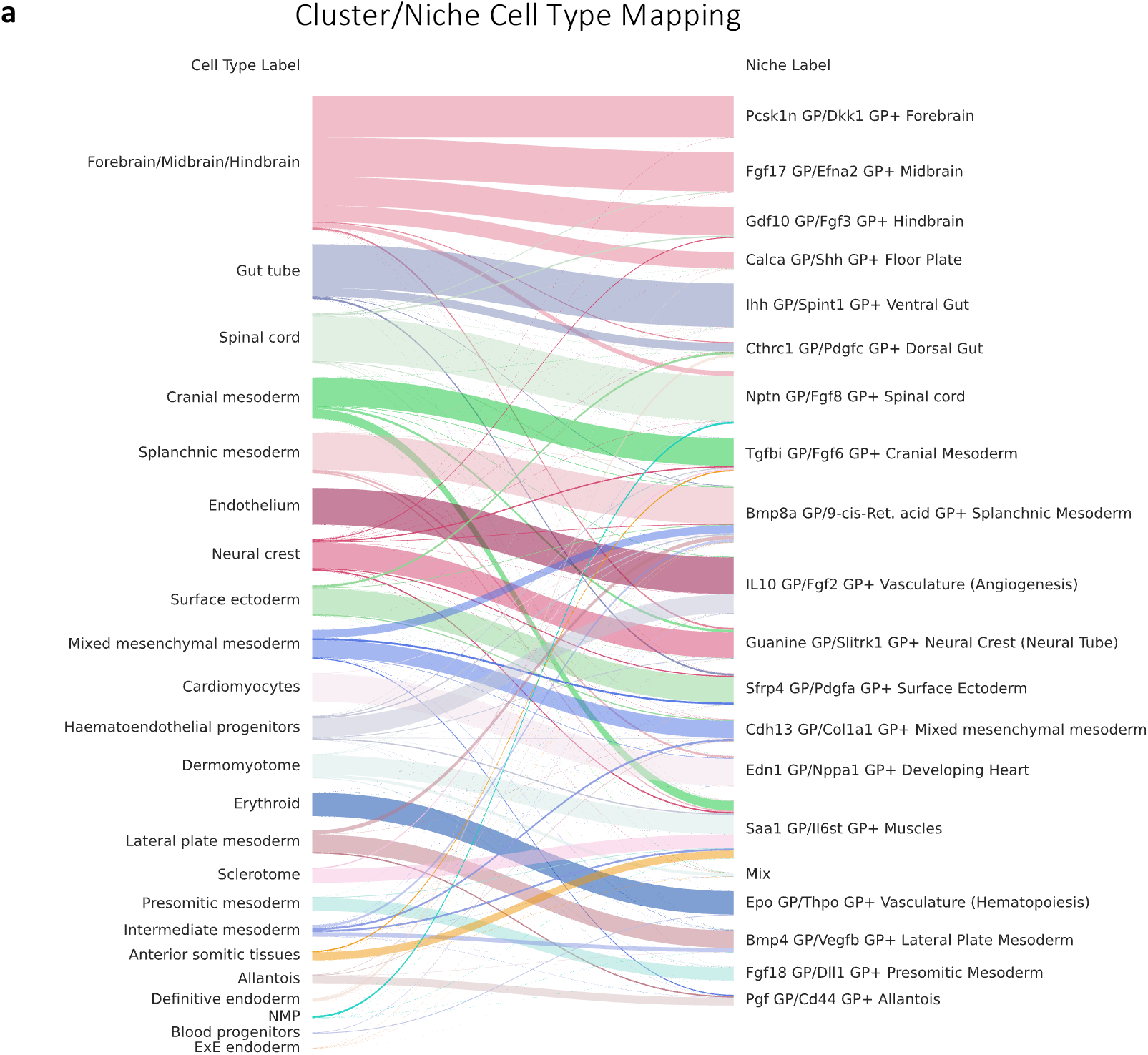
Mouse organogenesis cluster/niche cell type mapping. **a,** Mapping of the original cell type/region annotations to the clusters obtained from clustering the NicheCompass latent GP space, used for the annotation of clusters with niche labels.

**Supplementary Fig. 4.**
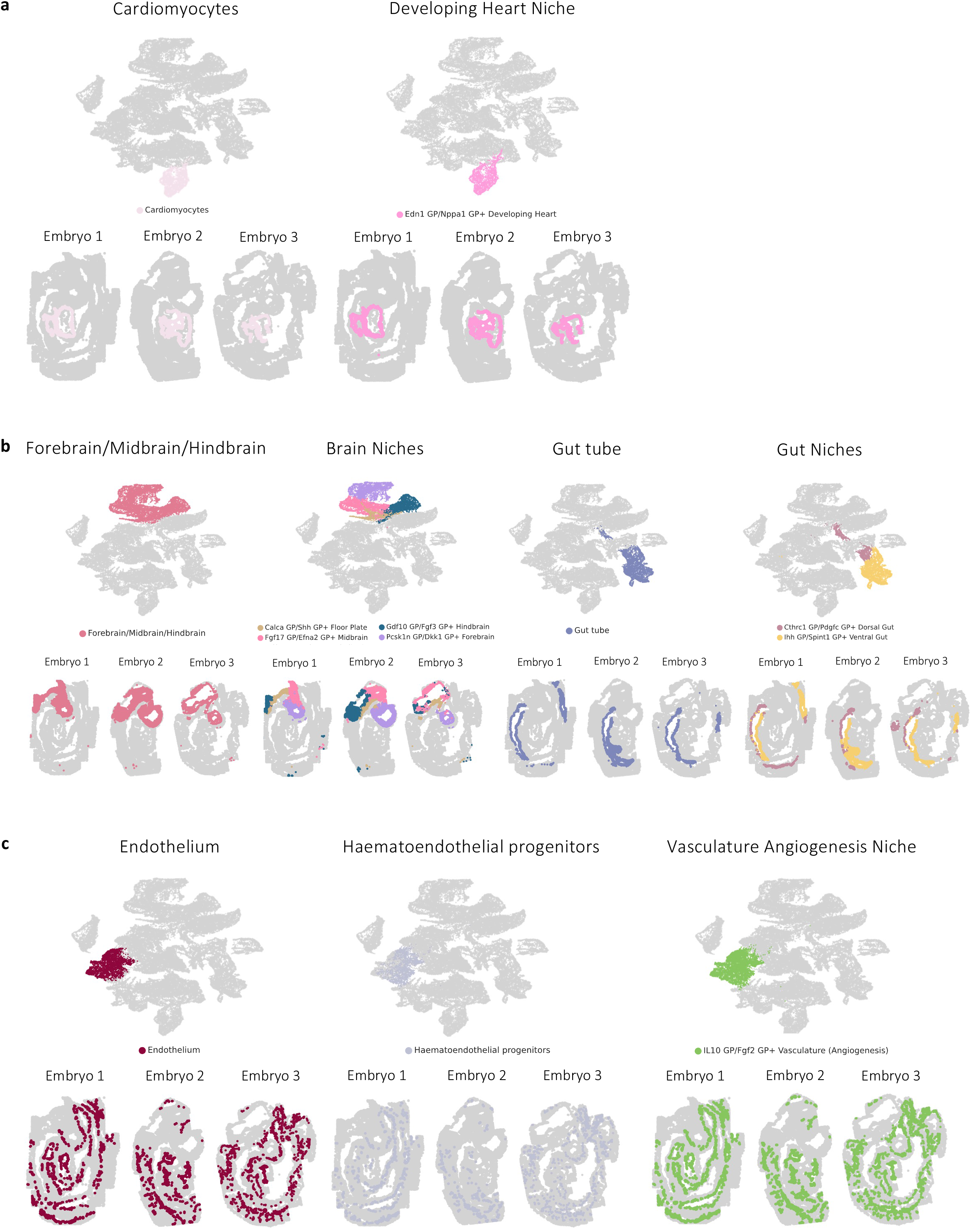
Mouse organogenesis niche heterogeneity. **a,** The developing heart niche as an example of a niche consisting of a homogenous cell population containing all cells of a certain cell type, as illustrated by an adjacent visualization of cardiomyocyte cells. **b,** The brain and gut niches as an example of niches consisting of a homogenous cell population but only containing a subset of all cells annotated with the corresponding cell type, as illustrated by adjacent visualizations of the forebrain/midbrain/hindbrain and gut tube cells. **c,** The vasculature angiogenesis niche as an example of a niche consisting of heterogeneous cell populations. Only cells that are part of the respective niches are colored.

**Supplementary Fig. 5.**
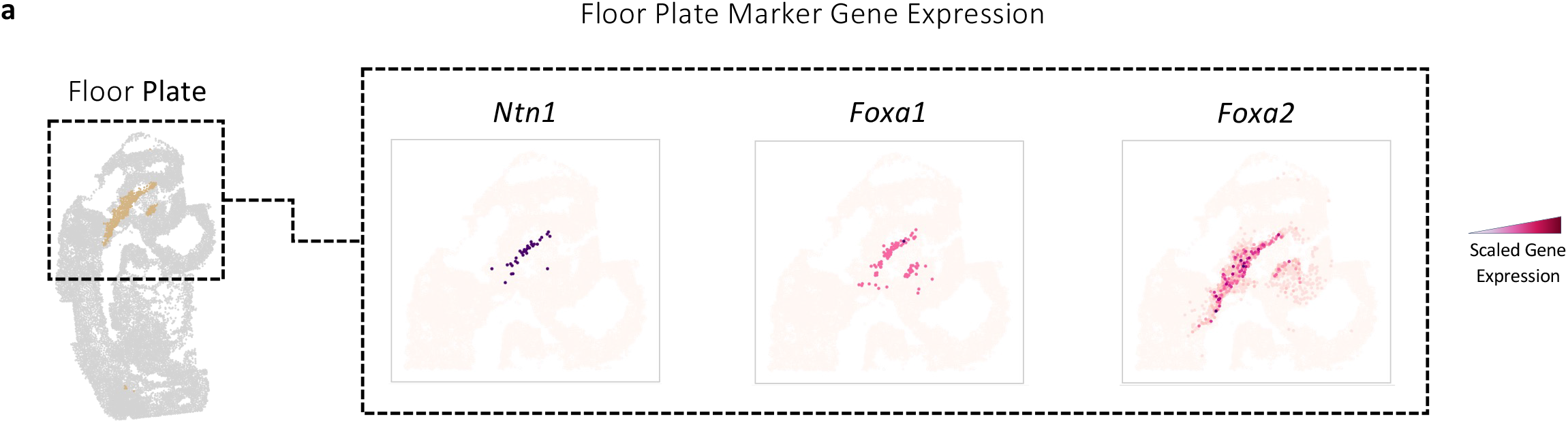
Floor plate marker genes. **a,** The Floor Plate niche of the second embryo and the gene expression of three bona-fide floor plate marker genes, validating the identification and annotation of the niche.

**Supplementary Fig. 6.**
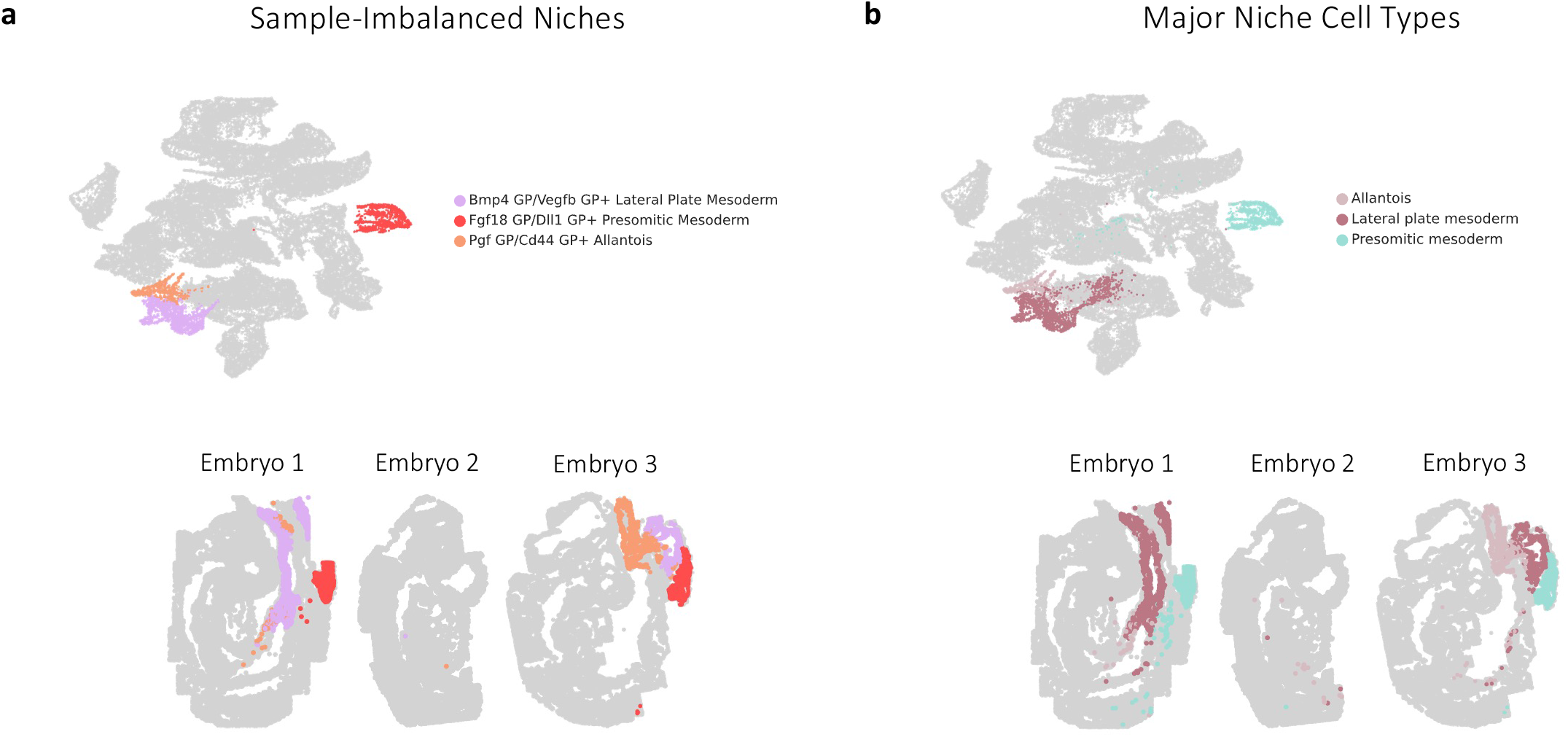
Imbalanced niches across samples and their corresponding major cell types. **a,** Three niches that were imbalanced across samples with considerable presence only in the first and third mouse embryo sample. **b,** The three corresponding major cell types of the niches from **a** show similar presence in only the first and third mouse embryo samples.

**Supplementary Fig. 7.**
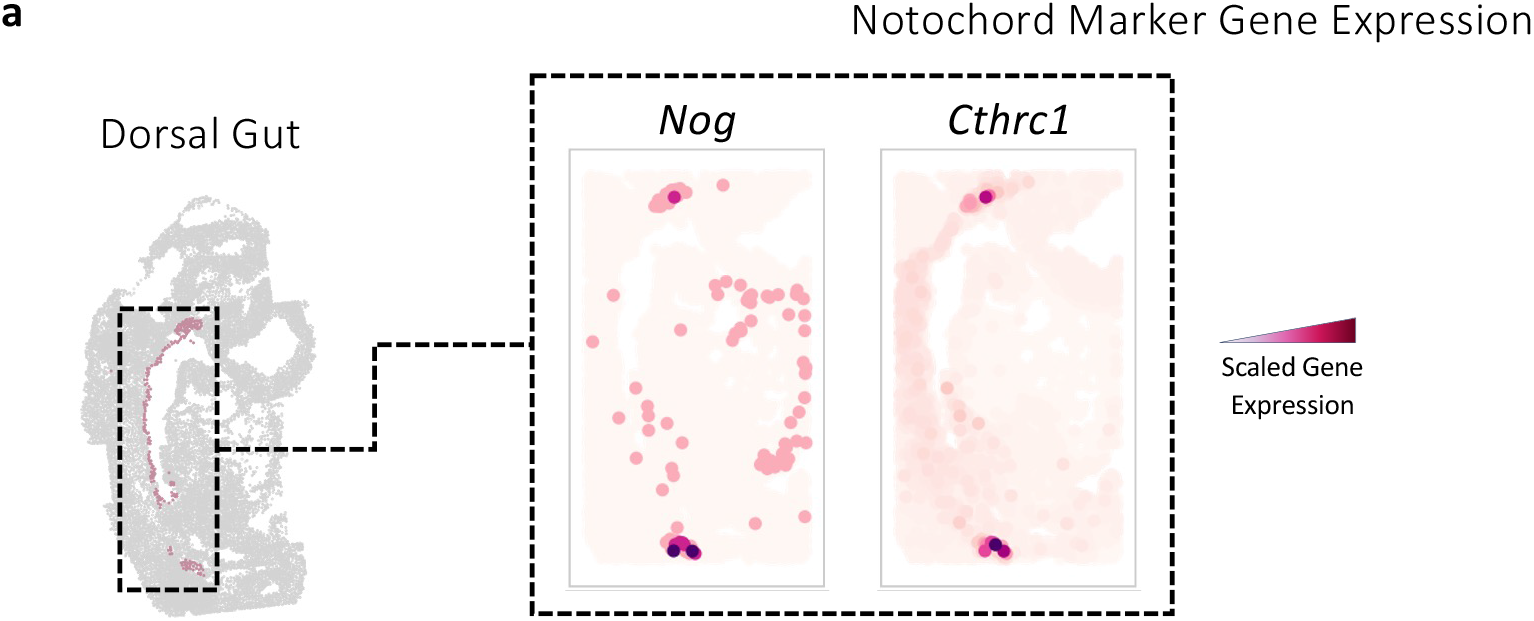
Notochord marker genes. **a,** Two known marker genes of the notochord locating this embryonic midline structure in the Dorsal Gut niche.

**Supplementary Fig. 8.**
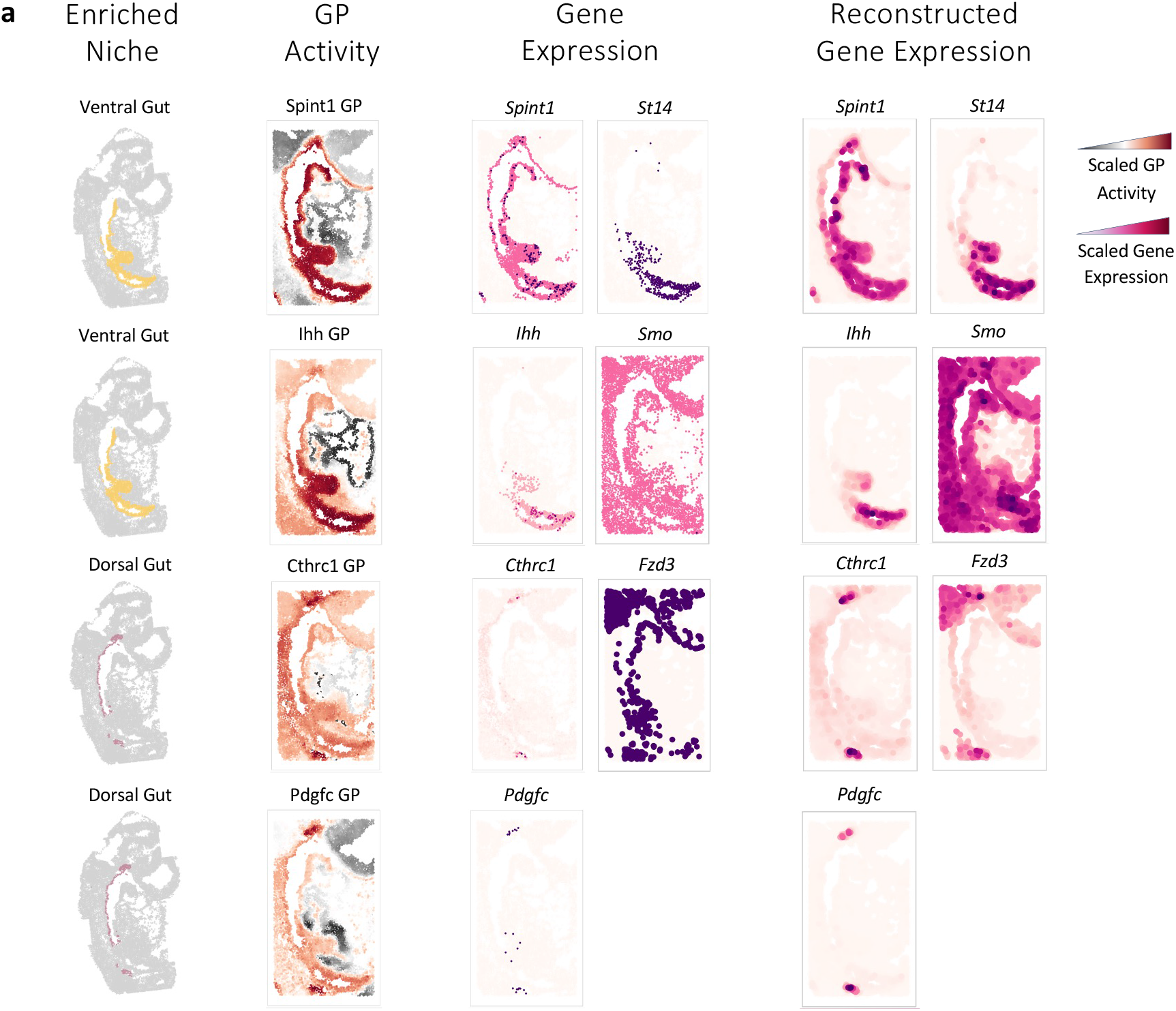
Enriched gene programs in the gut niches. **a,** The activity of characterizing GPs enriched in gut niches shows strong spatial correlation with the expression of important ligand and receptor genes of these GPs. Additionally, there is a high correlation with the model-reconstructed ligand and receptor expression, which closely resembles yet smoothens the original gene expression.

**Supplementary Fig. 9.**
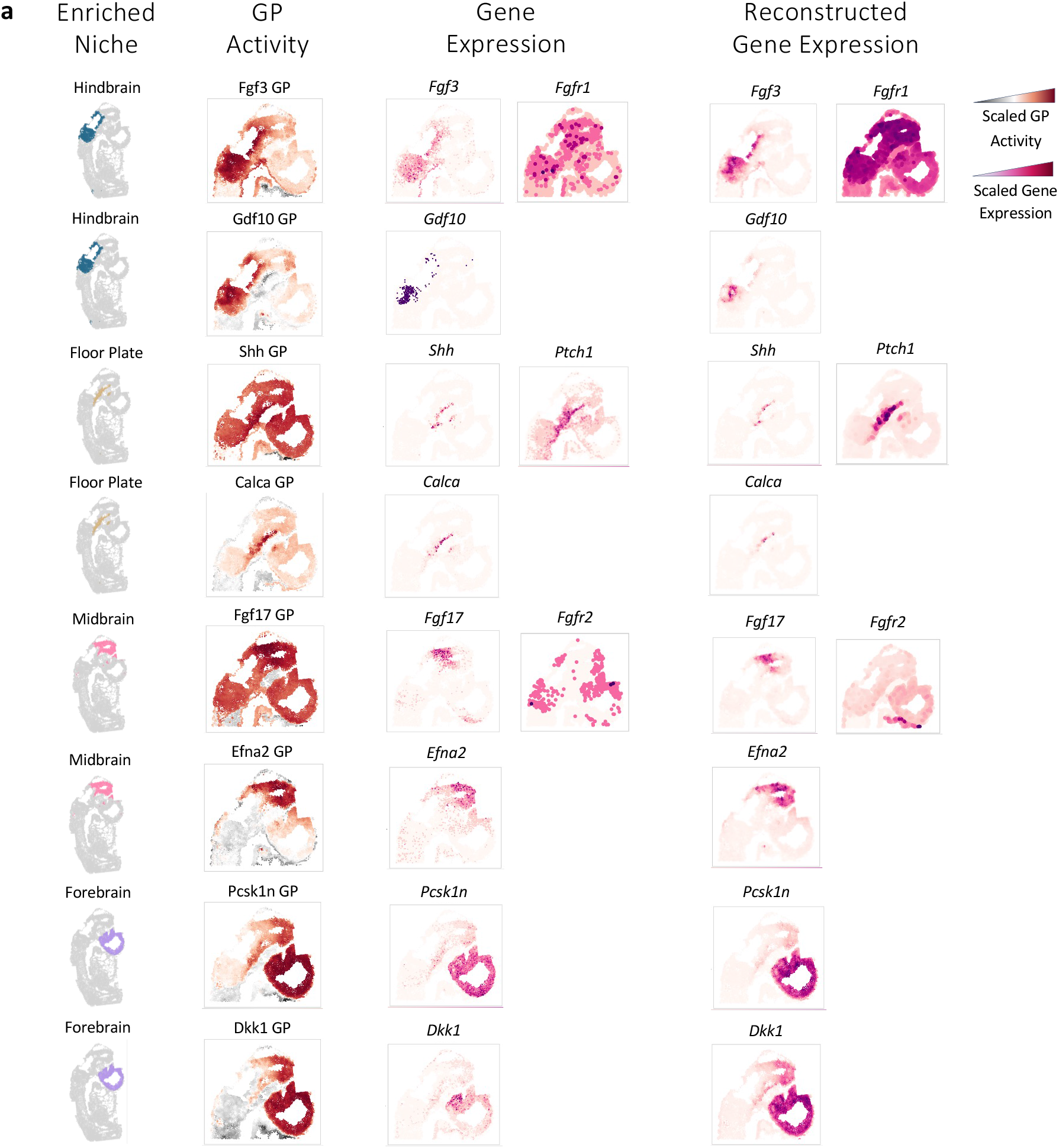
Enriched gene programs in the brain niches. **a,** The activity of characterizing GPs enriched in brain niches shows strong spatial correlation with the expression of important ligand and receptor genes of these GPs. Additionally, there is a high correlation with the model-reconstructed ligand and receptor expression, which closely resembles yet smoothens the original gene expression.

**Supplementary Fig. 10.**
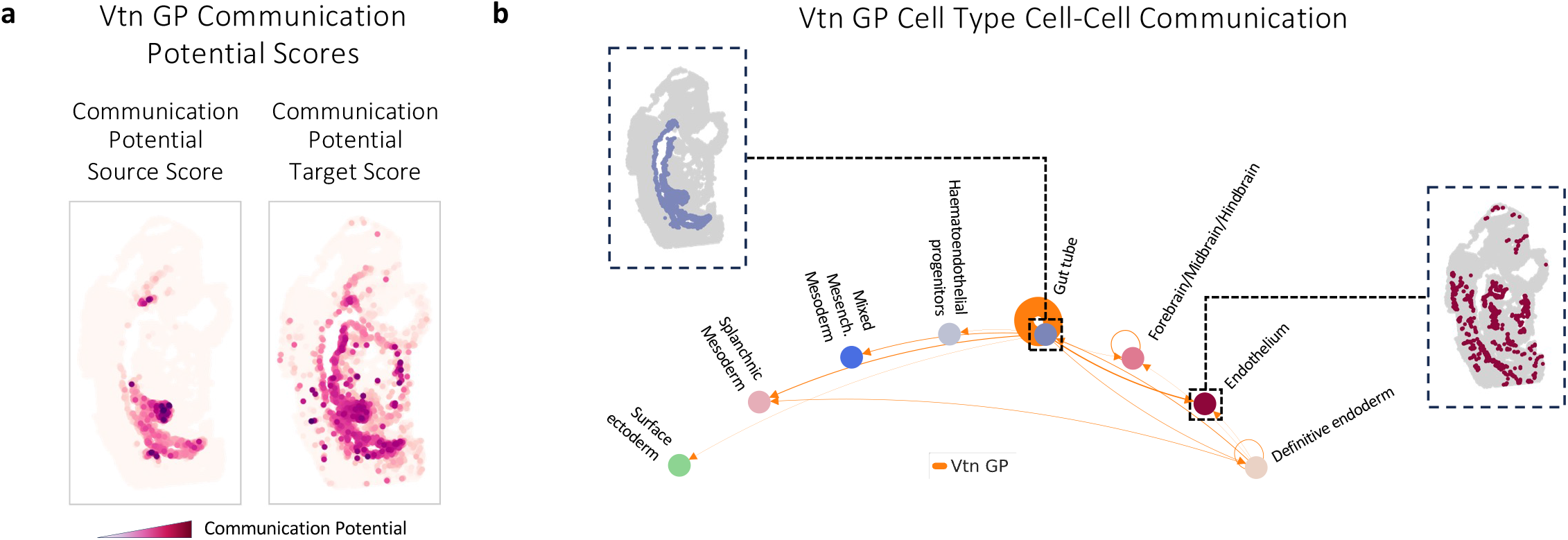
Cell-cell communication inference for the vitronectin gene program. **a,** Source- and target-specific cell-cell communication potential scores for the Vtn GP. **b,** Cell-pair communication strengths of the Vtn GP aggregated by cell types, highlighting interaction within gut tube cells and between the gut tube and endothelial cells.

**Supplementary Fig. 11.**
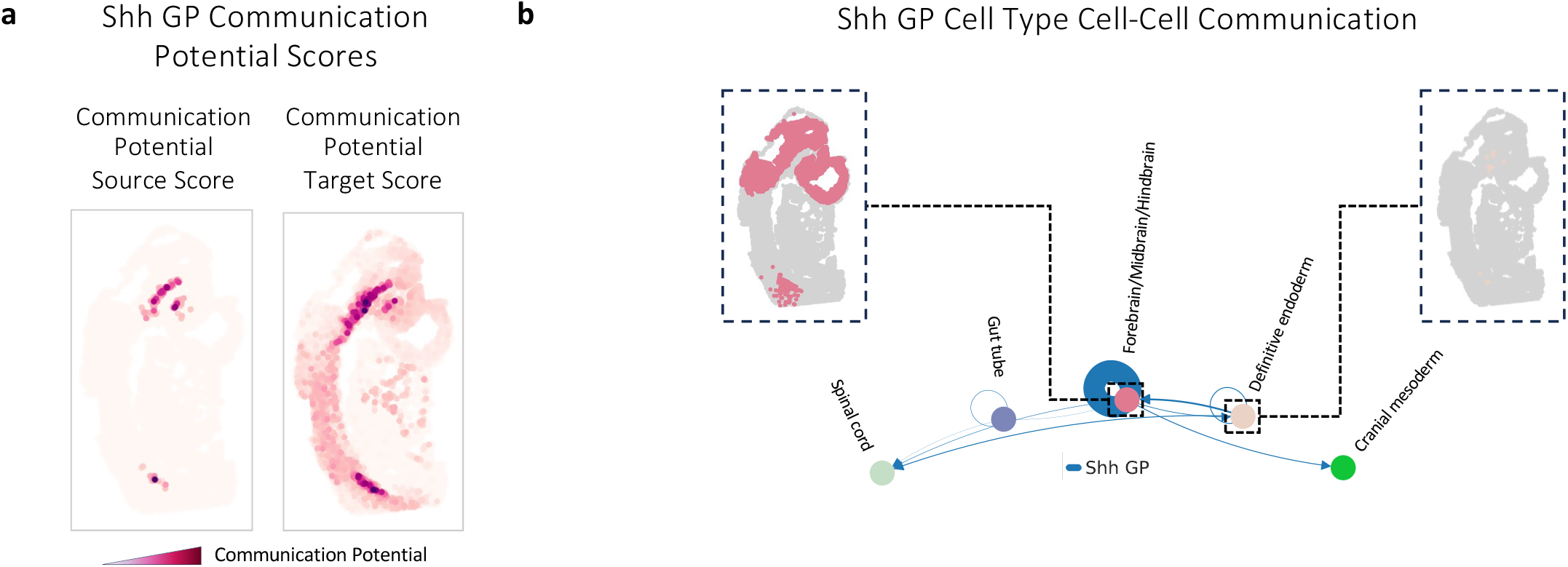
Cell-cell communication inference for the Sonic Hedgehog gene program. **a,** Source- and target-specific cell-cell communication potential scores for the Shh GP. **b,** Cell-pair communication strengths of the Shh GP aggregated by cell types, highlighting interaction within brain cells and between brain and definitive endoderm cells.

**Supplementary Fig. 12.**
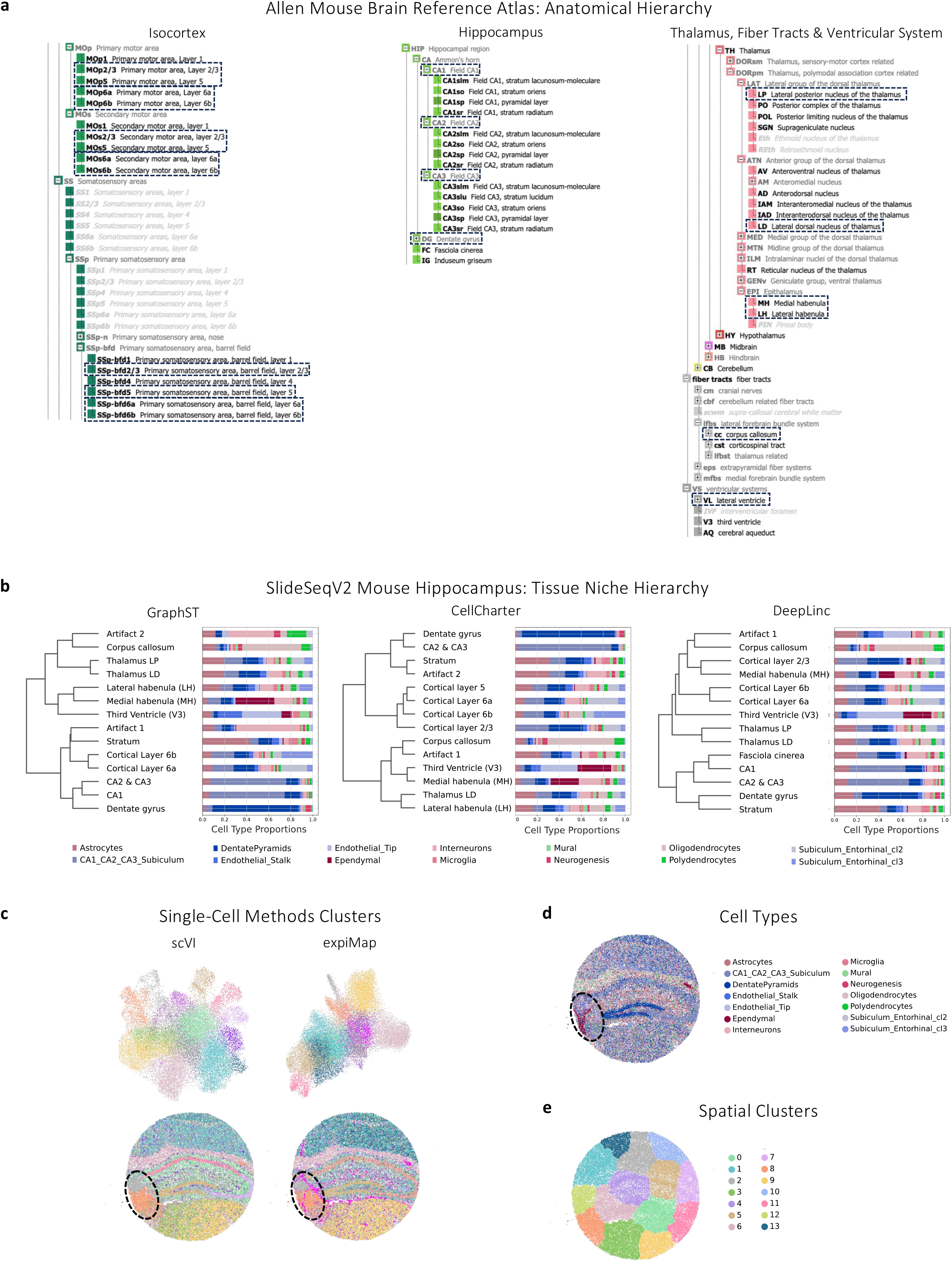
SlideSeqV2 mouse hippocampus benchmarking. **a,** An anatomical hierarchy of the mouse brain, obtained from the Allen Mouse Brain Reference Atlas. **b,** Dendrograms obtained through hierarchical clustering of the latent feature spaces of other spatial embedding methods, showing each method’s resulting tissue niche hierarchy. **c,** UMAP representations of the latent feature spaces retrieved from single-cell methods as well as the mouse hippocampus tissue, colored by the obtained clusters; highlighted is a region that shows clusters capturing spatial effects beyond cell types. **d,** The mouse hippocampus tissue, colored by original cell type annotations. **e,** The mouse hippocampus tissue, colored by clusters obtained from clustering of the spatial coordinates, illustrating spatially uniform, not biologically meaningful clusters.

**Supplementary Fig. 13.**
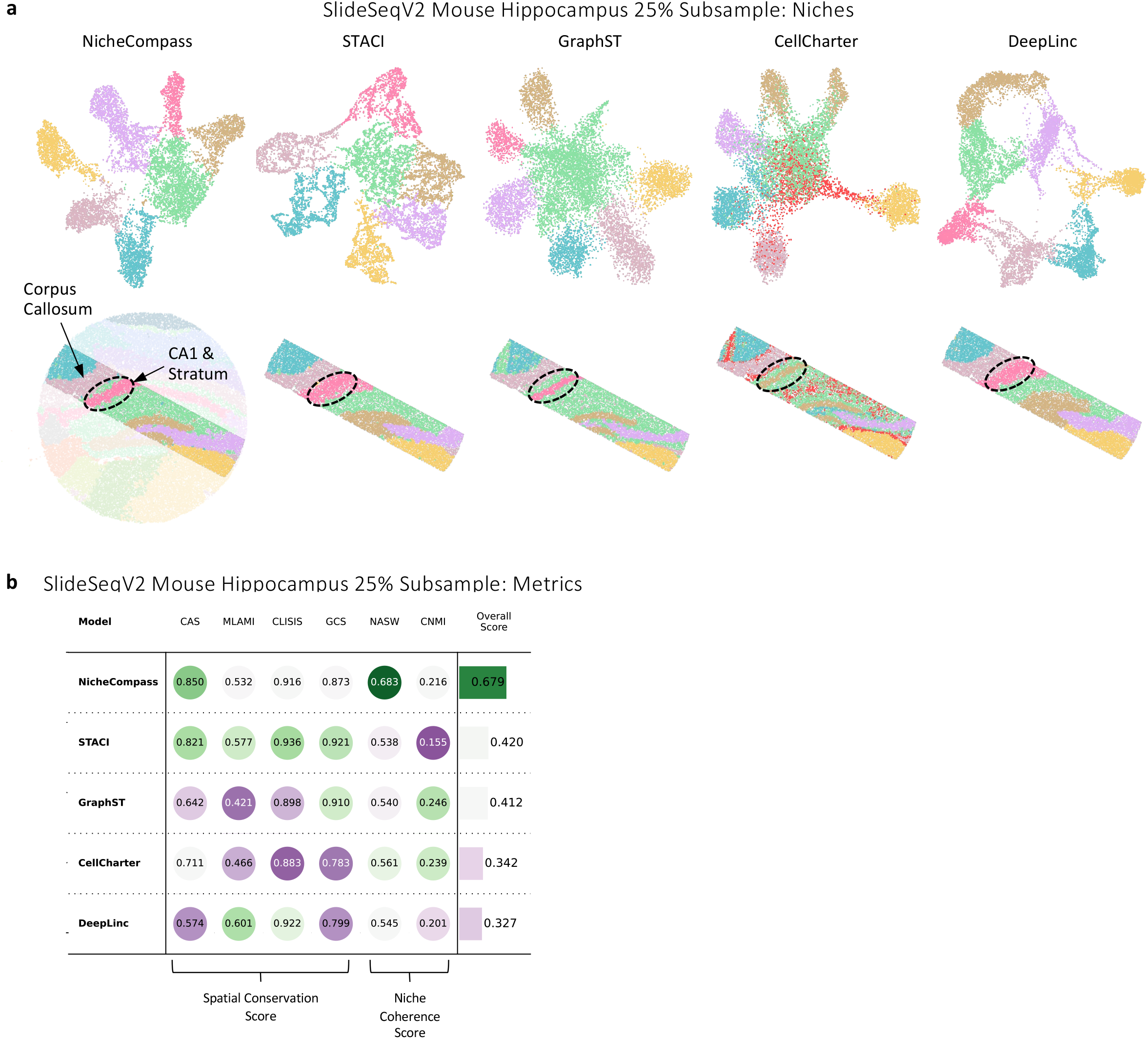
SlideSeqV2 mouse hippocampus 25% subsample benchmarking. **a,** UMAP representations of the latent feature spaces retrieved from NicheCompass and four competing methods as well as the 25% subsample of the mouse hippocampus tissue, colored by niches identified through clustering of the latent feature spaces. The number and color of clusters match between methods. The tissue below NicheCompass is overlaid with the one from the full dataset, showing high consistency of the obtained niches **b,** Six benchmarking metrics across two categories are normalized and aggregated into an overall score to quantify the performance of each method on this dataset.

**Supplementary Fig. 14.**
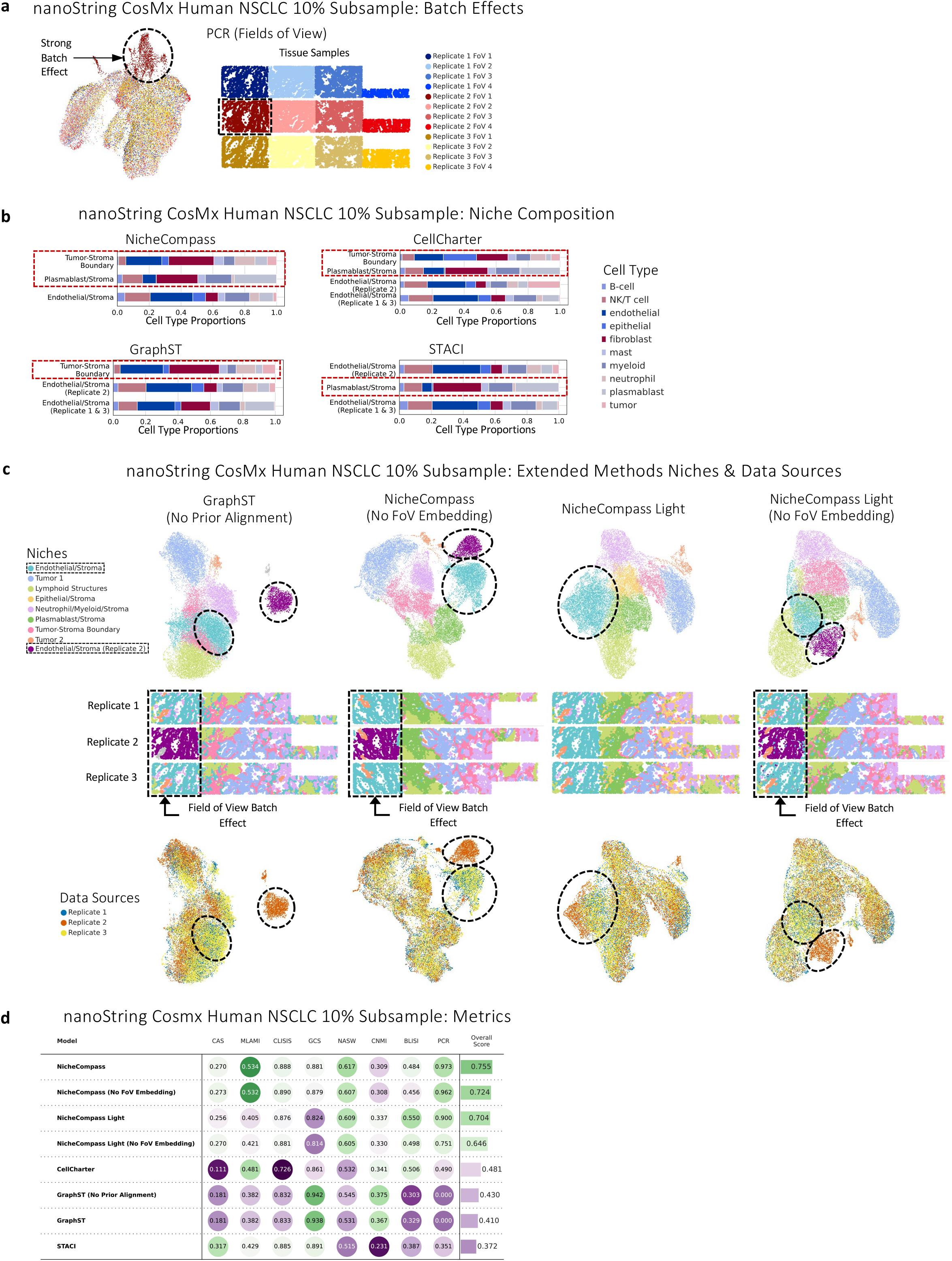
nanoString CosMx human NSCLC 10% subsample benchmarking. **a,** UMAP representation after applying PCR to the raw gene expression of the three replicates, showing the presence of strong batch effects in the first field of view of the second tissue sample. **b,** Niche composition of important niches for each method. NicheCompass identified a Tumor-Stroma Boundary niche and could differentiate between Stroma enriched by endothelial cells and Stroma enriched by plasmablast cells. In the case of STACI, both Endothelial-enriched Stroma niches showed very similar cell type composition, suggesting they should be unified into one niche. In the case of GraphST, a similar issue was observed, except for the misallocation of plasmablast cells to the Endothelial-enriched Stroma. **c**, A comparison of the integration performance of further method variants. Illustrated are the UMAP representations of the learned latent feature spaces of the models and the tissue, colored by annotated niches. Niches in the first field of view are highlighted, showing differences in batch effect removal capabilities. UMAP representations colored by data source further emphasize differences in batch effect removal for the first field of view. FoV: Field of View. GraphST (No Prior Alignment) was trained without alignment through PASTE before model training, the use of which is recommended by the authors. **d,** Metrics for the training runs from **c** and Fig. 3d. The overall score is computed by aggregating normalized individual metric scores into the two categories spatial conservation and niche coherence, followed by equal weighting of these categories. NicheCompass Light is a variation of our model that uses graph convolutional layers instead of dynamic graph attention layers.

**Supplementary Fig. 15.**
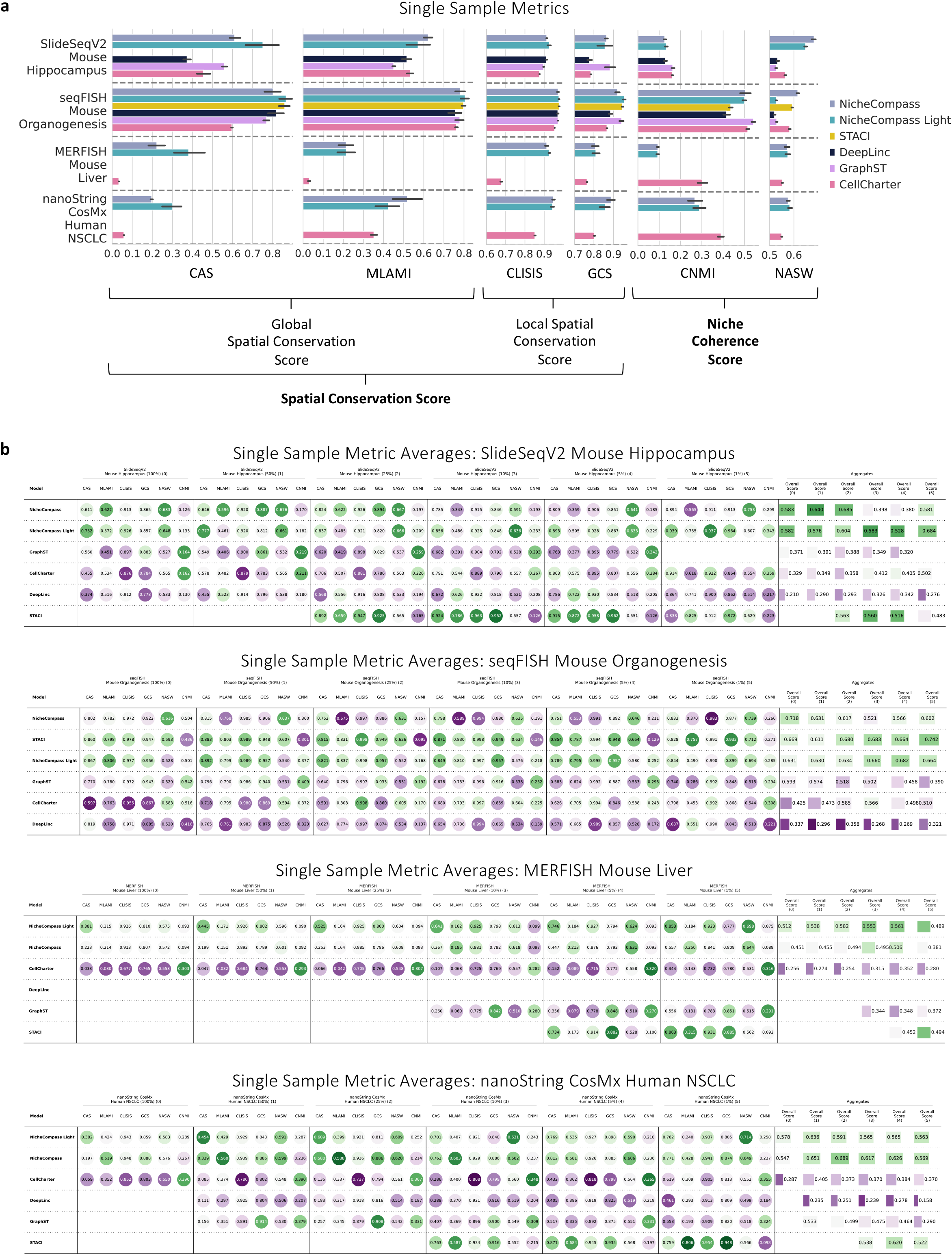
Extended single sample benchmarking results. **a,** Individual metrics results from single sample benchmarking across diverse datasets. The bars display the mean across eight training runs for each method and dataset with varying numbers of neighbors. The error bars display the 95% confidence interval. **b,** Mean metrics across eight training runs for different subsample sizes of all datasets. Missing entries are either due to memory overflow or failure of the model to converge.

**Supplementary Fig. 16.**
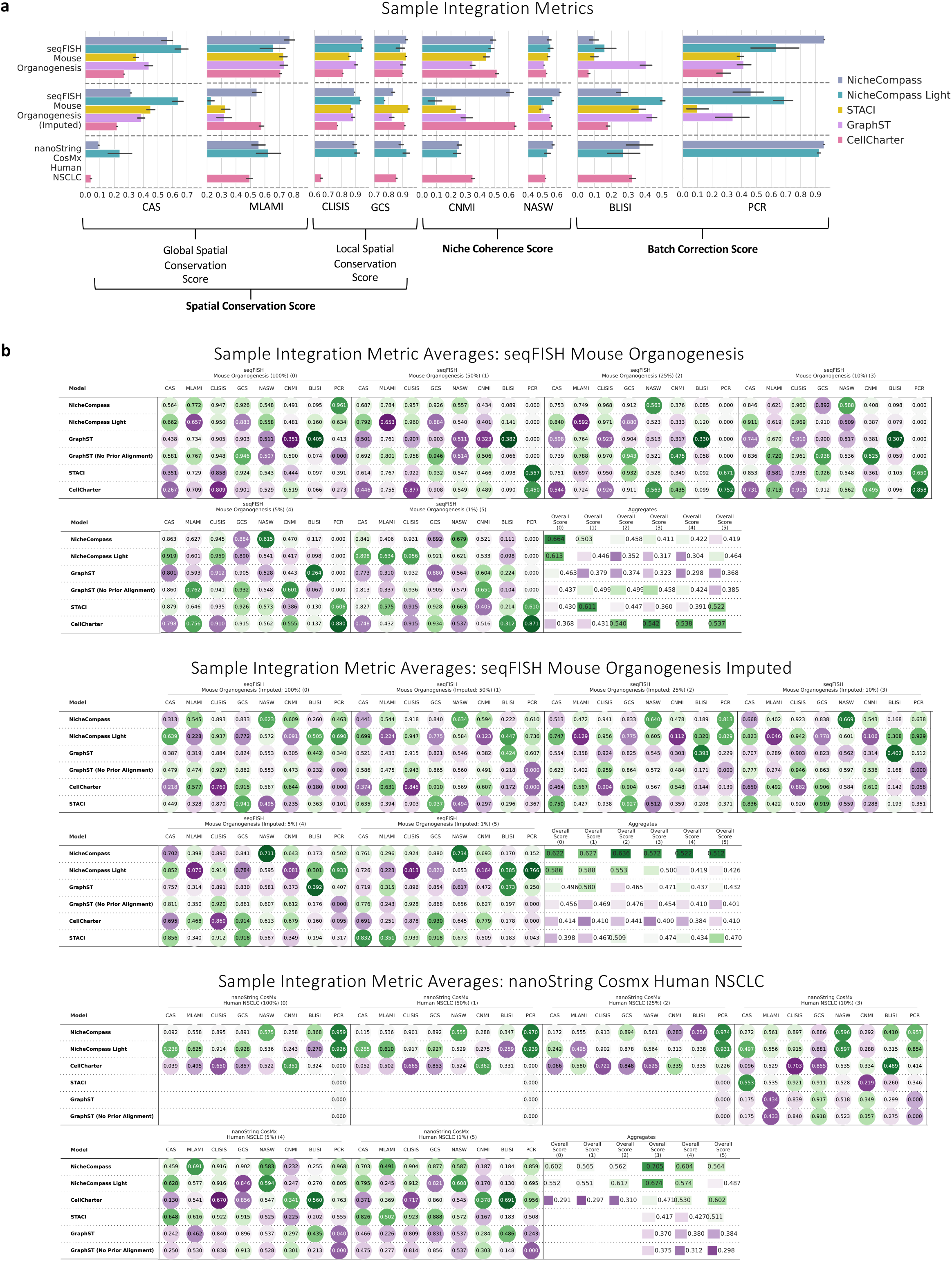
Extended sample integration benchmarking results. **a,** Individual metrics results from sample integration benchmarking across diverse datasets. The bars display the mean across eight training runs for each method and dataset with varying numbers of neighbors. The error bars display the 95% confidence interval. **b,** Mean metrics across eight training runs for different subsample sizes of all datasets. Missing entries are either due to memory overflow or failure of the model to converge.

**Supplementary Fig. 17.**
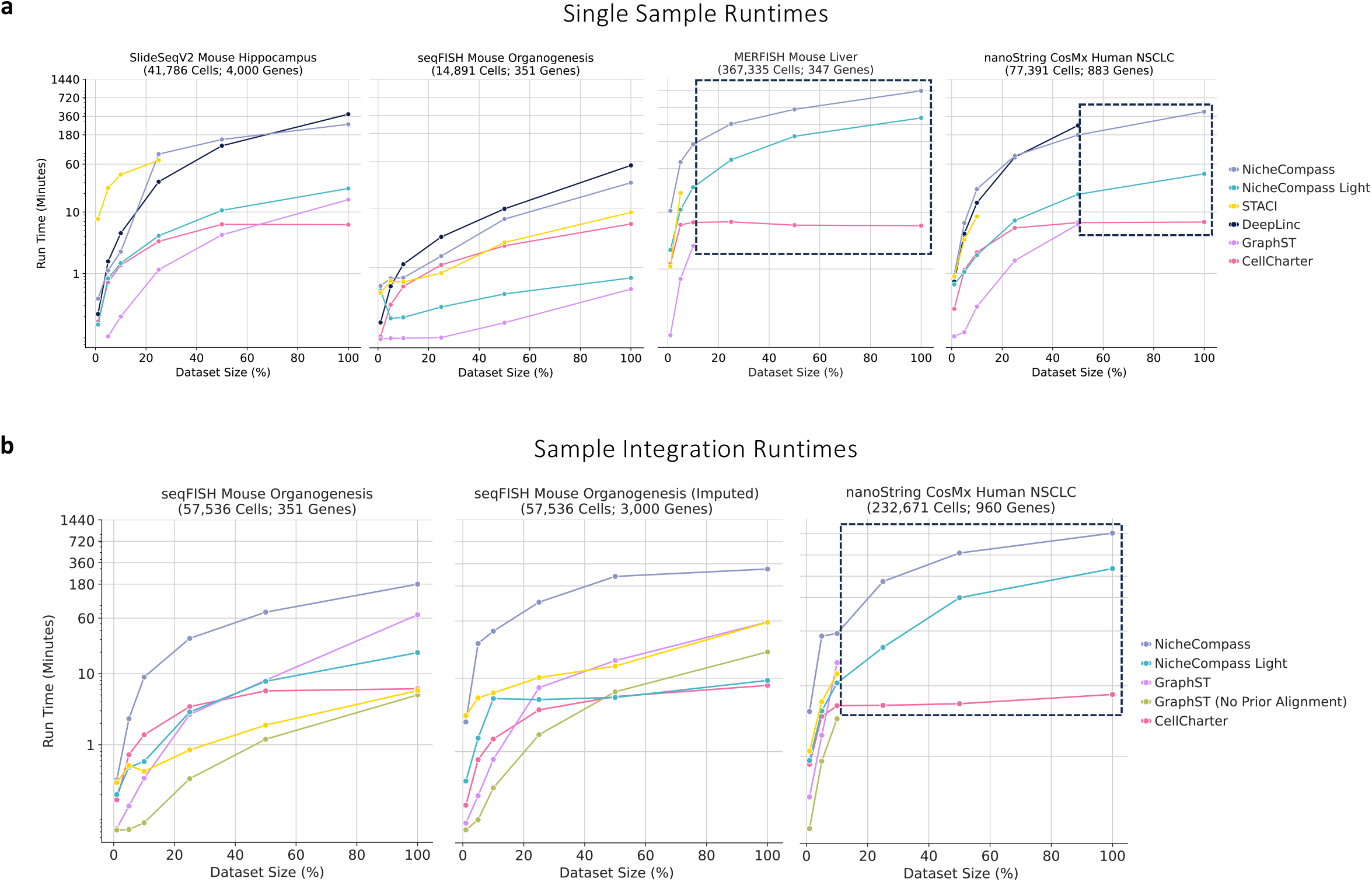
Benchmarking runtimes. **a,** Mean runtimes across eight training runs for all models trained during single sample benchmarking. Highlighted is the regime where only NicheCompass and CellCharter ran successfully. **b,** Same as **a** but for models trained during sample integration benchmarking.

**Supplementary Fig. 18.**
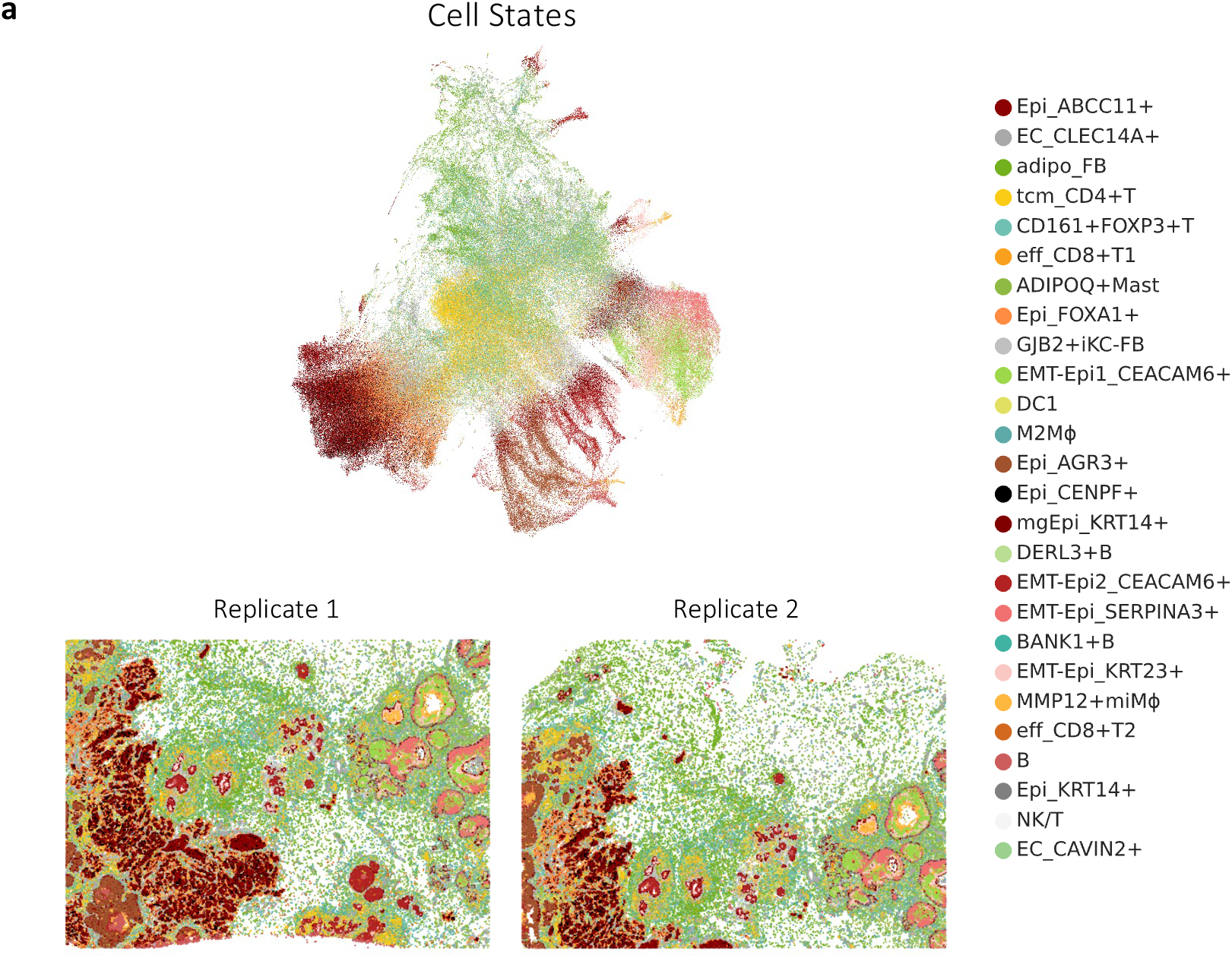
Xenium human breast cancer cell states. **a,** UMAP representation and the two tissue replicates, annotated by cell states.

**Supplementary Fig. 19.**
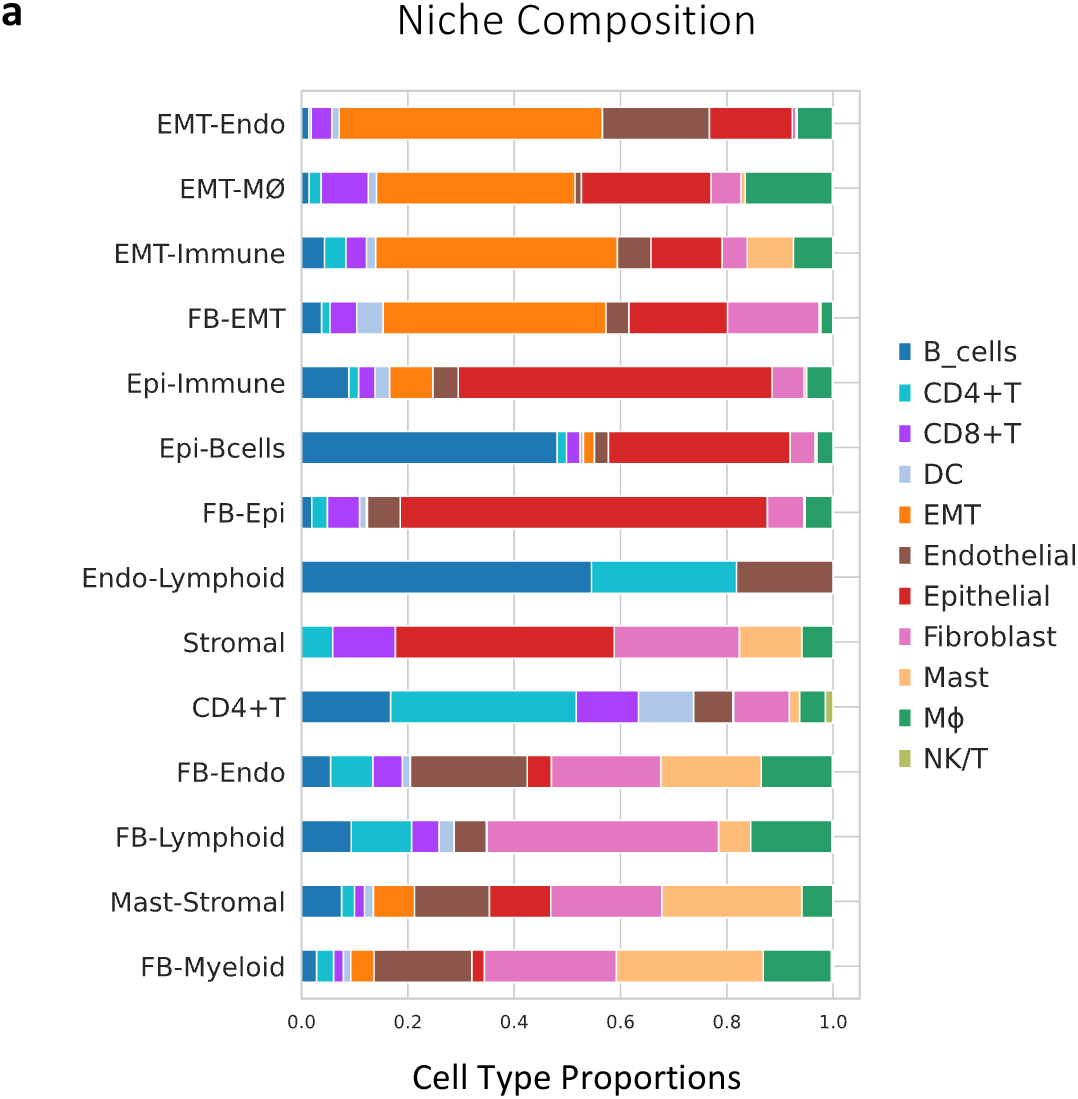
Xenium human breast cancer niche composition. **a,** Cell type proportions of all niches.

**Supplementary Fig. 20.**
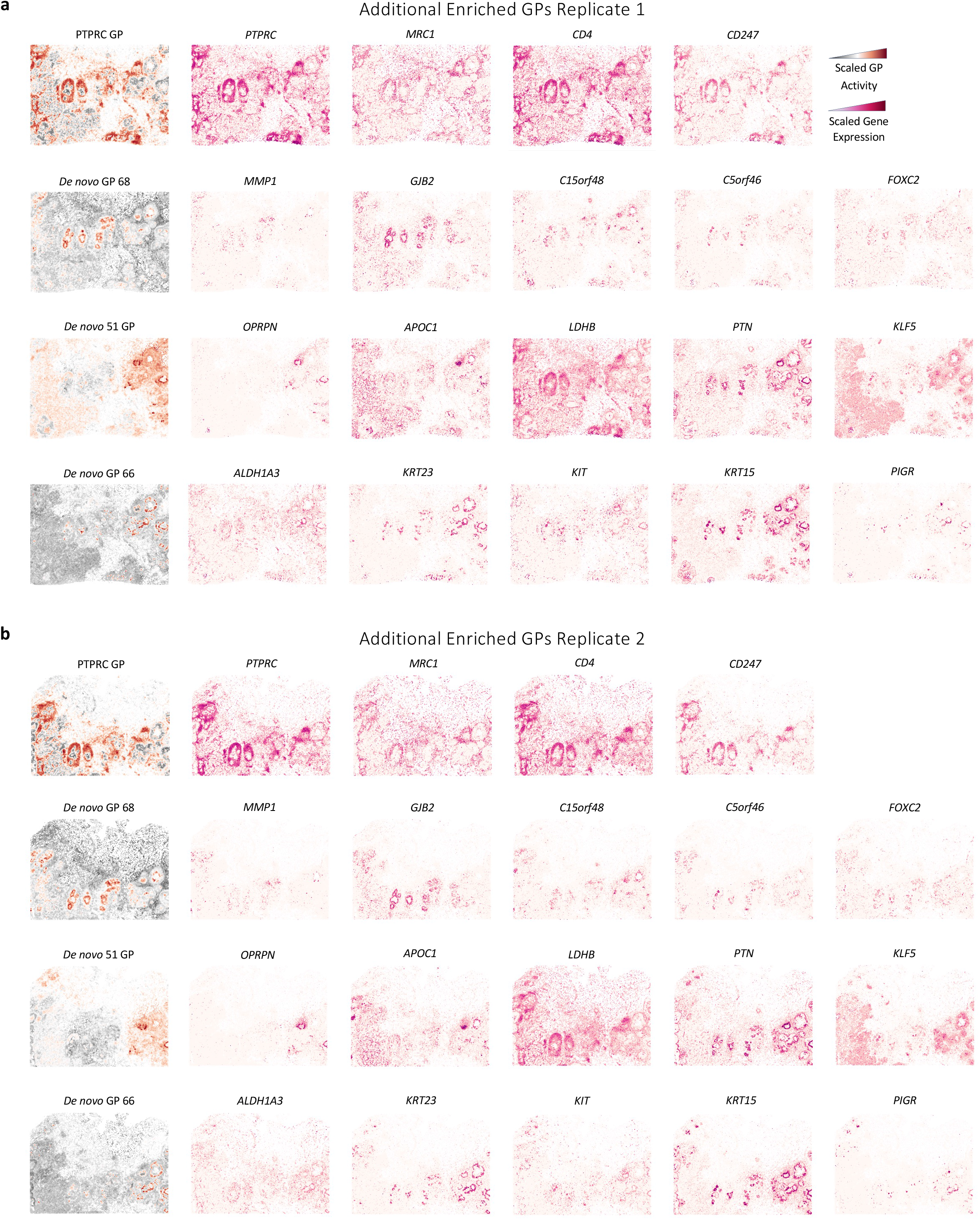
Xenium human breast cancer additional enriched gene programs. **a,** GP activity and gene expression of the most important genes of additional enriched GPs in niches of the Xenium human breast cancer dataset.

**Supplementary Fig. 21.**
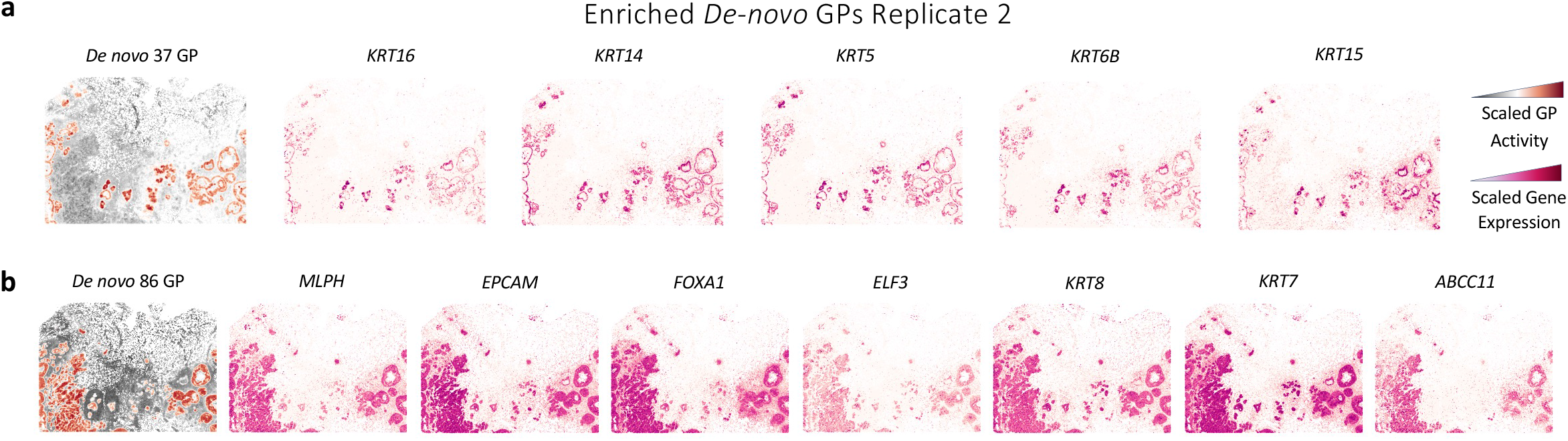
Xenium human breast cancer enriched *de novo* gene programs in replicate 2. **a,** GP activity and gene expression of the most important genes of enriched *de novo* GP 37 in replicate 2 of the Xenium human breast cancer dataset. **b,** Same as **a** but for *de novo* GP 86.

**Supplementary Fig. 22.**
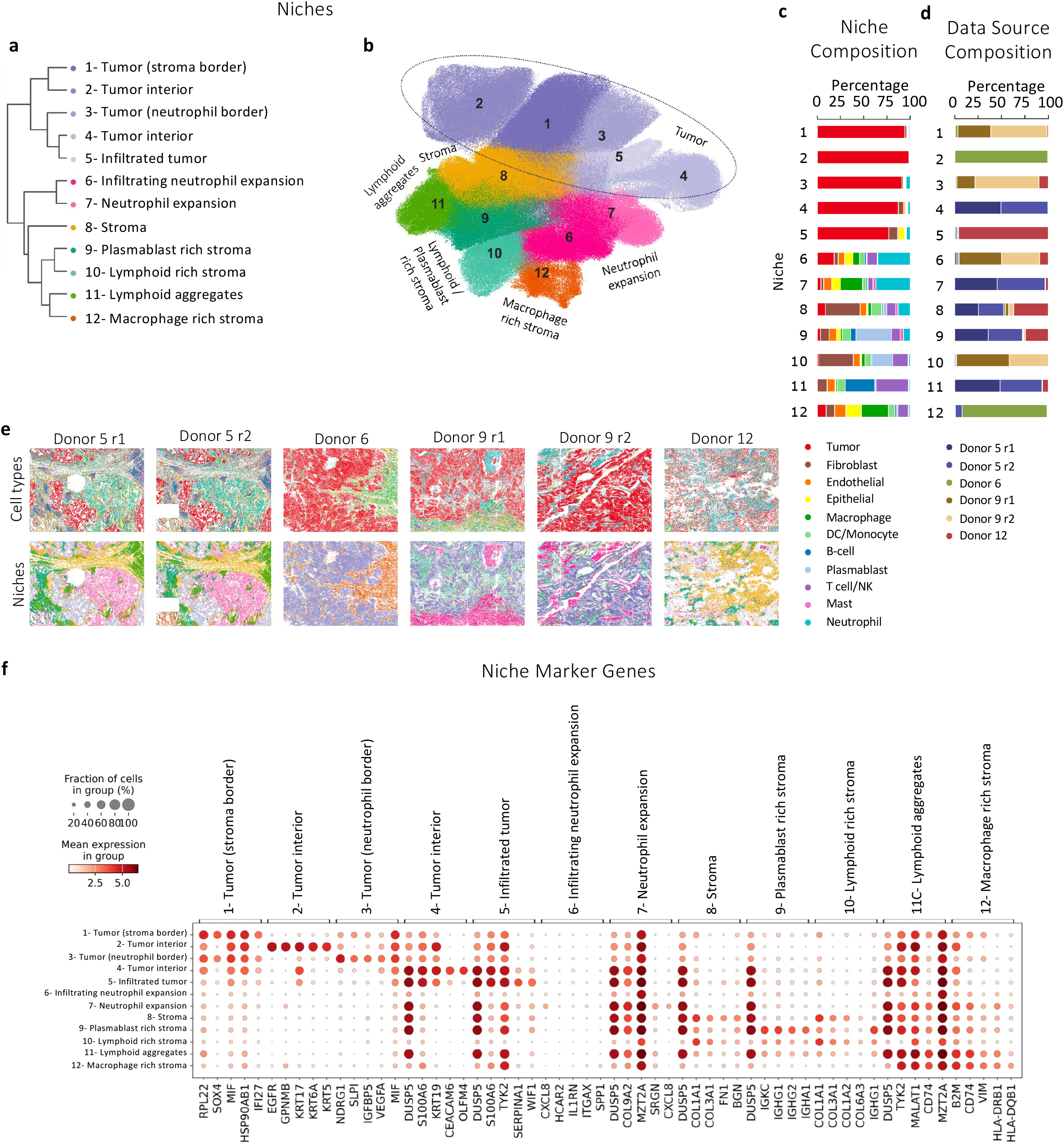
Analysis of inter-tumoral heterogeneity using NicheCompass. **a,** A dendrogram computed based on average GP activities, showing a hierarchy of niches. **b,** UMAP representation of the reference atlas, colored by niches identified with NicheCompass. **c, d,** Barplots representing the cellular composition (**c**) and donor composition (**d**) of the identified niches. **e,** Spatial visualization of the six tissue sections included in the reference, colored by cell type and niche. **f,** Dotplot showing the five most differential genes expressed in each niche compared to the rest. The dot size represents the fractions of cells in a niche with expression higher than 0, while the dot color represents the mean expression level within the expressing cells.

**Supplementary Fig. 23.**
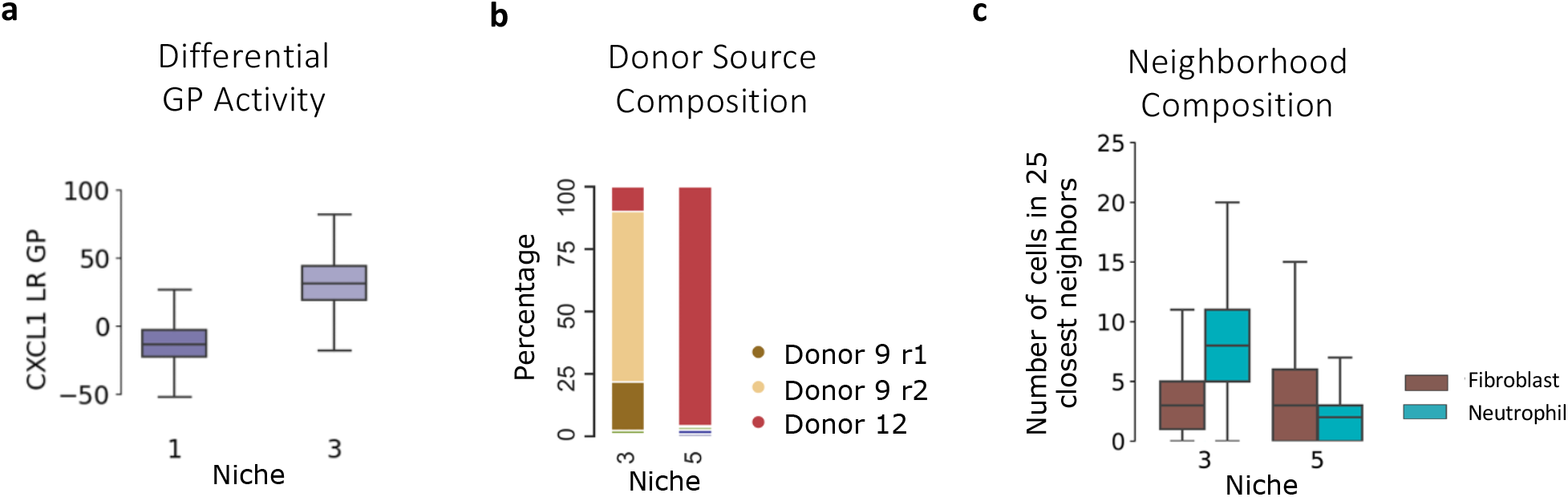
Identification of tumor niches interacting with neutrophils using NicheCompass. **a,** Boxplot representing the CXCL1 ligand-receptor GP activity distribution in the cells from niche 1 and niche 3 in donor 9. Boxplot elements are defined as: center line, median; box limits, upper and lower quartiles; whiskers, 1.5x interquartile range. **b,** Barplots representing the donor composition of niche 3 and 5. **c**, Neighborhood composition in tumor niche 3 and 5 in donor 12. A boxplot per niche and neighboring cell type represents the distribution among all cells in a niche of the number of cells of a given cell type among the 25 physically closest cells. Boxplot elements are defined as in **a**. For clarity, only cell types composing on average more than 5% and less than 60% of the neighborhood of any niche are shown.

**Supplementary Fig. 24.**
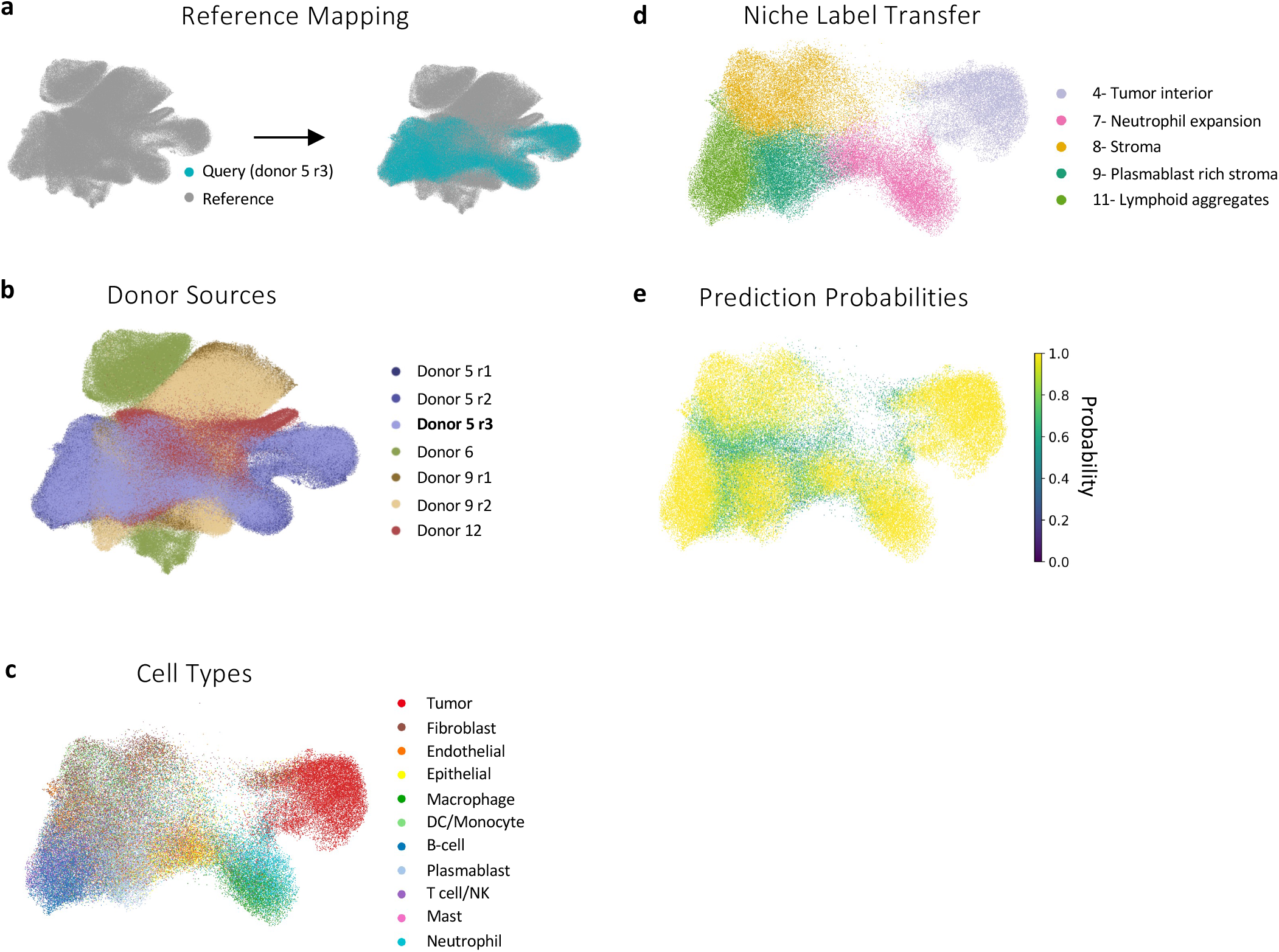
NicheCompass correctly integrates unseen biological replicates with shared biology into a pre-built reference atlas. a,b, UMAP representation of reference and query cells in the NicheCompass latent GP space, obtained by mapping query cells onto the reference latent space with fine-tuning, colored by mapping entity (**a**) and sample donor and replicate (**b**). **c,d,e,** UMAP representations of the query cells in the NicheCompass latent GP space, colored by cell type (**c**), niche label (**d**) as predicted by a kNN classifier trained on the reference, and prediction probability of the classifier (**e**).

**Supplementary Fig. 25.**
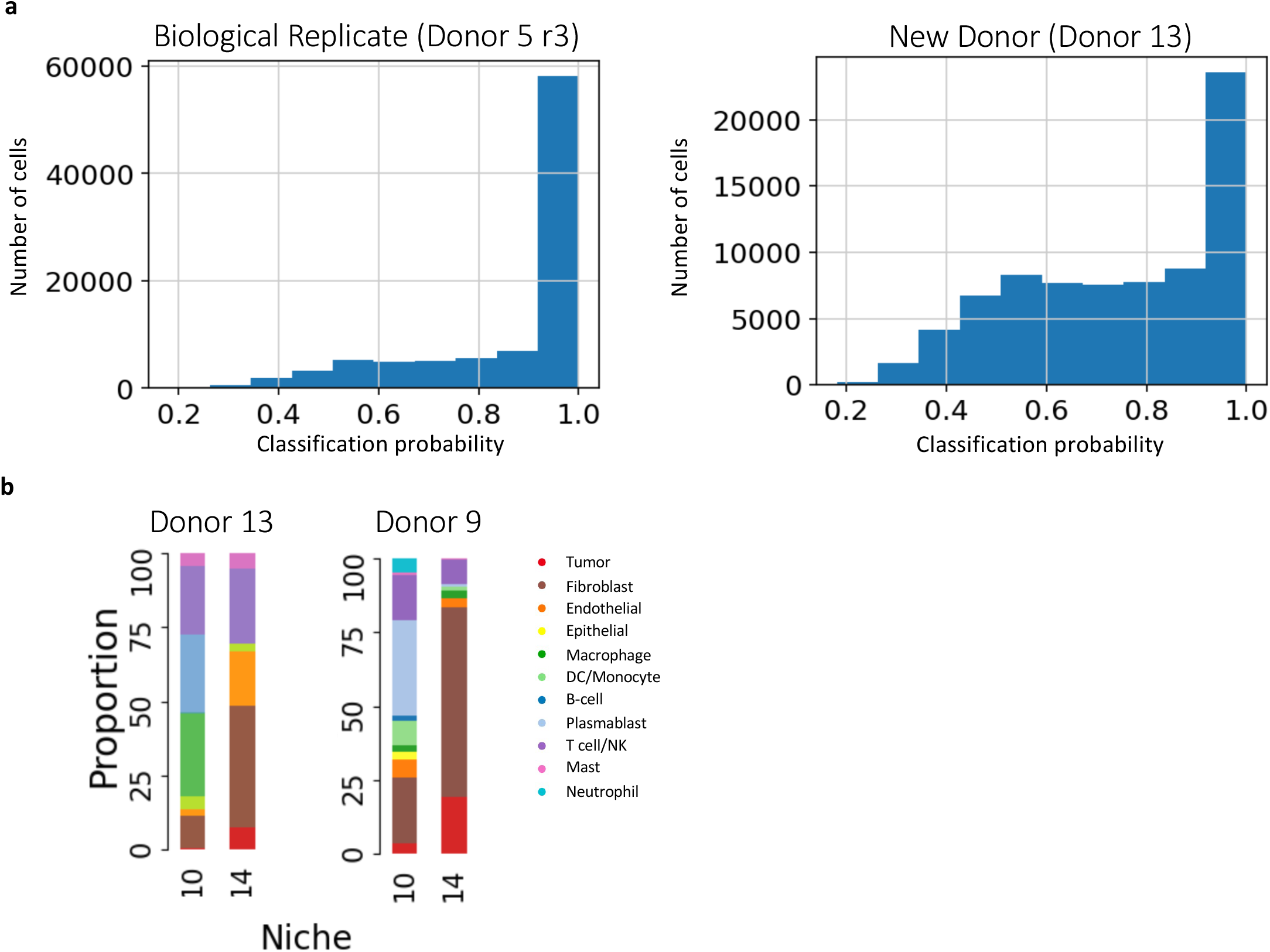
Label transfer from reference to query sample distinguishes seen from unseen niches. **a,** Classification probability distribution of niches, obtained from a kNN classifier trained on the reference atlas and applied to an unseen biological replicate (left) and an unseen donor (right). **b,** Barplots representing the cellular composition of the infiltrating stromal niches 10 and 14, which are shared between the new donor 13 (left) and the reference donor 9 (right).

**Supplementary Fig. 26.**
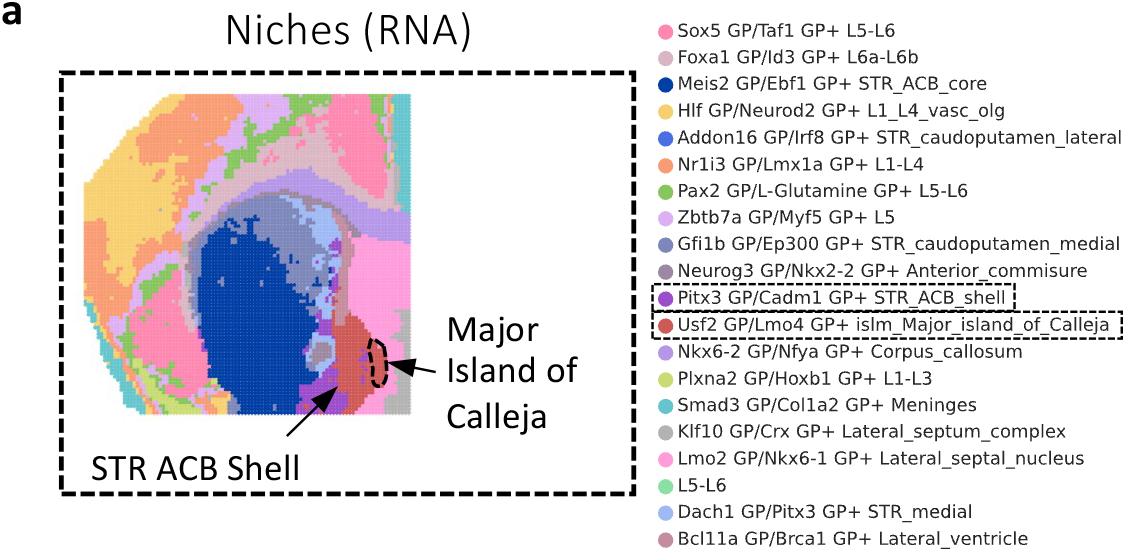
Spatial ATAC-RNA seq mouse brain NicheCompass niches (RNA modality). **a,** The mouse brain tissue, colored by niches identified by a NicheCompass model trained on just the RNA modality; highlighted is the failure of the model to separate the Major Island of Calleja and the STR ACB Shell into separate niches.

**Supplementary Fig. 27.**
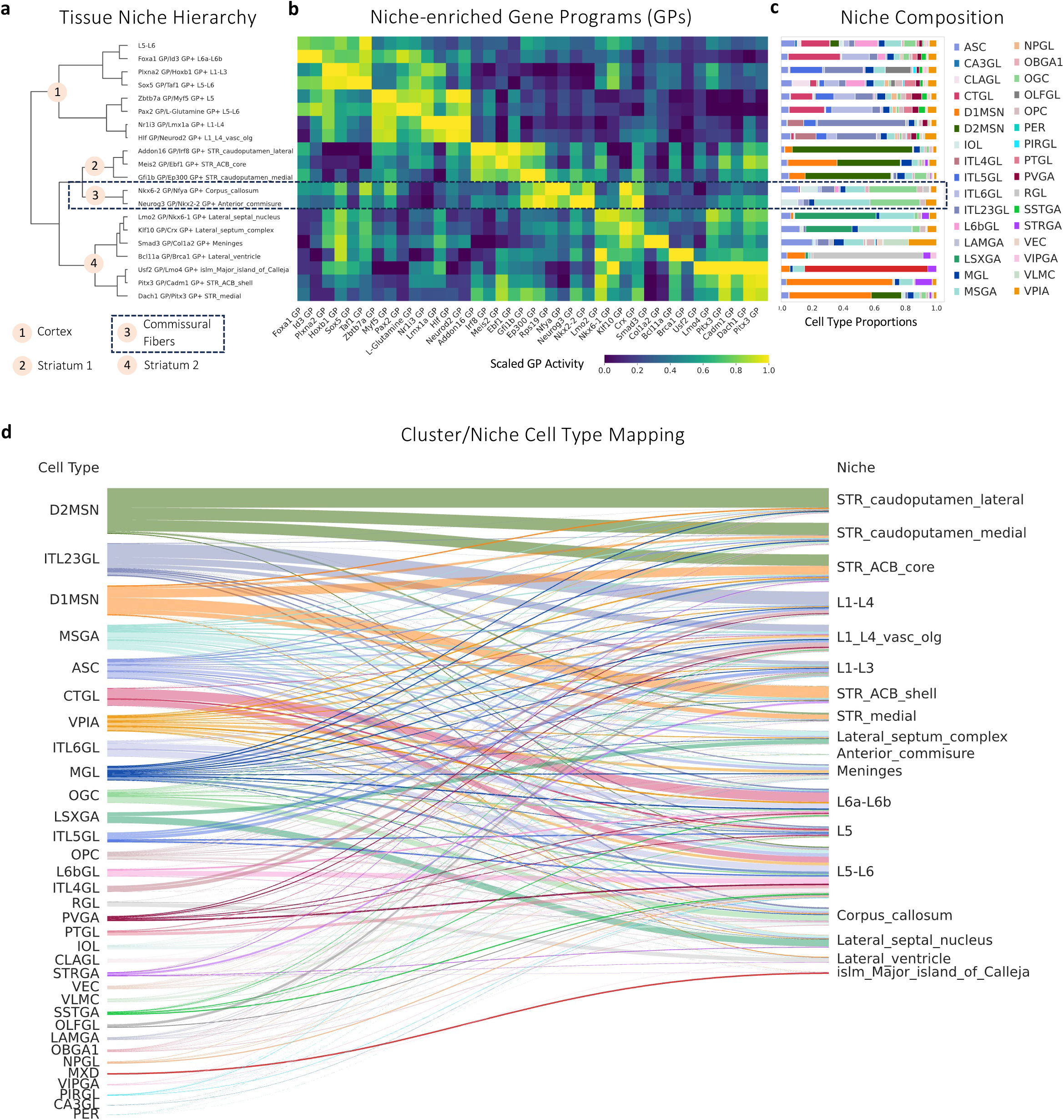
Spatial ATAC-RNA seq mouse brain NicheCompass tissue niche organization. **a,** A dendrogram computed based on average GP activities with the annotated niche labels, showing a functional niche hierarchy of higher-order functional components. **b,** A heatmap containing for each niche from **a** the normalized activity of two characterizing GPs. A gradient along the clustering obtained from **a** is visible. **c,** Cell type proportions for each niche from **a**. **d**, Mapping of the original cell type annotations to the clusters obtained from clustering the NicheCompass latent GP space.

**Supplementary Fig. 28.**
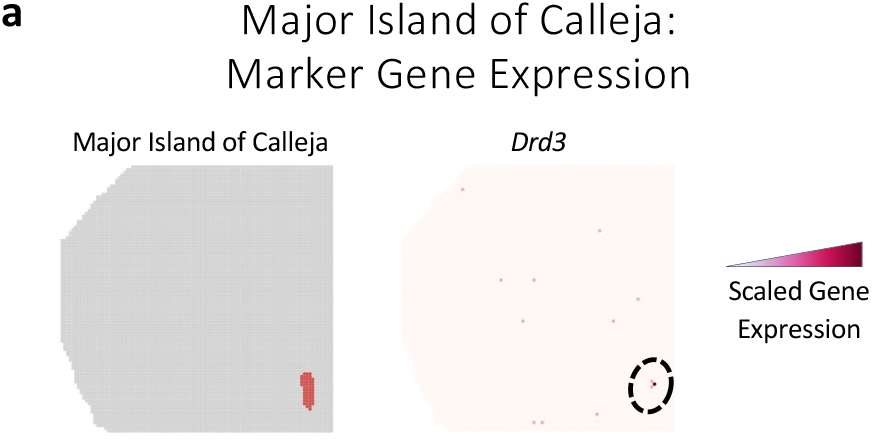
Major Island of Calleja marker genes. **a,** The Major Island of Calleja niche and gene expression of the dopamine D3 receptor *Drd3* gene, which has been described as a marker gene of the Major Island of Calleja.

**Supplementary Fig. 29.**
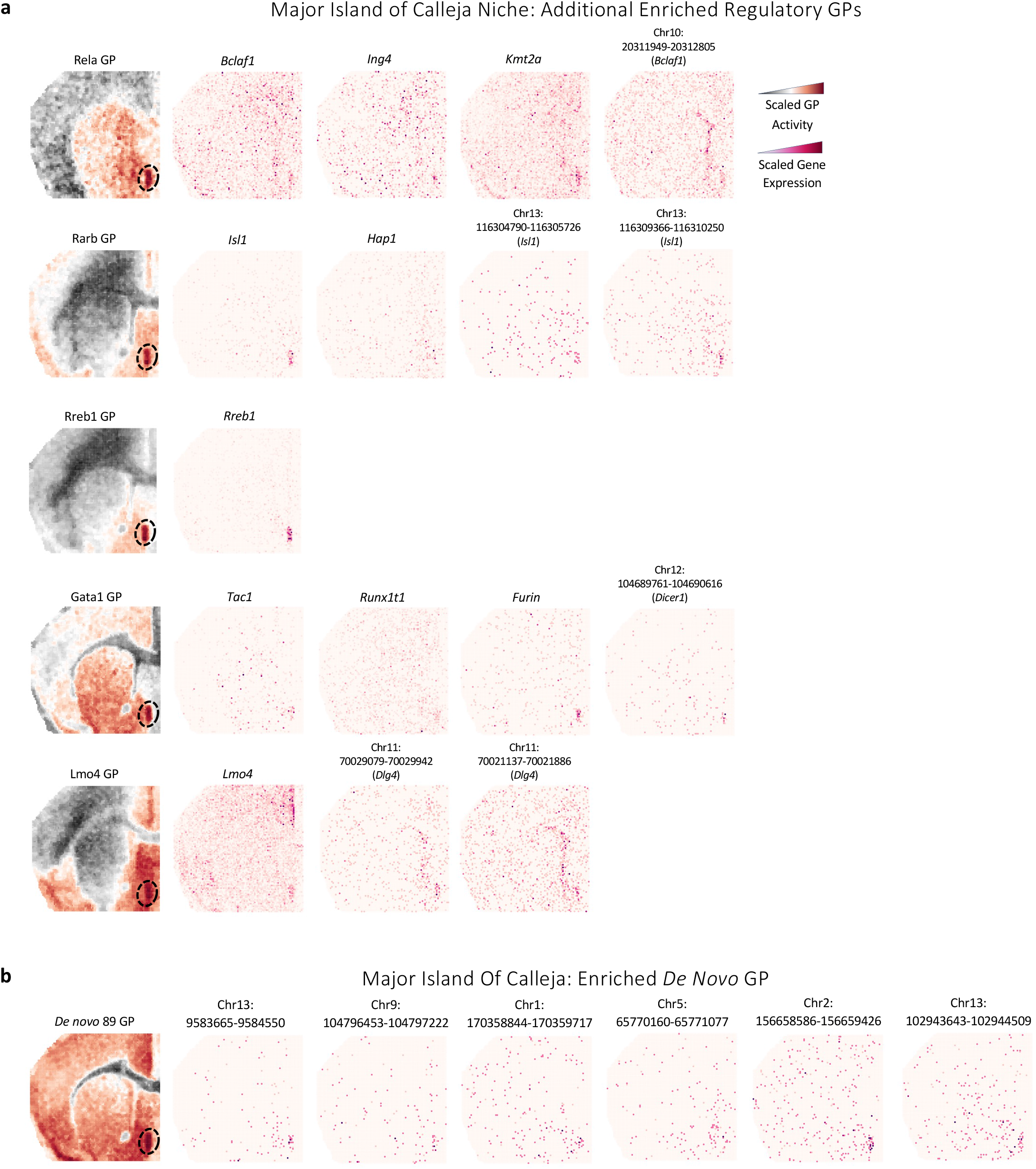
Additional enriched GPs in the Major Island of Calleja niche. **a,** The GP activity and gene expression of important GP genes of additional enriched regulatory GPs in the Major Island of Calleja niche. **b,** Same as **a** but showing an enriched *de novo* GP.

**Supplementary Fig. 30.**
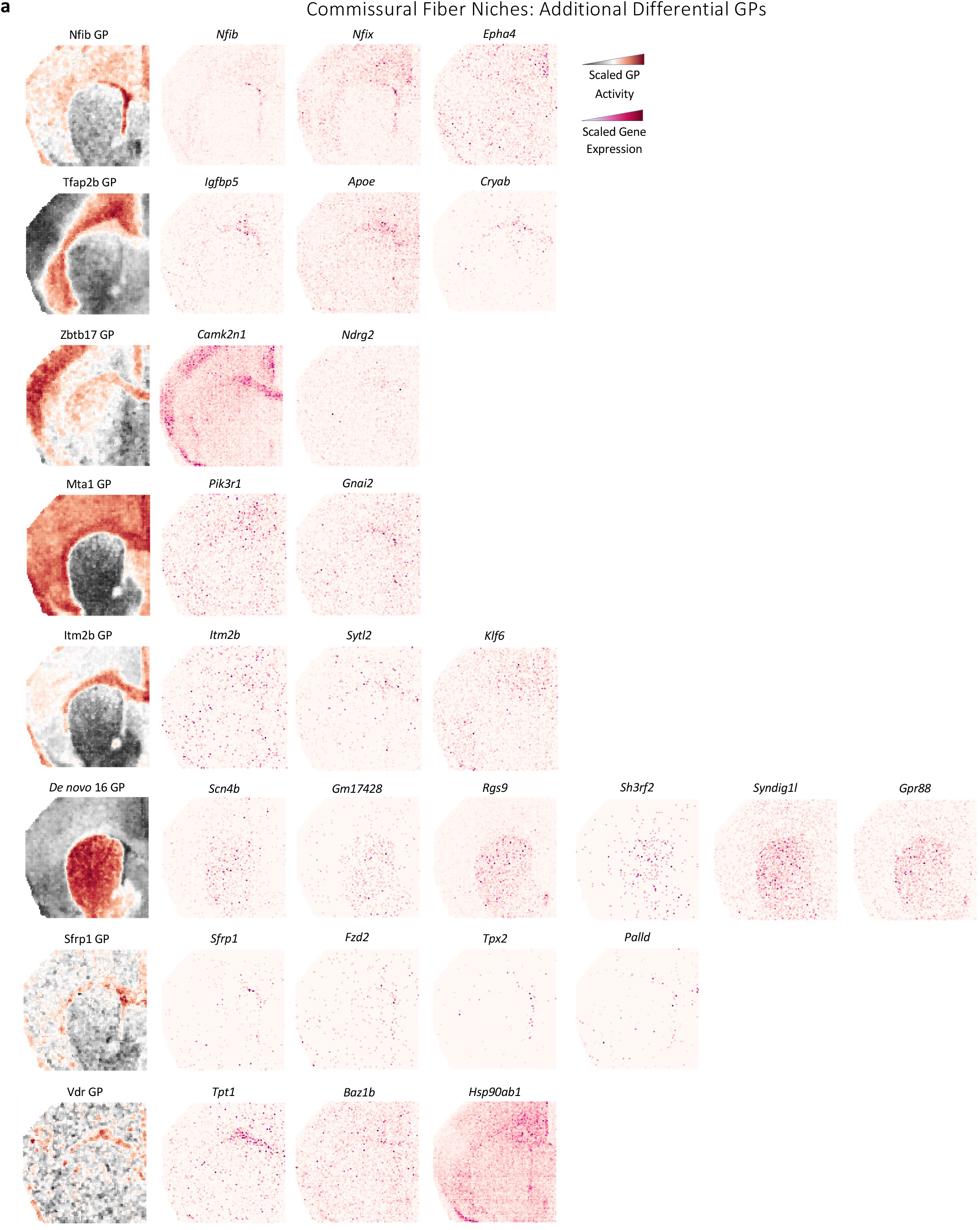
Additional differential GPs in the Commissural Fiber niches. **a,** The GP activity and gene expression of important GP genes of additional GPs differentially active in the Commissural Fiber niches Corpus Callosum and Anterior Commissure.

**Supplementary Fig. 31.**
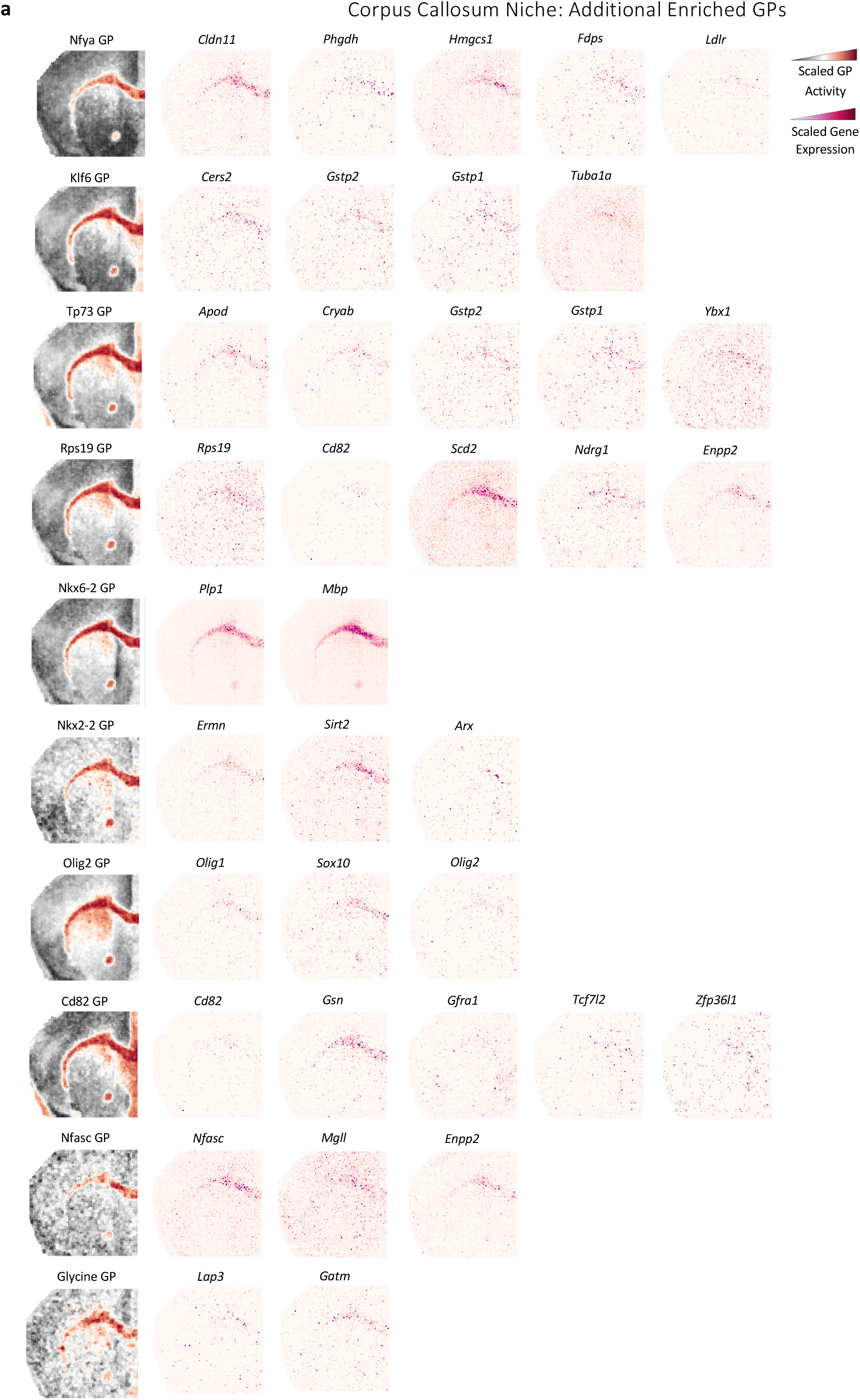
Additional enriched GPs in the Corpus Callosum niche. **a,** The GP activity and gene expression of important GP genes of additional enriched GPs in the Corpus Callosum niche.

**Supplementary Fig. 32.**
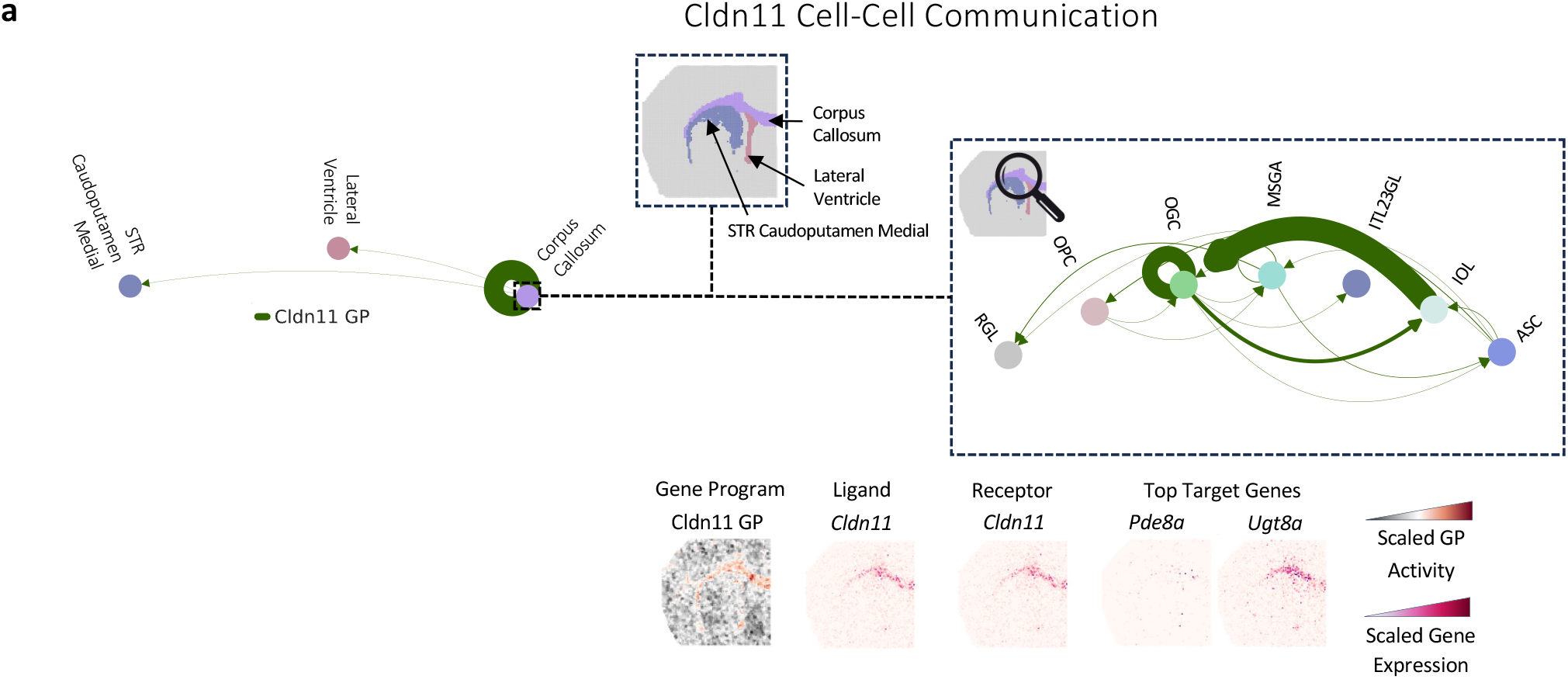
Cldn11 cell-cell communication inference. **a,** A detailed analysis of the Cldn11 GP enriched in the Corpus Callosum niche. The first circle plot displays inferred niche-to-niche communication strengths. Interactions within the Corpus Callosum niche dominate the GP with fewer interactions between the Corpus Callosum niche and neighboring niches Lateral Ventricle and STR Caudoputamen Medial. The GP activity and expression of ligand, receptor, and top target genes substantiate this analysis. Zooming into interactions involving the Corpus Callosum niche highlights communicating cell types. RGL: radial glia-like cells; OPC: oligodendrocytes precursor cells; OGC: myelin-forming oligodendrocytes; MSGA: inhibitory neurons, medial septal; ITL23GL: excitatory neurons, cortex L2/3; IOL: newly formed oligodendrocytes; ASC: astrocytes, grey matter.

**Supplementary Fig. 33.**
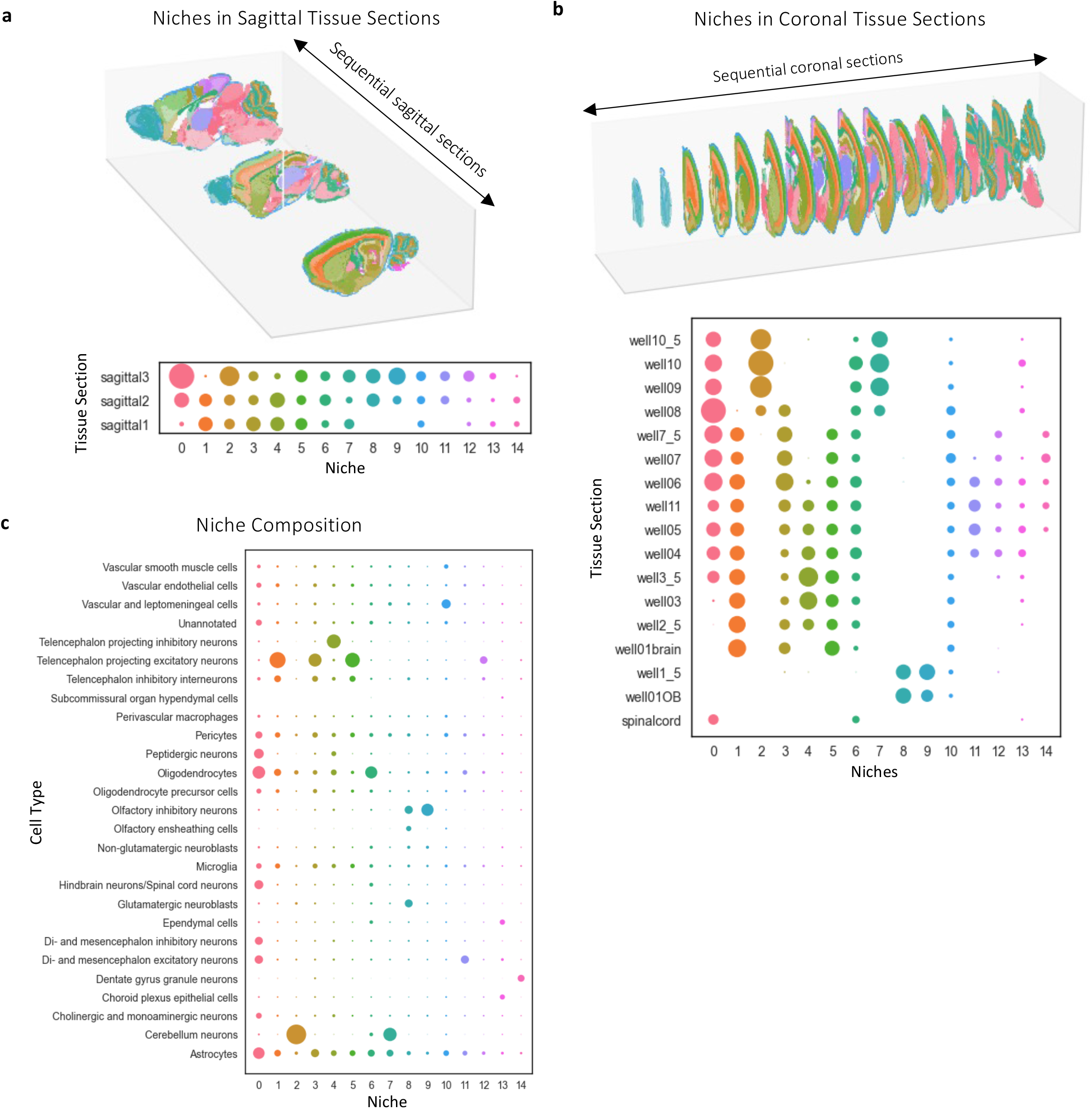
Niches identified in the mouse brain are consistent across tissue sections. **a,** Sagittal tissue sections ordered by 3D position and colored by the identified niches, showing consistency across sequential tissue sections. Below it the number of cells occurring in each tissue section for each niche. **b,** Same as **a** but for the coronal tissue sections (the spinal cord is not shown). Cell numbers are scaled separately for coronal and sagittal tissue sections. **c,** Number of cells of different cell types in each niche. 10,683 of 1,091,280 cells are not assigned to a niche and are not shown.

**Supplementary Fig. 34.**
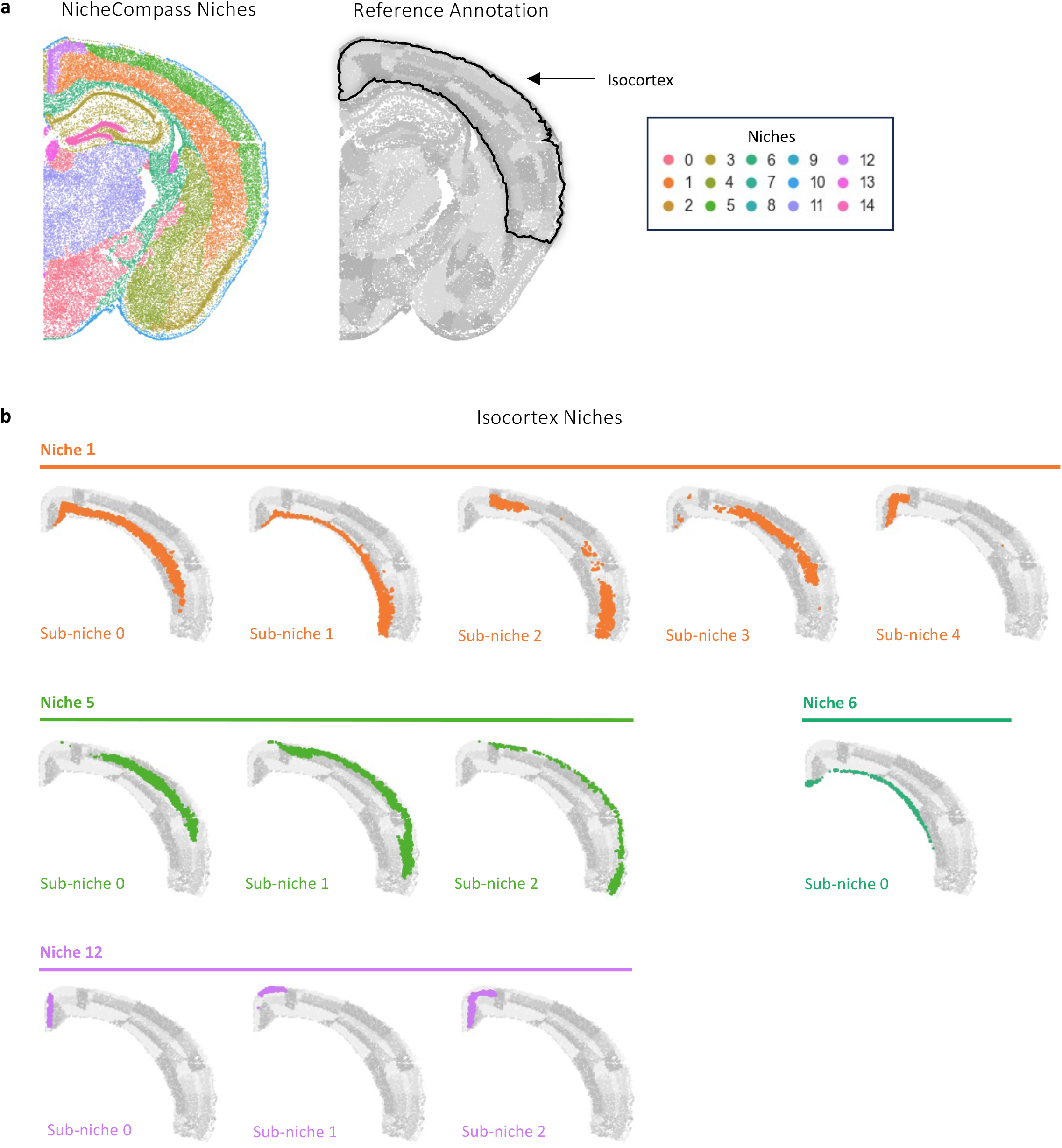
Niches identified in the mouse brain correspond to regions from a reference atlas. **a,** Coronal section showing NicheCompass niches obtained through clustering of the latent GP space (left) and regions from the Allen Mouse Brain Reference Atlas (right). The isocortex is highlighted. **b,** Magnified view showing cells assigned to the isocortex, based on the Allen Mouse Brain Reference Atlas annotations. Sub-niches with more than 250 cells annotated in this tissue section are shown. Sub-niches are obtained through clustering of cells in a niche and correspond with regions in the reference annotation.

**Supplementary Fig. 35.**
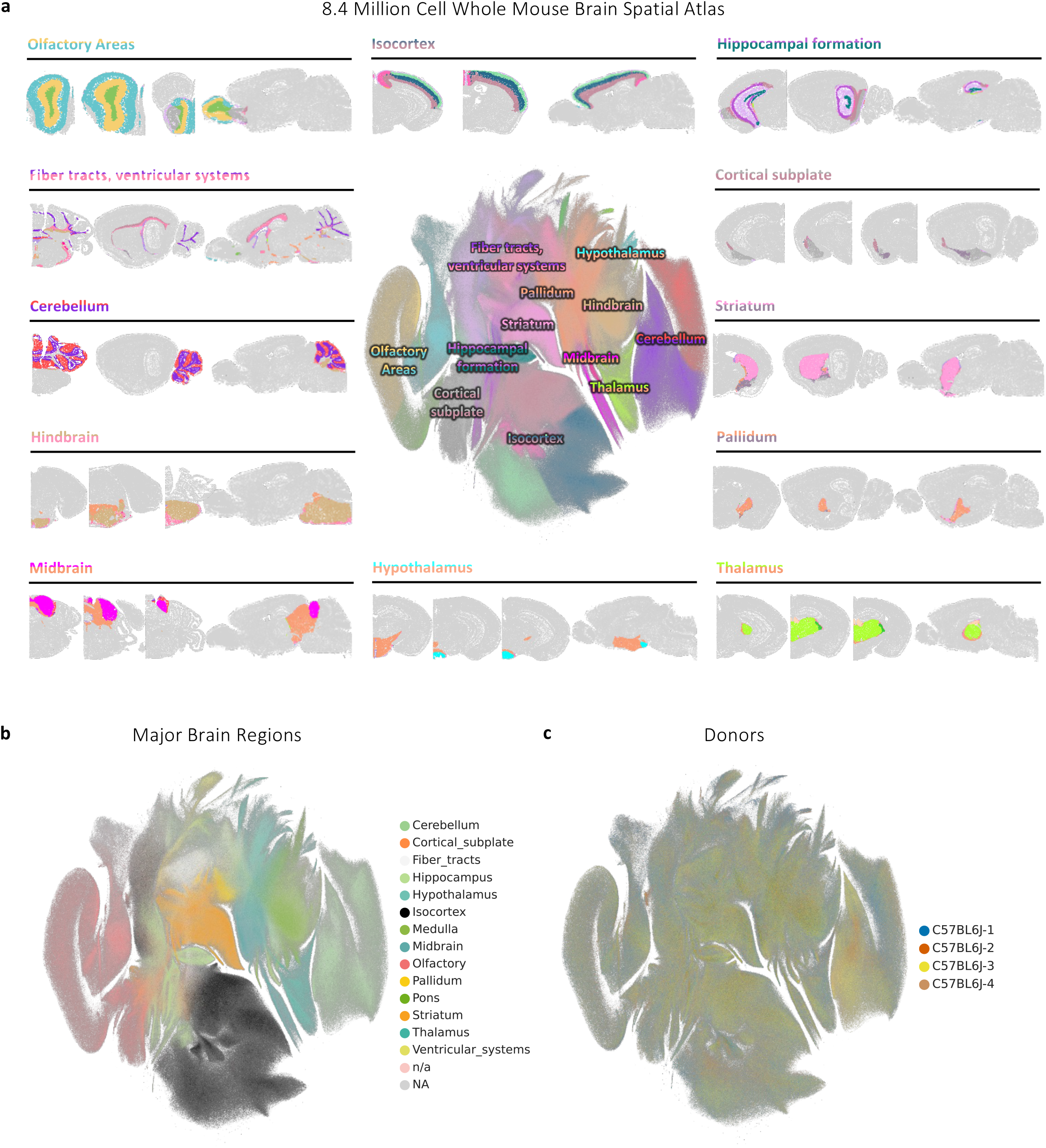
NicheCompass integrates 8.4 million cells across 239 tissue sections of a whole mouse brain spatial atlas. **a,** UMAP representation of the NicheCompass (Light) latent GP space, colored by identified niches. Around it, randomly selected tissue slices for each major brain region, also colored by identified niches. Only cells belonging to the specific region are shown. **b,c,** UMAP representations colored by major brain regions (**b**) and donor mouse (**c**), showing successful integration of cells in matching brain regions across donors.

## Methods

### Modeling Framework

#### Dataset

We define a spatial omics dataset as 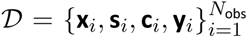, where *N*_obs_ is the total number of observations (either cells or spots), 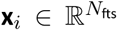 is the (self) omics feature vector of observation *i* with *N*_fts_ features, **s***_i_* ∈ ℝ^2^ is the 2D spatial coordinate vector of observation 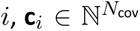 is the label-encoded covariates vector of observation *i* with *N*_cov_ categorical covariates (e.g. sample or field of view), and 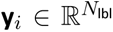 is the label vector of observation *i* with *N*_lbl_ labels (all vectors are defined as row vectors). In the unimodal scenario, **x***_i_* comprises raw gene expression counts, where raw implies the absence of normalization and scaling, and we define 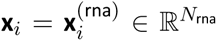, where *N*_rna_ is the number of genes. In the multimodal scenario, **x***_i_* is a concatenated vector of raw gene expression counts and raw chromatin accessibility peak counts, such that 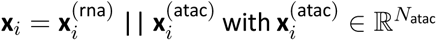, where *N*_atac_ is the number of peak regions. For all vectors, we define corresponding matrices across all observations with a bold uppercase letter, e.g. 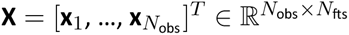.

#### Neighborhood graph

We model the spatial nature of 𝒟 with a neighborhood graph 𝒢 = (𝒱*, ℇ,* **X**, **Y**), where each node *v_i_* ∈ 𝒱 corresponds to an observation, an edge (*v_i_, v_j_*) ∈ *ℇ* denotes that observation *i* and observation *j* are spatial neighbors, **x***_i_* is the node attribute vector of node *v_i_*, and **y***_i_* is the node label vector of node *vi*. In our analyses and benchmarking experiments, 𝒢 is a disconnected graph, composed of sample-specific symmetric k-nearest neighbor subgraphs 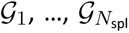, where *N*_spl_ is the number of samples. Neighbors are selected based on the pairwise Euclidean distances between the spatial coordinate vectors **s***_i_* and **s***_j_* of all nodes *v_i_, v_j_* ∈ 𝒱*_k_* that belong to the same sample *k*, with 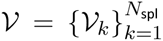. This means that (*v_i_, v_j_*) ∈ *ℇ* if observation *i* is among the k-nearest neighbors of observation *j* or vice versa, where only observations are considered for which *k_i_* = *k_j_*. We further derive a spatial adjacency matrix 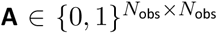 from 𝒢, where A*_i,j_* = 1 if (*v_i_, v_j_*) ∈ *ℇ* and A*_i,j_* = 0 otherwise. By defining 𝒢 as a symmetric neighborhood graph, we follow previous work ^1^ to have a flexible neighborhood-radius threshold around observations, which accounts for observation density changes in the tissue; however, other approaches can be employed, e.g. 𝒢 can be defined with a fixed neighborhood-radius threshold if different numbers of neighboring observations based on the local observation density are desired. Equally, a weighted version of 𝒢, where 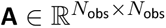, can be used to reflect differences in Euclidean distances between neighboring observations.

#### Node labels

Given 𝒢 and **X**, we first define a neighborhood omics feature vector for each observation *i*:

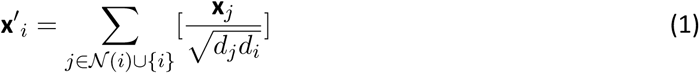

where *d_i_* is the node degree of node *i* and includes an inserted self-loop 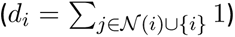. As per equation (1), **x**′*_i_* is computed for each node *i* by aggregating the omics feature vector **x***_i_* of node *i* and the omics feature vectors **x***_j_* across all neighboring nodes *j* ∈ *N*(*i*) of node *i* in 𝒢. We include node *i* to model autocrine signaling and neighboring nodes to model juxtacrine and paracrine signaling. The aggregation is based on a graph convolution norm operator ^2^, which is similar to a standard graph convolutional network layer with the important difference that it omits the linear projection with learnable weights applied to the norm-aggregated feature vector. The node label vectors are then defined as **y***_i_* = **x***_i_* || **x***^′^_i_* (concatenation of the self and neighborhood omics feature vectors).

#### Covariates

We use the label-encoded covariates vector **c***_i_* of observation *i* to explicitly model confounding factors and mitigate potential batch effects. If 𝒟 contains multiple samples, we utilize the sample *k* as the first covariate so that C*_i,_*_1_ = *k_i_*. If available, we additionally include the field of view and donor as covariates to model hierarchical batch effects. We further introduce a notation of covariates as one-hot-encoded vectors with each covariate *l* = 1*, …, N*_cov_ represented by a separate vector 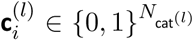, where 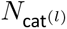 is the number of unique categories of covariate *l*. Due to the construction of 𝒢 as a disconnected graph composed of sample-specific subgraphs, one important consideration is that realized values of covariates can be directly tied to the connectivity of 𝒢 such that all observations *i* belonging to a connected component of 𝒢 will have the same realized value (e.g. sample or donor); other covariates, however, may realize different values within one connected component of 𝒢 (e.g. field of view). We denote the set of the former type of covariates *l* with *L_p_* (pure) and the set of the latter with *L_m_* (mixed).

#### Spatial prior and *de novo* gene programs (GPs)

To represent spatial prior GPs, we define two binary GP gene matrices 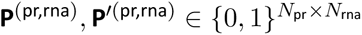, where *N*_pr_ is the number of known interaction pathways represented as prior GPs. Elements in **P**^(pr,rna)^ are assigned values of 1 for genes that are in the self-component of prior GPs, elements in **P***^′^*^(pr,rna)^ for genes that are in the neighborhood-component. In the multimodal scenario, we additionally define two binary GP peak matrices 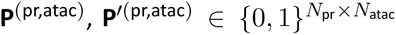. Elements in **P**^(pr,atac)^ are assigned values of 1 for peaks associated to genes in the self-component of prior GPs, elements in **P***^′^*^(pr,atac)^ for peaks associated to genes in the neighborhood-component. **P**^(pr,rna)^ and **P***^′^*^(pr,rna)^ must be provided to NicheCompass either through the in-built database APIs or as custom inputs by the user. **P**^(pr,atac)^ and **P***^′^*^(pr,atac)^ can equally be supplied by the user to represent specific gene regulatory networks; however, by default, these are determined based on the GP gene matrices, where peaks in the GP peak matrices are linked to genes in the GP gene matrices if they overlap in the gene body or proximal promotor regions (the standard configuration is up to 2000 base pairs upstream of the transcription start site).

To represent spatial *de novo* GPs, we analogously define 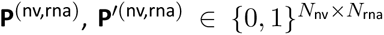, and, in the multimodal scenario, 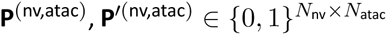, where *N*_nv_ is the number of *de novo* GPs, specified by the user (the standard configuration is *N*_nv_ = 100). Elements in **P**^(nv,rna)^ and **P***^′^*^(nv,rna)^ are assigned values of 1 for all genes that are not included in the respective component (self or neighborhood) in any prior GP. Elements in the GP peak matrices are assigned values of 1 for peaks that are linked to genes. We define *N*_gp_ = *N*_pr_ + *N*_nv_ as the total number of spatial GPs.

#### Default prior gene programs

We provide a set of default prior GPs via in-built APIs in the NicheCompass Python package. These are retrieved from four popular interaction databases, one for each GP category. For cell-cell-communication GPs, we use OmniPath ^3^ to retrieve protein-mediated interactions of ligands and receptors and MEBO-COST ^4^ to retrieve metabolite-mediated interactions of metabolites and sensors, including metabolite enzymes that serve as proxy for metabolite presence. For transcriptional regulation GPs, we retrieve transcription factors and their downstream target genes from CollecTRI ^5^,a curated collection of 12 re-sources (we access CollecTRI via the decoupler ^6^ Python package). For combined interaction GPs, we use NicheNet’s regulatory potential matrix ^7^, consisting of ligands, receptors and potential downstream target genes. We use the latest released version (V2) and follow the recommendation from MultiNicheNet ^8^ to filter the regulatory potential matrix so that each GP has a maximum of 250 target genes, ranked by their regulatory score. All prior GPs are compared, subsets are filtered, and GPs are combined if they have at least 90% overlapping source and target genes with any other GP. This results in a total of 2,925 (2,904) default mouse (human) prior GPs; thereof, 548 (490) are ligand-receptor GPs, 114 (116) are metabolite-sensor GPs, 1,286 (1,225) are combined interaction GPs, and 977 (1,073) are transcriptional regulation GPs. We use the first three GP categories across our analyses and benchmarking experiments. In the multimodal scenario, we additionally use transcriptional regulation GPs.

### NicheCompass Model

#### Model overview

NicheCompass is inspired by the variational graph autoencoder framework ^9^ but incorporates several adaptations to enable interpretable, scalable, and integrative generative modeling of spatial multiomics data. It consists of a graph encoder module and a multi-module decoder. We employ these in conjunction with a self-supervised, multi-task learning setup composed of both node- and edge-level tasks, aiming to generate spatially and molecularly consistent latent vectors 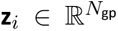 for each observation *i*. The multi-module decoder consists of a graph decoder module to reconstruct **A** from **Z**, and two omics decoder modules per omics modality. The first omics decoder module reconstructs the modality-specific (self) omics features **X**^(mod)^ (self omics decoder), and the second omics decoder module predicts the modality-specific neighborhood omics features **X***^′^*^(mod)^ (neighborhood omics decoder). This combination of decoder modules encourages the latent representations **Z** to retain spatial information from 𝒢, while incorporating molecular information from **X** and **X**′.

We use the GP matrices to mask the reconstruction of **X** and **X***^′^*, thus making the learned latent representations **Z** interpretable, with each latent feature *u* in **Z**_:,*u*_ representing a spatial GP. A subset of the latent features represents prior GPs, defined as 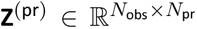, and the remaining latent features represent *de novo* GPs, defined 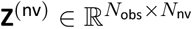, with 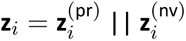.

Following the established standard for variational autoencoders, we use a standard normal distribution as prior for the latent random variables, denoted as 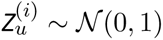, and make use of the reparameterization trick to enable end-to-end model training with backpropagation.

NicheCompass’ API interoperates with AnnData ^10^ to leverage functionalities of established single-cell and spatial analysis frameworks, such as scanpy ^11^ and squidpy ^12^, for up- and downstream workflows.

#### Encoder

The first layer of the graph encoder module consists of a fully connected layer of hidden size *N*_hid_, where we set *N*_hid_ = *N*_gp_. The purpose of this layer is two-fold: First, it enables the model to learn relevant internal cell/spot representations from the full omics feature vector **x***_i_* before considering the microenvironment (neighborhood) through message passing layers. Second, in scenarios where *N*_fts_ > *N*_gp_, this layer serves to reduce the dimensionality of **x***_i_* for lower memory consumption of the subsequent message passing layers. Its node-wise formulation is:

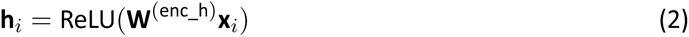

where 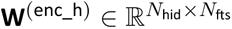 is a learnable weight matrix (including biases omitted for simplicity).

This layer is followed by two parallel message passing layers that return the mean and logstd vectors ***µ****_i_* and *log*(σ*_i_*) of the variational posterior, respectively. In our default model, we use graph attention layers with dynamic attention ^13^ and *N*_head_ = 4 attention heads that are averaged, defined node-wise as:

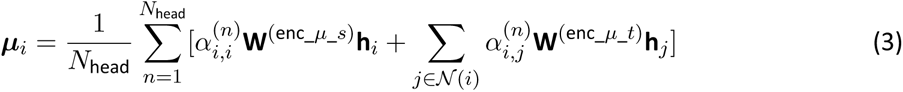

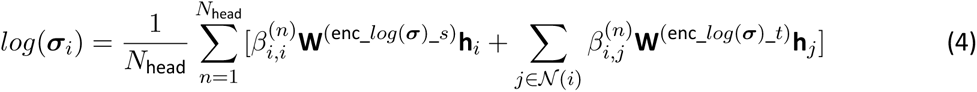

where the attention coefficients of attention head 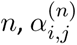 and 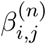, are computed as:

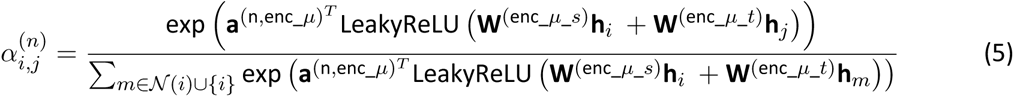

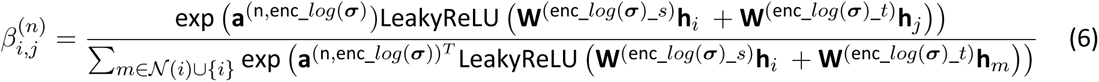

with 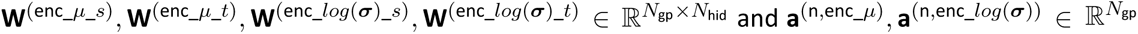 being learnable weight matrices (including biases omitted for simplicity).

In the case of NicheCompass Light, a low memory version of our model, we employ graph convolutional layers ^2^ instead of graph attention layers for message passing, so that:

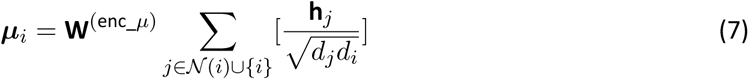

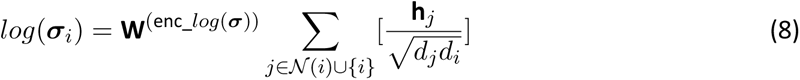

Additionally, the model learns an embedding matrix 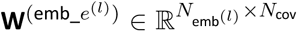 for each covariate *l* to retrieve an embedding vector 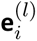 from the one-hot-encoded vector representation 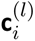, where 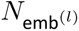 is the covariate-specific embedding size. We define 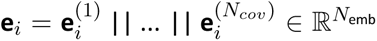 as the concatenation of embedding vectors across all covariates.

#### Decoder

The graph decoder module to reconstruct **A** computes cosine similarities of pair-wise latent feature vectors, formulated node-wise as:

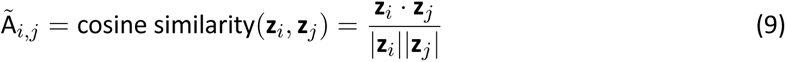

The omics decoder modules predict the node labels **Y** by estimating mean parameters 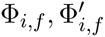 of negative binomial distributions underlying each omics feature 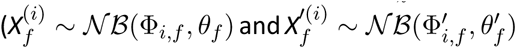, where *f* is an omics feature, *X*^(^*^i^*^)^ and *X^′^*^(^*^i^*^)^ represent omics feature random variables, and 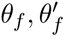 represent feature-specific inverse dispersion parameters constant across observations). They are composed of modality-specific single-layer linear decoders. To incentivize each latent feature *u* in 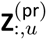 to learn the activity of a specific prior GP, the prior GP matrices mask these linear decoders. Specifically, **P**^(pr,rna)^ and, if defined, **P**^(pr,atac)^, are used as masks in the reconstruction of **X** (self-component of **Y**), and **P***^′^*^(pr, rna)^ and **P***^′^*^(pr,atac)^ are used as masks in the reconstruction of **X***^′^* (neighborhood-component of **Y**). Consequently, the GP matrices are constraining the predictive capacity of latent features to specific omics features in the self- and neighborhood-components. 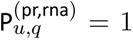 for latent feature *u* and gene *q*, the latent feature with representations **Z**_:*,u*_ is connected to gene *q* in the self decoder and can contribute to its reconstruction. Conversely, if 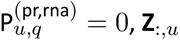 cannot contribute to the reconstruction of gene *q*. Similar statements hold true for 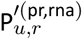 for the prediction of gene *r* in the neighborhood-component, 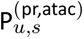 for the reconstruction of peak *s* in the self-component, and 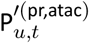 for the prediction of peak *t* in the neighborhood-component. Z*_i,u_* can therefore be interpreted as observation *i*′s representation of GP *u* where the self-component of *u* is composed of all genes *q* and peaks *s* for which 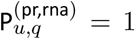 and 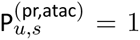, and its neighborhood-component of all genes *r* and peaks *t* for which 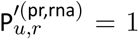 and 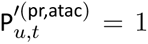. An analogous masking approach is then employed to incentivize each latent feature *u* in 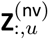 to learn the activity of a *de novo* GP, using **P**^(nv,rna)^, **P**^(nv,atac)^, **P***^′^*^(nv,rna)^ and **P***^′^*^(nv,atac)^ as GP masks. Consequently, each *de novo* GP can, in principle, reconstruct the same set of omics features that are not included in prior knowledge. Moreover, to remove batch effects from **Z**, we inject the covariates embedding vector **e***_i_* for each observation *i* into the omics decoders, which are hence formulated on observation-level as:

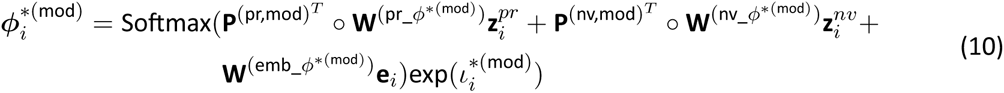

where the “*” symbol indicates either the self- or neighborhood component, mod is a placeholder for the different modalities (rna and atac), and 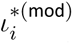 is the empirical log library size of observation *i* for modality mod. 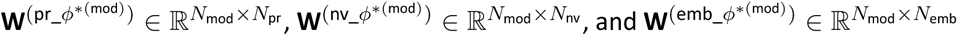 are learnable weight matrices. The Softmax activation operates across all features of an omics decoder module, constraining it to output mean proportions. The multiplication with the empirical library size ensures same size factors as in the input domain.

#### Neighbor sampling data loaders

To enable NicheCompass to scale to large datasets, which give rise to big neighborhood graphs 𝒢, and to reduce its computational footprint, we use mini-batch training with inductive neighbor sampling graph data loaders ^14^. This is in contrast to existing methods ^1,15^ that load the entire graph into memory, which is prohibitive for datasets with hundreds of thousands of observations or for the construction of spatial atlases of entire organs. Specifically, for each node *v_i_* ∈ 𝒱, we only utilize *n_spl_* = 4 sampled neighbors out of the k-nearest neighbors in 𝒢 for message passing during model training.

Due to the nature of our multi-task architecture, we employ two separate, task-centric data loaders, one for node-level tasks (node-level loader) and one for edge-level tasks (edge-level loader). One iteration of the model includes one forward pass per data loader and a joint backward pass for simultaneous gradient computation. The node-level loader is used for the omics prediction tasks including the reconstruction of **X** and the prediction of **X***^′^*. The edge-level loader is used to reconstruct **A**. To load a batch from the node-level loader, we retrieve 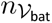 nodes from 𝒱 in random order, shuffled at each iteration, denoted with 𝒱_bat_. To load a batch from the edge-level loader, we retrieve 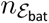 positive node pairs (*i, j*) ∈ *ℇ* from **A**, for which A*_i,j_* = 1, also in random order shuffled per iteration, where 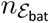 is the edge batch size. We then randomly sample 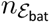 negative node pairs (*i, j*) from **A** for which A*_i,j_* = 0. We denote the corresponding combined batch of positive and sampled negative node pairs as *ℇ*_bat_. However, sampled negative node pairs in *ℇ*_bat_ might have different values for pure covariates *l* ∈ *L_p_*, i.e. 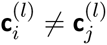. It is undesirable to include such node pairs as negative examples since there was no chance for an edge to exist. Therefore, we only keep node pairs from the initially sampled negative pairs for which both nodes have the same values for pure categorical covariates, such that the final set of node pairs used for the edge reconstruction task is 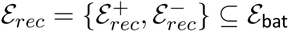 where 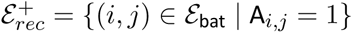 is the set of positive node pairs, and 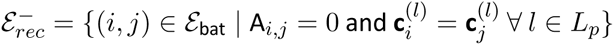 is the set of negative node pairs.

#### Gene program pruning

Not all prior GPs provided to the model may be pertinent for a particular dataset. In addition, overlapping genes between GPs can lead to weaker correlations between the learned GP representations **Z** and GP member gene and peak counts, as the variance of omics features is shared across multiple latent features. Consequently, we introduce a dropout mechanism for GP pruning during model training for the prioritization of relevant GPs and deactivation of irrelevant ones. This dropout mechanism permanently drops a subset of GPs after a certain number of warm-up epochs of model training. After the threshold of warm-up epochs has been reached, pruning is done based on the contribution of GPs to the reconstruction of **X**^(rna)^ and **X***^′^*^(rna)^ in each epoch. Specifically, we first aggregate absolute values of the learned gene expression decoder weights on a GP level (across all self- and neighborhood-component genes). Subsequently, we normalize the aggregated values with estimates of the mean absolute GP representations across all observations to obtain a GP contribution *b_u_* for each GP *u* (since the contribution of GP *u* to the reconstruction of **X**^(rna)^ and **X***^′^*^(rna)^ is the product between its latent representations **Z**_:*,u*_ and the respective gene expression decoder weights). Similar to batch normalization layers ^16^, we use an exponential moving average to compute these estimates with our batch-wise forward passes. The highest contribution across all GPs, *δ_max_*, is used as a reference value. All GP contributions are then compared with this reference and an active GP threshold ratio τ is provided as a hyperparameter. If a GP’s contribution is below τ *\* δ_max_* in any epoch after the warm-up period, the GP is dropped. For the aggregation, we use both a sum and a non-zero mean operator (two rounds of aggregation). The first round will avoid that GPs with multiple unimportant and few important genes are dropped, which would lead to a low non-zero mean-based contribution *δ_u_*. The second round will prevent a strong prioritization of GPs with many genes (high sum-based contribution *δ_u_*), which would, for example, lead to the dropout of all ligand-receptor GPs (since they have very few genes). We perform GP pruning separately for prior and *de novo* GPs, i.e. separate *δ_max_* will be computed.

In our initial experiments, we further tested group lasso regularization ^17^, with GPs representing groups, as an additional pruning mechanism; however, this resulted in less robust outcomes.

#### Gene program regularization

As many prior GPs are generic and can contain hundreds of genes, we implement a regularization mechanism to prioritize important genes within GPs during model training. To account for different functional importances of genes within GPs (e.g. a receptor is more critical than a potential downstream target gene), we first categorize genes within each GP into different categories (ligand, enzyme, tf, receptor, sensor, and target gene). We then employ a category-specific L1 regularization loss that is only applied to the gene expression decoder weights of genes belonging to the specified categories. We call this form of regularization selective regularization. In our analyses, we apply selective regularization to target genes; however, the user can specify arbitrary categories.

Similarly, *de novo* GPs can contain hundreds to thousands of genes. To encourage the model to learn GPs with high specificity, we apply a separate L1 regularization (with a different weight) to all genes contained in *de novo* GPs. For both prior and *de novo* GPs, if gene expression decoder weights are regularized to 0 during model training, corresponding chromatin accessibility decoder weights are equally set to 0, resulting in the corresponding peaks being turned off in that GP.

##### Loss function

In the unimodal scenario, our loss function consists of four core components: a binary cross-entropy loss for the reconstruction of edges in **A**, a negative binomial loss for reconstruction of the self-component **X**^(rna)^ of the node label, i.e. the nodes’ gene expression counts, a negative binomial loss for prediction of the neighborhood-component **X***^′^*^(rna)^ of the node label, i.e. the aggregated gene expression counts of node neighborhoods, and the Kullback-Leibler divergence between the variational posteriors and the standard normally distributed priors of the latent variables. In the multimodal scenario, in which case the node label is a concatenation of gene expression and peak counts, additional negative binomial losses are employed for reconstructing the self-component peak counts **X**^(atac)^ and predicting the neighborhood-component peak counts **X***^′^*^(atac)^.

The mini-batch wise formulation of the binary cross-entropy loss for edge reconstruction is:

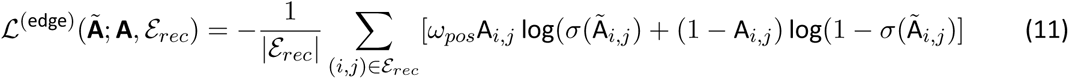

where **Ã** is the edge reconstruction logits matrix, and the edge reconstruction logit between node *i* and node *j* is computed with the cosine similarity graph decoder module. This loss component encourages nodes that are connected via an edge in the input domain to be embedded close together in the latent feature space. Since 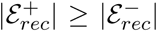 due to the removal of negative edge node pairs that do not fulfill 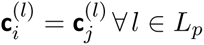, positive edge node pairs are weighted with 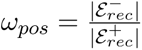 to make their contribution to *L*^(edge)^ balanced with negative edge node pairs.

The mini-batch-wise formulation of the modality-specific negative binomial losses for node label prediction is:

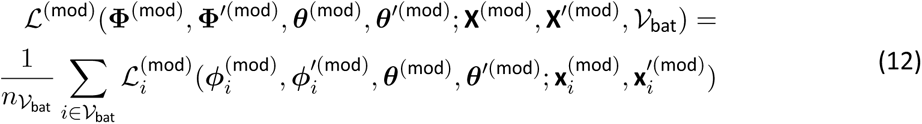

with the observation-level loss consisting of the self- and neighborhood-component negative binomial losses (NBL):

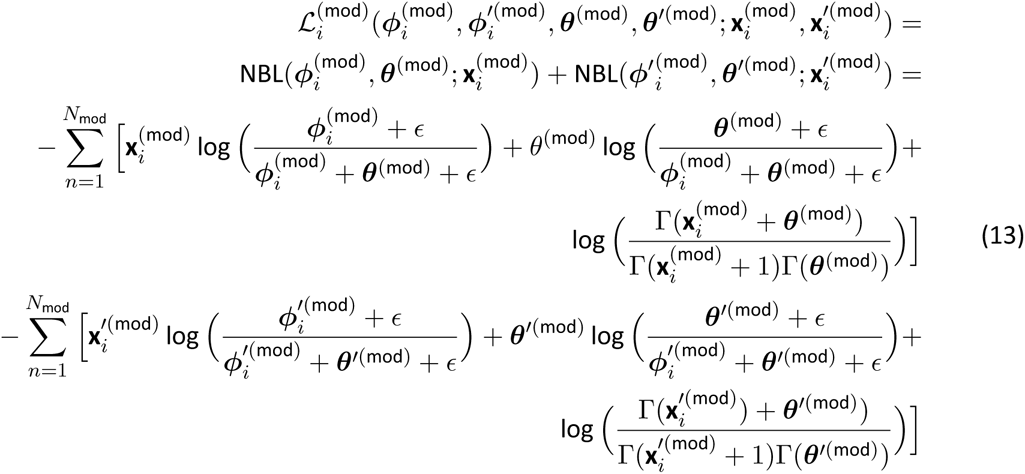

where mod is a placeholder for the modality (rna or atac), *θ*^*(mod)^ are omics feature-specific learned inverse dispersion parameters, 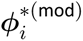 are the estimated means of the negative binomial distributions, retrieved as output of the omics decoders, and *ɛ* is a numerical stability constant.

The L1 regularization losses are defined as:

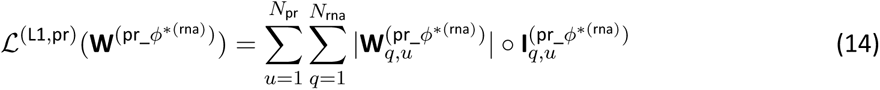

and

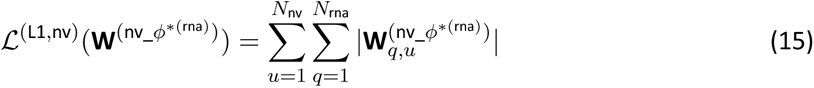

where 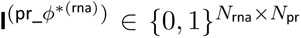 is an indicator matrix for selective regularization of prior GPs, with values of 1 for all genes that are in categories to which the L1 regularization loss should be applied to and 0 otherwise.

The mini-batch-wise formulation of the KL divergence consists of the node-level KL divergence, which includes nodes from the node-level batch, and the edge-level KL divergence, which includes both nodes from positive and negative edge node pairs in the edge-level batch:

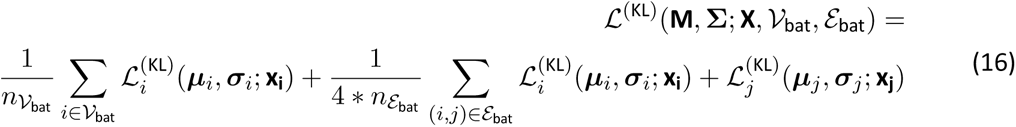

with the observation-level loss:

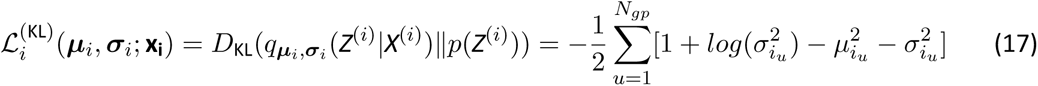

where the variational parameters ***µ****_i_* and σ*_i_* are the estimated mean and standard deviation of the approximate posterior normal distribution, and are retrieved as output of the graph encoder module.

The final mini-batch-wise loss that is optimized during model training is thus:

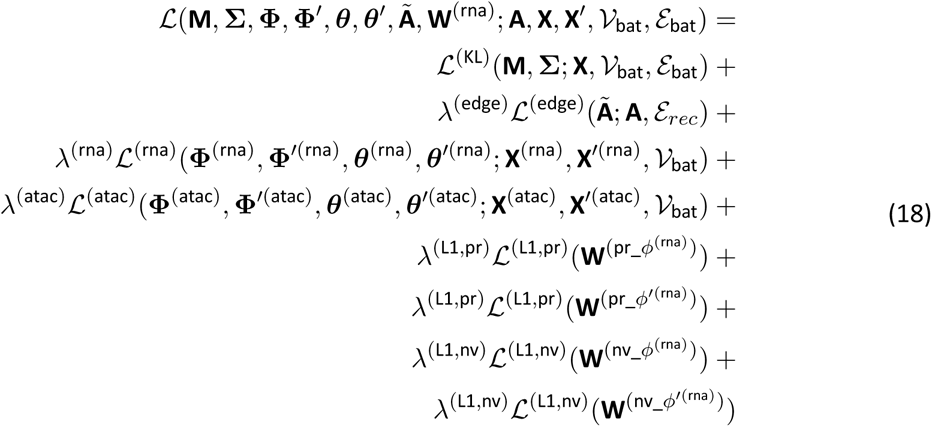

where the weighting factors of the loss components are denoted with λ.

#### Spatial reference mapping

Inspired by architectural surgery ^18^, we enable mapping of unseen query datasets onto spatial reference atlases through weight-restricted fine-tuning. A NicheCompass model is first trained on a reference atlas, containing one or multiple datasets. During subsequent training on the query dataset, all neural network weights are frozen except for weights in the covariate embedding matrices 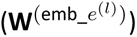. This design prevents catastrophic forgetting, while allowing the model to learn query-specific variation in the covariate embeddings. Batch effects are thus removed from query cell/spot GP representations, which can be used to contextualize the query with the reference atlas, and identify novel niches and differential latent GPs. In addition, GP pruning is active during query training through the computation of exponential moving averages of GP representations, allowing different GPs to be active in the query dataset.

### Model Evaluation Metrics

We define three metric categories: (1) spatial conservation, (2) niche coherence, and (3) batch correction. Each category comprises various individual metrics (1: CAS, CLISIS, MLAMI, GCS; 2: NASW, CNMI; 3: BLISI, PCR).

#### General specifications

We construct k-nearest neighbor graphs using the nearest_neighbors.pynndescent() functionality from the scib-metrics ^19^ package with k=15 neighbors (k=50 for CLISIS). scIB metrics are computed via the scib-metrics package with default parameters. For clustering, we employ the Leiden algorithm from scanpy ^11^, using tl.leiden(). All package versions are available at https://github.com/Lotfollahi-lab/nichecompass-reproducibility.

#### Spatial conservation metrics

Building upon previous work^19–22^, we introduce two cell type annotation-based (CAS, CLISIS) and two unsupervised (MLAMI, GCS) metrics to evaluate spatial conservation of the learned latent representations and identified cell niches at global and local scale. All four metrics are applicable in both single sample and sample integration scenarios.

CAS. The Cell Type Affinity Similarity (CAS) quantifies the conservation of spatial cell type organization in the latent space of a model compared to the physical (tissue) space. Specifically, it measures how well cell-cell contact maps ^20^ (cell type enrichment scores in a cell’s neighborhood measured across all cell types) are preserved. The CAS is scaled between 0 and 1, with higher values indicating superior cell-cell contact map similarity and hence better preservation of global spatial cell type organization. To compute the CAS, we first construct a k-nearest neighbor graph based on the physical space and count the number of edges for all pairs of cell types. The cell types are then randomly permuted across cells by sampling without replacement (1000 times), and the edges for all pairs of newly assigned cell types are counted again. The physical cell-cell contact map 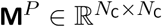, where *N*_C_ is the number of cell types, is constructed by measuring the enrichment of connections with the original cell types compared to the random permutations. In detail, the cell-cell contact map consists of cell type pair z-scores that quantify cell type affinity and are computed by first subtracting the mean edge count across permutations from the originally observed edge counts, followed by division by the standard deviation across permutations (we use squidpy’s ^12^ pl.nhood_enrichment() functionality). High values in **M***^P^* thus indicate cell type pair enrichment in the physical space compared to a random allocation. The same procedure is repeated with a k-nearest neighbor graph constructed based on the model’s latent space to create the latent cell-cell contact map 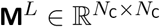. We then compute 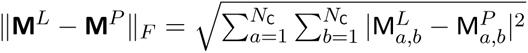 to quantify the matrix distance between **M***^L^* and **M***^P^*. Finally, we subtract this distance from 1 and perform scaling to obtain the CAS. In the case of sample integration scenarios, we obtain a joint k-nearest neighbor graph by combining the sample-wise k-nearest neighbor graphs as isolated components, while computing the latent k-nearest neighbor graph based on the integrated latent space.

MLAMI. The Maximum Leiden Adjusted Mutual Info (MLAMI) is an unsupervised metric to evaluate the degree of overlap between clusters of cells in the latent space of a model and clusters of cells in physical space. The MLAMI quantifies the preservation of global spatial organization and ranges from 0 to 1, with higher values indicating a more accurate preservation of global spatial organization. To compute the MLAMI, we construct k-nearest neighbor graphs based on the physical and latent spaces, and use them to perform three distinct clusterings, respectively, varying the cluster resolutions between 0.1 and 1.0 at increments of 0.45. To assess the similarity between the obtained clusters, we calculate the Adjusted Mutual Info (AMI) of all clustering pairs (the AMI ensures normalization against chance and compensates for mutual information to be higher for two clusterings with a larger number of clusters, even if they possess the same amount of mutual information as two clusterings with fewer clusters). We report the maximum AMI score across all clustering pairs as MLAMI. In sample integration scenarios, we obtain physical clusters for each sample separately, while computing joint clusters on the integrated latent space. In this case, the MLAMI is computed for each individual sample between the joint latent clusters, considering only cells that are part of a sample, and the sample-specific physical clusters. The mean MLAMI, computed over all samples, is then reported.

CLISIS. The Cell Type Local Inverse Simpson’s Index Similarity (CLISIS) indicates how well spatial cell type heterogeneity is preserved in the latent space of a model compared to the physical space. It is based on the Local Inverse Simpson’s Index (LISI) ^21^, and ranges between 0 and 1, with higher values indicating more accurate preservation of the local neighborhood cell type heterogeneity. To calculate the CLISIS, we first compute the cell-level Cell Type Local Inverse Simpson Indeces (CLISI) ^19^ based on the physical and latent space, respectively. The cell-level CLISI captures the degree of cell mixing in a local neighborhood around a given cell. We then divide the latent cell-level CLISIs by the physical ones and compute the logarithm to get relative local heterogeneity scores ^22^. Subsequently, we normalize these scores by the maximum possible score, compute the median of the absolute values across all cells, and subtract from 1 to arrive at the CLISIS. In the case of sample integration scenarios, we compute the latent CLISI scores on the integrated latent k-nearest neighbor graph and use separate physical k-nearest neighbor graphs per sample. The CLISIS then aggregates relative local heterogeneity scores across all samples.

GCS. The Graph Connectivity Similarity (GCS) is an unsupervised metric to compare the overlap between the graph connectivity of cells in the latent feature space of a model with that in physical space. The GCS quantifies how well local spatial organization is preserved, and it ranges between 0 and 1, with 1 indicating perfect graph connectivity similarity and 0 indicating no graph connectivity similarity. To compute the GCS, we construct k-nearest neighbor graphs based on the physical and latent space, respectively. Subsequently, we compute the Frobenius norm of the differences between the two derived adjacency matrices to measure overall dissimilarity between graph connectivity patterns. We then normalize this measure by the minimum graph connectivity overlap and perform scaling. In sample integration scenarios, the GCS is computed for each sample separately by constructing sample-specific latent and physical k-nearest neighbor graphs and we report the mean.

#### Niche coherence metrics

NASW. We define the Niche Average Silhouette Width (NASW) to measure the separability of latent clusters identified via clustering of the latent space of a model. To compute the NASW, we construct a k-nearest neighbor graph based on the latent space and perform three clusterings by varying resolutions between 0.1 and 1.0 at increments of 0.45. We then compute the Average Silhouette Width over all three clusterings and report the mean as NASW.

CNMI. We compute the Cell Type Normalized Mutual Info (CNMI) from scIB ^19^.

#### Batch correction metrics

BLISI and PCR. We compute the Batch Local Inverse Simpson’s Index (BLISI) and Principal Component Regression score (PCR) from scIB ^19^.

#### Overall score

We compute the overall score through category-balanced aggregation of scaled individual metrics. To account for natural differences in the range of individual metrics, we scale them between 0 and 1 across all compared model runs so that each individual metric has a comparable contribution to the overall score. To obtain category scores for spatial conservation (*s_s_*), niche coherence (*s_n_*), and batch correction (*s_b_*), we aggregate all scaled individual metrics from the respective category with equal weighting. To obtain the overall score *s_o_*, we aggregate all category scores with equal weighting. In single-sample scenarios, the overall score is thus computed as:

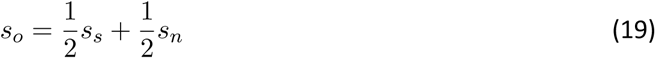

In sample integration scenarios, the overall score is computed as:

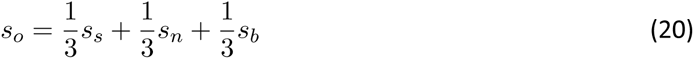

with 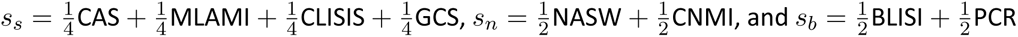.

### Benchmarking

All benchmarking experiments can be reproduced following the notebooks at https://github.com/Lotfollahi-lab/nichecompass-reproducibility. They were performed on high performance computing infrastructure with an NVIDIA A100-PCIE-40GB GPU.

#### Model configurations

NicheCompass. We trained all models with the Adam optimizer using an initial learning rate of 10^−3^,a learning rate scheduler with a patience of 4 epochs reducing the learning rate by a factor of 0.1, and early stopping with a patience of 8 epochs. We used default hyperparameters, a detailed description of which can be found at https://github.com/Lotfollahi-lab/nichecompass-reproducibility.

CellCharter. Following the tutorial at https://cellcharter.readthedocs.io/en/latest/notebooks/codex_mouse_spleen.html, we used the scVI ^23^ Python package (version 0.20.1) for dimensionality reduction and data integration and the CellCharter ^24^ Python package (version 0.2.0) with default hyperparameters to retrieve aggregated latent cell representations.

STACI. Hidden layer sizes of STACI were set to be three times the size of the input features (approximately equivalent to the configuration used in the original publication ^1^). The number of epochs was set to 1000 after which no further improvements were observed. Other hyperparameters were set according to the script at https://github.com/uhlerlab/STACI/blob/master/train_gae_starmap_multisamples.ipynb. Since there was no published Python package, the codebase was taken from GitHub (https://github.com/uhlerlab/STACI/blob/master) on 23.11.2023.

GraphST. We used default hyperparameters, following tutorial 1 at https://deepst-tutorials. readthedocs.io/en/latest for model training, using the GraphST ^15^ Python package (version 1.1.1).

DeepLinc. We used default hyperparameters for model training. Since there was no published Python package, the codebase was taken from GitHub (https://github.com/xryanglab/DeepLinc) on 22.05.2023.

#### Experiments

SlideSeqV2 mouse hippocampus. We carried out one training run for each method, using a symmetric k-nearest neighbor graph with k = 4 neighbors. We clustered the latent representations to obtain a similar number of niches for all methods. In the case of CellCharter, we used the inbuilt tl.Cluster() functionality for clustering. For all other methods, we computed clusters using the tl.leiden() functionality from scanpy ^11^.

SlideSeqV2 mouse hippocampus 25% subsample. To obtain a spatially consistent 25% subsample of the full dataset, we sampled cells from the middle of the tissue in both y-coordinate directions while retaining the full x-coordinate range. The training methodology was equivalent to the experiment on the full dataset.

NanoString CosMx human non-small cell lung cancer 10% subsample. To obtain a spatially consistent 10% subsample of the full dataset, we sampled cells according to fields of view, starting with the first field of view and stopping after the threshold of 10% had been reached. We used the same training methodology as in the SlideSeqV2 mouse hippocampus experiment. We computed separate k-nearest neighbor graphs for each sample and integrated them into a disconnected graph. In the standard NicheCompass model, we used the sample and field of view as covariates. We matched the number of identified niches between all methods and annotated them based on cell type proportions.

Single-sample and sample integration benchmarking. We performed eight training runs for each method on the full and subsampled datasets, varying the number of neighbors between 4 and 16 at increments of 4 (2 runs each). For subsampling, we included 1%, 5%, 10%, 25% and 50% of the full dataset while maintaining spatial consistency.

### Analysis

#### Data visualization

We used uniform manifold approximation and projection (UMAP) to embed cells in 2D for visualization. Specifically, we computed a k-nearest neighbor graph on the latent representations using scanpy’s ^11^ pp.neighbors() functionality with default parameters, followed by computation of the UMAP representations with tl.umap(), also with default parameters. In the case of the 8.4 million cell whole mouse brain spatial atlas, before neighborhood graph computation we first computed PCA of the latent feature representations, using tl.pca() with default parameters.

#### Hierarchical niche identification

The identification of tissue niche hierarchies is performed through a two-step process. First, Leiden clustering is employed on the latent space of a model with the scanpy ^11^ tl.leiden() functionality, where each obtained cluster represents a niche. To subcluster specific niches, multiple rounds of Leiden clustering are computed. Second, hierarchical clustering is computed with the scanpy tl.dendrogram() functionality with niche labels as input and “ward” or “single” as linkage method.

#### Gene program feature importances

We use the learned weights of the linear omics decoders to determine GP-specific gene and peak importances. Specifically, we take the absolute values of the gene expression (chromatin accessibility) decoder weights and normalize them across all genes (peaks) in the self- and neighborhood-components of a GP. The resulting gene (peak) importances thus sum up to 1 for each GP.

#### Gene program activities

The inferred latent GP representations of NicheCompass quantify how active pathways are in different cells/spots. However, NicheCompass is agnostic to the sign of these; for instance, a negative latent value with a negative learned weight in an omics decoder would lead to the same outcome as a positive latent value with a positive learned weight. For positive latent values to represent upregulation of a GP, we thus reverse the sign of latent GP representations based on the sign of the weights in the omics decoders. For prior GPs, we reverse the sign if the weights of ligand and receptor or enzyme and sensor genes are negative. For *de novo* GPs, we reverse the sign if the majority of weights is negative. We refer to the sign-corrected latent GP representations as GP activities.

##### Differential testing of gene program activities

Inspired by previous work ^25^, we perform differential testing of GP activities to determine enriched GPs in a group of interest compared to a comparison group. Specifically, we test the hypothesis 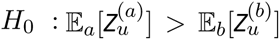 versus the alternative hypothesis 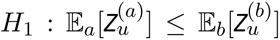, where *u* is the index of the tested GP, and *Z*^(^*^a^*^)^ and *Z*^(^*^b^*^)^ denote random variables for the GP activities of group of interest *a* and the comparison group *b*, respectively. Following a previously introduced approach ^23^, we use the logarithm of the Bayes Factor *K*, a Bayesian generalization of the p-value, as test statistic. The magnitude of this statistic can be interpreted as significance level and it is calculated as the ratio between the probabilities of *H*_0_ and *H*_1_:

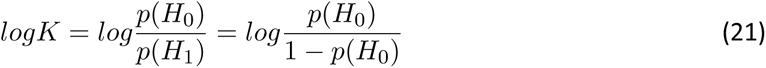

We can compute *p*(*H*_0_) as:

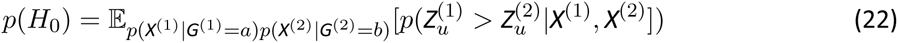

where *G*^(1)^ and *G*^(2)^ denote the independent random variables for group membership with the index corresponding to the omics features random variables *X*^(1)^ and *X*^(2)^ and

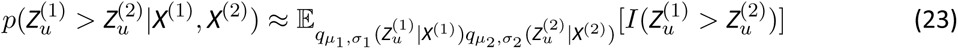

Since the approximate posterior 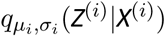 is a Gaussian distribution in our case, the expectation from equation (23) can be calculated via:

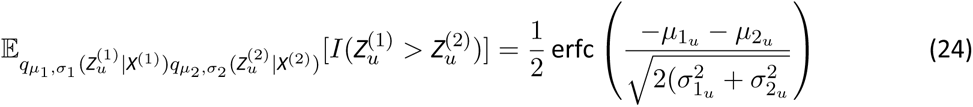

In our analyses, we consider GPs with a test statistic of *|log K|* ≥ 2.3 as differentially expressed. This threshold corresponds to strong evidence according to the interpretation scale introduced by Kass et. al ^26^ and is equivalent to a relative ratio of probabilities of exp(2.3) ≈ 10.

#### Selection of characterizing gene programs for each niche

To obtain characterizing GPs (also used for niche annotation), we first compute the set of enriched GPs for each niche with a one-vs-rest differential log Bayes Factor test. From the set of enriched GPs, we then select two characteristic GPs per niche based on correlation between GP activities and the expression of the GP’s important target genes and ligand and receptor or enzyme and sensor genes.

#### Gene program communication potential scores

GP activities reflect how strongly present pathways are within cells/spots, taking into account their microenvironment (neighborhood). As a result, high activities can be due to high counts of neighborhood-component omics features in a cell’s/spot’s neighborhood or high counts of self-component omics features in a cell/node itself, where the highest activity is expected if both coincide. While these activities are useful in terms of quantifying the presence of spatial pathways, they are not directly suited for the inference of cell-cell communication events between specific interaction partners. To address this, we propose source and target communication potential scores, representing the source- and target-specific pathway activities of cells/spots. Communication potential scores build on inferred GP activities of cell-cell communication or combined interaction GPs and serve as robust intermediate measures for the inference of cell-cell communication events.

To compute source and target communication potential scores, we first scale the gene expression counts for each gene between 0 and 1 to avoid bias towards highly-expressed genes. For each GP, we then multiply the scaled gene expression of each member gene by its corresponding omics decoder weights learned by the model. This results in GP-specific scores for each gene in the self- and neighborhood-component of a GP. Subsequently, we compute the average across all genes in the self- and neighborhood-component, respectively. We then multiply these averages by the inferred GP activities. Multiplication with the self-component average results in the target communication potential score and multiplication with the neighborhood-component average results in the source communication potential score. Negative communication potential scores are set to 0.

#### Gene program communication strengths

GP communication strengths build on GP communication potential scores to quantify the likelihood for cell-cell communication events between specific interaction partners. To compute GP communication strengths, we first generate a GP-specific neighborhood graph to reflect the length scale of that specific GP (by default we use 𝒢). All nodes that are neighbors in this graph are potential interaction partners. We then perform pair-wise multiplications using the source and target communication potential scores of all neighboring nodes. The resulting values represent node-wise directional communication strengths that can then be aggregated, e.g. on cell- or niche-level. Once aggregated, communication strengths are normalized between 0 and 1.

### Datasets

All datasets were previously published. Preprocessing beyond the one performed in the original publications can be reproduced as described in notebooks at https://github.com/Lotfollahi-lab/nichecompass-reproducibility. All preprocessed data is available as AnnData-compatible “.h5ad” files ^10^ as described at https://github.com/Lotfollahi-lab/nichecompass-reproducibility. Cell type labels and metadata were obtained from the original publications unless stated otherwise.

#### seqFISH mouse organogenesis

Both the original and imputed versions of the seqFISH mouse organogenesis data ^20^ were retrieved from https://marionilab.cruk.cam.ac.uk/SpatialMouseAtlas/. This dataset consists of 57,536 single cells across six sagittal 8-12 somite stage mouse embryo tissue sections, obtained from three different embryos with the following composition: 19,451 cells (embryo 1), 14,891 cells (embryo 2), and 23,194 cells (embryo 3). The authors probed 351 barcoded genes in the original dataset and performed imputation based on a single-cell gastrulation reference atlas to generate a full transcriptome spatial map with 29,452 features. For use with NicheCompass, we converted the original Bioconductor RDS files into AnnData-compatible “.h5ad” files ^10^. We further filtered cells annotated as low quality by the original authors. Since the imputation was performed on log count level, we computed a reverse log normalization to obtain estimated raw counts as required by NicheCompass. This resulted in some genes with very high imputed raw counts; consequently, we filtered genes based on their maximum imputed raw counts per cell: genes with raw counts greater than 141 were removed since this was the maximum raw count per cell for genes in the original dataset. This resulted in 29,239 remaining features; of these, we kept the 5,000 most spatially variable ones as determined by the spatial autocorrelation score Moran’s I, computed with the gr.spatial_autocorr() functionality of squidpy ^12^. For models that were trained on multiple samples, we defined the sample as the only covariate, with each of the embryo tissue sections representing a separate sample.

#### SlideSeqV2 mouse hippocampus

The SlideSeqV2 dataset ^27^ was retrieved via the datasets.slideseqv2() functionality of squidpy ^12^. This dataset consists of a mouse hippocampus puck with 41,786 observations at near-cellular resolution and 4,000 genes. Since the dataset was preprocessed and contained log counts, we computed a reverse log normalization to obtain raw counts as required by NicheCompass.

#### MERFISH mouse liver

The MERFISH mouse liver dataset was retrieved from https://info.vizgen.com/mouse-liver-access (animal 1, replicate 1). This dataset consists of a mouse liver tissue section with 367,335 cells and 347 probed genes. We followed the preprocessing vignette from https://squidpy.readthedocs.io/en/stable/notebooks/tutorials/tutorial_vizgen_mouse_liver.html.

#### nanoString CosMx human non-small cell lung cancer

The nanoString CosMx human non-small cell lung cancer dataset ^28^ was retrieved from https://nanostring.com/products/cosmx-spatial-molecular-imager/ffpe-dataset/nsclc-ffpe-dataset/. This dataset consists of 8 tissue sections from 5 donors, with a total of 800,327 cells, distributed as follows: 93,206 cells (donor 1, replicate 1), 93,206 cells (donor 1, replicate 2), 91,691 cells (donor 1, replicate 3), 91,691 cells (donor 2), 77,391 cells (donor 3, replicate 1), 115,676 (donor 3, replicate 2), 66,489 cells (donor 4) and 76,536 cells (donor 5). The expression level of a total of 960 genes is measured. Each tissue section contains multiple field of views, ranging in numbers from 20 to 45. We filtered cells with less than 50 counts as low quality cells, leaving 766,313 remaining cells. For model training with multiple samples, we defined the sample, field of view, and donor as covariates.

#### Xenium human breast cancer

The Xenium human breast cancer dataset ^29^ was retrieved from https://www.10xgenomics.com/products/xenium-in-situ/preview-dataset-human-breast. It contains a total of 282,363 cells across two replicates: 164,000 cells (replicate 1) and 118,363 cells (replicate 2) with 313 probed genes. We filtered cells that had less than 10 counts across all genes or non-zero counts for less than 3 different genes. To annotate cell types and cell states, we used a typical scanpy ^11^ workflow, encompassing PCA to retrieve 50 principal components, neighborhood graph computation with 50 neighbors, and Leiden clustering followed by annotation based on differential expression of marker genes.

#### STARmap PLUS mouse central nervous system

The STARmap PLUS mouse central nervous system dataset ^30^ was downloaded from Zenodo https://zenodo.org/records/8327576 and processed to AnnData-compatible “.h5ad” files ^10^. Genes were selected based on expression in at least 10% of cells across all samples in the dataset. STAlign ^31^ was used to align a coronal tissue section with the Allen Mouse Brain Reference Atlas ^32^.

#### MERFISH whole mouse brain

The MERFISH whole mouse brain dataset ^33^ was downloaded from CELLxGENE (https://cellxgene.cziscience.com/collections/0cca8620-8dee-45d0-aef5-23f032a5cf09). The dataset has 1,122 genes and, after preprocessing performed by the original authors, contains a total of 8.4 million cells across 239 coronal and sagittal sections from four animals. For model training, we defined the sample and donor as covariates.

#### Spatial ATAC-RNA seq mouse brain

The Spatial ATAC-RNA seq mouse brain dataset ^34^ was retrieved from https://www.ncbi.nlm.nih.gov/geo/query/acc.cgi?acc=GSE205055 (gene expression counts and spatial coordinates) and https://brain-spatial-omics.cells.ucsc.edu/ (peak counts and cell type labels). We used the postnatal day 22 brain sample. This sample consists of 9,215 observations at spot-level resolution with 22,914 genes and 121,068 peaks. For use with NicheCompass, we converted the original Seurat RDS file containing peak counts into an AnnData-compatible “.h5ad” file ^10^. We filtered genes and peaks with counts in less than 46 cells (0.05% of all cells) and kept the 3,000 most spatially variable genes and 15,000 most spatially variable peaks, determined by the spatial autocorrelation score Moran’s I as described earlier. To enable mapping to peaks, we further dropped all genes that were not annotated in the GENCODE 25 release (https://ftp.ebi.ac.uk/pub/databases/gencode/Gencode_mouse/release_M25/gencode.vM25.chr_patch_hapl_scaff.annotation.gtf.gz), resulting in 2,785 remaining genes. Finally, in the process of mapping peaks to genes, we dropped all peaks that did not overlap with any gene body or promoter region in the resulting gene set, leaving 3,337 remaining peaks.

